# Genome-wide association studies reveal novel loci associated with pyrethroid and organophosphate resistance in *Anopheles gambiae* s.l.

**DOI:** 10.1101/2023.01.13.523889

**Authors:** Eric R. Lucas, Sanjay C. Nagi, Alexander Egyir-Yawson, John Essandoh, Sam Dadzie, Joseph Chabi, Luc S. Djogbénou, Adandé A. Medjigbodo, Constant V. Edi, Guillaume K. Ketoh, Benjamin G. Koudou, Arjen E. Van’t Hof, Emily J. Rippon, Dimitra Pipini, Nicholas J. Harding, Naomi A. Dyer, Louise T. Cerdeira, Chris S. Clarkson, Dominic P. Kwiatkowski, Alistair Miles, Martin J. Donnelly, David Weetman

## Abstract

Resistance to insecticides in *Anopheles* mosquitoes threatens the effectiveness of the most widespread tools currently used to control malaria. The genetic underpinnings of resistance are still only partially understood, with much of the variance in resistance phenotype left unexplained. We performed a multi-country large scale genome-wide association study of resistance to two insecticides widely used in malaria control: deltamethrin and pirimiphos-methyl. Using a bioassay methodology designed to maximise the phenotypic difference between resistant and susceptible samples, we sequenced 969 phenotyped female *An. gambiae* and *An. coluzzii* from ten locations across four countries in West Africa (Benin, Côte d’Ivoire, Ghana and Togo), identifying single nucleotide polymorphisms (SNPs) and copy number variants (CNVs) segregating in the populations. The patterns of resistance association were highly multiallelic and variable between populations, with different genomic regions contributing to resistance, as well as different mutations within a given region. While the strongest and most consistent association with deltamethrin resistance came from the region around *Cyp6aa1*, this resistance was based on a combination of several independent CNVs in *An. coluzzii*, and on a non-CNV bearing haplotype in *An. gambiae*. Further signals involved a range of cytochrome P450, mitochondrial, and immunity genes. Similarly, for pirimiphos-methyl, while the strongest signal came from the region of *Ace1*, more widespread signals included cytochrome P450s, glutathione S-transferases, and a subunit of the *nAChR* target site of neonicotinoid insecticides. The regions around *Cyp9k1* and the *Tep* family of immune genes were associated with resistance to both insecticide classes, suggesting possible cross-resistance mechanisms. These locally-varying, multigenic and multiallelic patterns highlight the challenges involved in genomic monitoring and surveillance of resistance, and form the basis for improvement of methods used to detect and predict resistance. Based on simulations of resistance variants, we recommend that yet larger scale studies, exceeding 500 phenotyped samples per population, are required to better identify associated genomic regions.

## 1. Introduction

Vector-borne diseases such as malaria kill an estimated 700,000 people every year (Roth et al., 2018) but are vulnerable to methods that target the vectors that are essential for transmission. This has been exploited in malaria, where the primary, and most effective form of intervention, remains the control of the vector population through the use of insecticides (Bhatt et al., 2015). In the same way as interventions targeting pathogens lead to drug resistance, interventions against vectors have led to the widespread evolution of resistance to insecticides (Hancock et al., 2020). Understanding the genetic basis of this resistance is crucial for managing and informing malaria control interventions. This not only applies to current widely-used insecticides, but also to new compounds coming to market, where the opportunity exists to understand the basis of resistance at the earliest stages of deployment.

The primary malaria vectors in Sub-Saharan Africa are *Anopheles gambiae* and *An. coluzzii*, two sister species of mosquito that have largely similar genomes and well-documented capacity for hybridization and introgression (Clarkson et al., 2014; Grau-Bové et al., 2020, 2021; Weetman et al., 2012). Major mutations with large effects on resistance, often at the insecticide’s target site of action, have been discovered in these species, yet a large portion of the phenotypic variance in resistance remains unexplained (Donnelly et al., 2016). This “residual” resistance could involve a variety of detoxification mechanisms, such as increased expression or efficacy of genes that bind, metabolise or transport the insecticide. Identifying these mechanisms is challenging because the pool of genes that could be involved is much larger than that of the target site genes, and because modification of gene expression can occur in many ways (copy number variation, mutation of regulatory regions, modulation of transcription factors).

Studies of residual resistance thus require a scale, both in terms of genomic coverage and sample size, that has until recently been unfeasible. However, the falling costs of whole genome sequencing, large data centres and software pipelines capable of handling large sample sizes now make this a viable option. To this end, we set up the Genomics for African *Anopheles* Resistance Diagnostics (GAARD, https://www.anophelesgenomics.org) project, a collaborative endeavour to investigate the genomics of insecticide resistance through large scale whole genome sequencing. Here we investigate the genomic basis of resistance in West Africa to two insecticides widely used in malaria control. The first insecticide, deltamethrin, is a pyrethroid and a common constituent of insecticide-treated bednets (ITNs) (Lindsay et al., 2021). Since ITNs have now been in circulation for many years, the mosquito populations in our study have a relatively long history of exposure. ITNs remain a cornerstone of vector control (Lindsay et al., 2021), and thus resistance to them is an important consideration when planning interventions. The second insecticide is pirimiphos-methyl (PM), an organophosphate insecticide that is deployed in vector control in the form of indoor residual spraying (IRS), where interior walls of buildings are coated in insecticide to kill mosquitoes when they rest. Use of PM is more recent, and more sporadic as IRS is less widely implemented (Oxborough, 2016) making PM resistance less widespread than deltamethrin resistance (Supplementary Fig. S1). We conducted a large-scale, multi-country genome-wide association study (GWAS) of insecticide resistance, testing mosquitoes from six different populations from four different West African countries for resistance against deltamethrin and PM.

## 2. Results

### 2.1. Overview of data

We obtained sequencing data for analysis from a total of 969 individual female mosquitoes across 10 sample sets (defined as samples from a single collection location of a given species phenotyped against one of the two insecticides, Fig. 1, Table 1). The phenotype of each individual was defined by whether they were alive (“resistant”) or dead (“susceptible”) after exposure to a given dose of insecticide.

**Figure 1.**
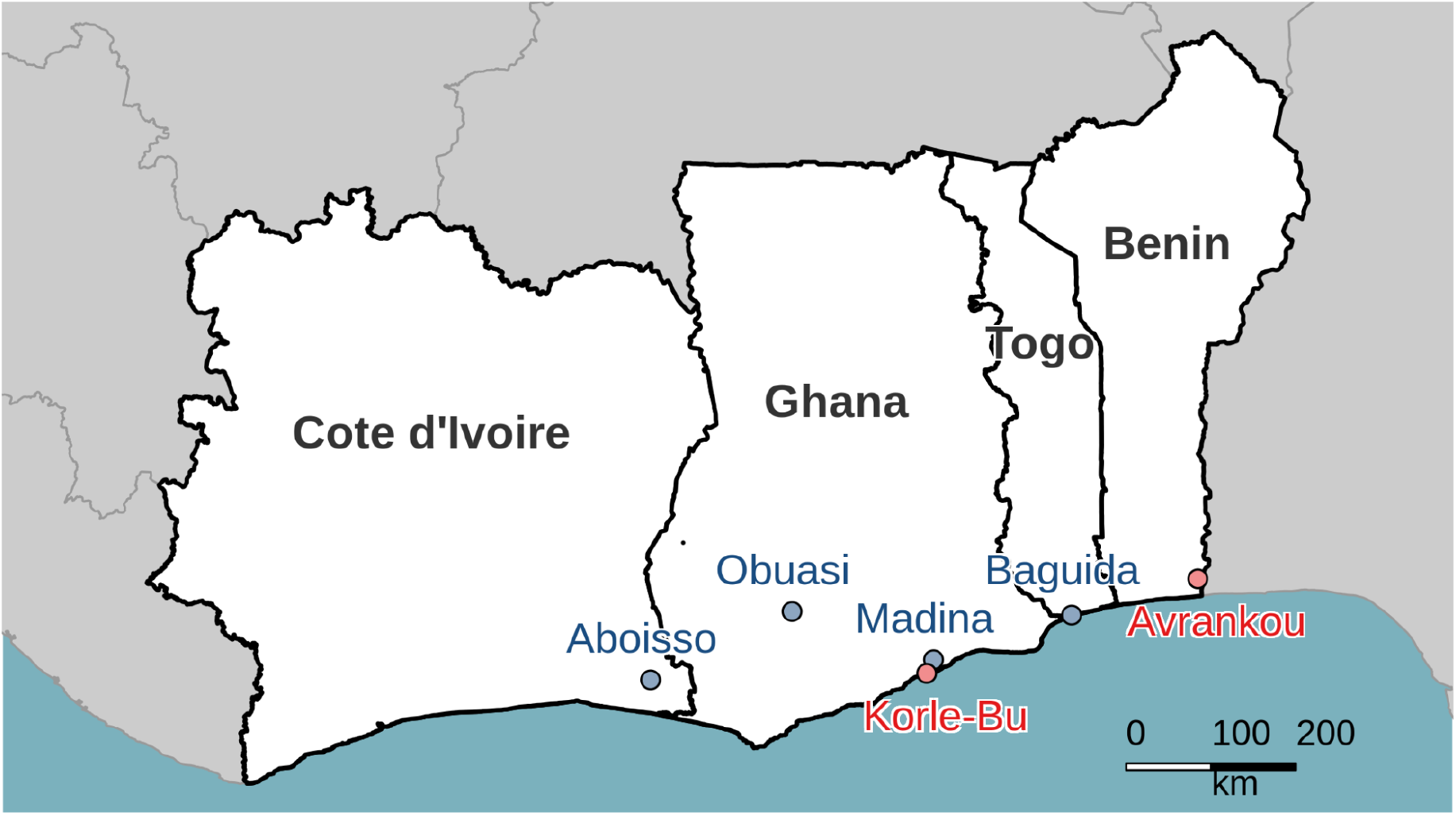
Sampling locations for the study, showing *An. gambiae* sites in blue and *An. coluzzii* sites in red.

**Table 1.**
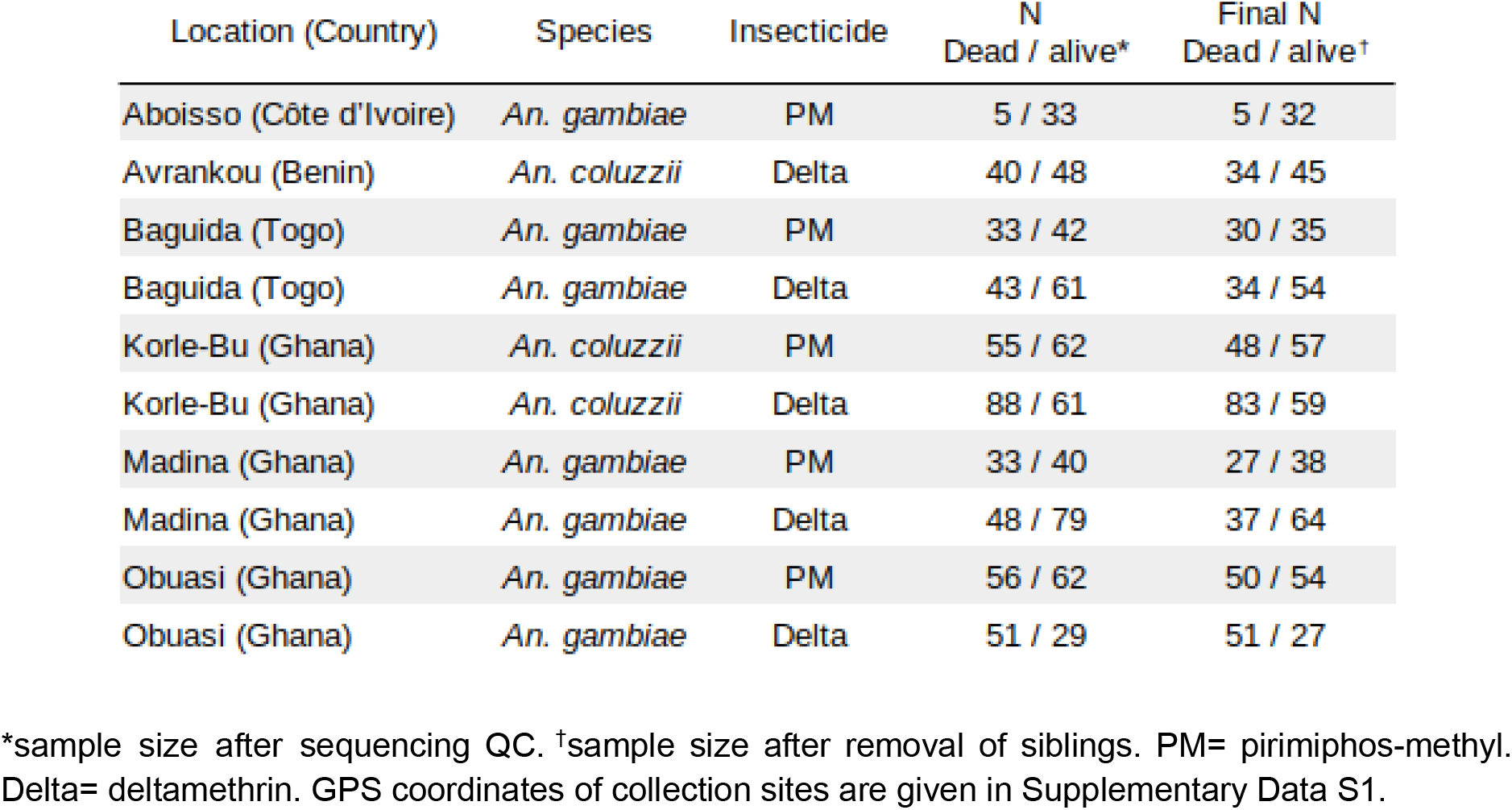
Number of samples sequenced in each of the sample sets

| Location (Country) | Species | Insecticide | N<br>Dead / alive* | Final N<br>Dead / alive† |
| --- | --- | --- | --- | --- |
| Aboisso (Côte d'Ivoire) | *An. gambiae* | PM | 5 / 33 | 5 / 32 |
| Avrankou (Benin) | *An. coluzzii* | Delta | 40 / 48 | 34 / 45 |
| Baguida (Togo) | *An. gambiae* | PM | 33 / 42 | 30 / 35 |
| Baguida (Togo) | *An. gambiae* | Delta | 43 / 61 | 34 / 54 |
| Korle-Bu (Ghana) | *An. coluzzii* | PM | 55 / 62 | 48 / 57 |
| Korle-Bu (Ghana) | *An. coluzzii* | Delta | 88 / 61 | 83 / 59 |
| Madina (Ghana) | *An. gambiae* | PM | 33 / 40 | 27 / 38 |
| Madina (Ghana) | *An. gambiae* | Delta | 48 / 79 | 37 / 64 |
| Obuasi (Ghana) | *An. gambiae* | PM | 56 / 62 | 50 / 54 |
| Obuasi (Ghana) | *An. gambiae* | Delta | 51 / 29 | 51 / 27 |

\*sample size after sequencing QC. †sample size after removal of siblings. PM= pirimiphos-methyl. Delta= deltamethrin. GPS coordinates of collection sites are given in Supplementary Data S1.

Given that mosquitoes were derived from larval collections, we investigated whether our samples included close kin pairs, which could potentially introduce population stratification. Pairwise calculations of kinship across all samples confirmed the presence of full siblings. We aggregated individuals into full sib groups by considering that two individuals that share a full sibling are also full siblings, resulting in the identification of 99 sib groups containing a total of 238 individuals (max sib group size: 9; 73% of sib groups were of size 2). Samples from different locations were never found in the same sib group. Depending on the analysis (see methods), we either discarded all but one randomly chosen individual per sib group per sample set (thus removing 105 samples) or performed permutations in which we varied which individuals were discarded in each sib group.

In *An. coluzzii* from Avrankou, *Cyp4j5-43F (P* = 0.007) and *Vgsc*-1527T *(P* = 0.02) were both associated with increased resistance to deltamethrin. *Vgsc*-402L (generated by either of two nucleotide variants: 402L(C) and 402L(T)) and *Vgsc-*1527T are completely linked, and thus *Vgsc*-1527T here refers to the haplotype carrying both SNPs. Since *Vgsc*-995F and *Vgsc-*402L/1527T are mutually exclusive, and wild-types are absent in our dataset, the significant association of *Vgsc-*402L/1527T suggests that this haplotype provides higher resistance to deltamethrin than does *Vgsc*-995F. None of the established resistance markers were associated with deltamethrin resistance in any other populations.

*Ace1*-280S was highly significantly associated with resistance to PM in all populations (*P* < 0.001 in all cases) except Baguida. In two populations, we also found PM-associations with other markers: in *An. gambiae* from Baguida, *Vgsc*-1570Y was positively associated with resistance (*P* = 0.01), which is a surprising finding for a pyrethroid target site mutation, whilst in *An. coluzzii* from Korle-Bu, *Vgsc*-995F (*P* = 0.04) and *Gste2*-114T (*P* = 0.02) were negatively associated with PM resistance.

### 2.2. CNVs

Mutations that increase the number of genomic copies of a gene, known as copy number variants (CNVs), have been shown to be commonly associated with insecticide resistance in insects, including *Anopheles* mosquitoes (Weetman et al., 2018). We identified CNVs in genes associated with metabolic resistance, using both gene copy number (using sequencing coverage to estimate the number of copies of each gene) and screening for known *An. gambiae* CNV alleles (CNVs for which the precise start and end points have been previously identified (Lucas, Miles, et al., 2019; The Anopheles Gambiae 1000 Genomes Consortium, 2021)). We found CNVs in *Cyp6aa1/2* and the *Gste* cluster at high frequency in *An. coluzzii* populations, while CNVs in *Cyp9k1* were far more common in *An. gambiae* (Fig. 3, Supplementary Figure S2). Copy number of *Cyp6aa1* was positively associated with deltamethrin resistance in *An. coluzzii*. This was significant in Korle-Bu (*P* = 0.01), with a trend in the same direction in Avrankou (*P* = 0.07). In *An. gambiae, Cyp6aa1* CNVs were much rarer and thus provide low power for statistical tests of association. CNVs encompassing *Cyp6p3* and *Cyp6p5* exist at appreciable frequencies in *An. gambiae* from Madina (17% and 21% of samples showing increased copy number respectively) but with no significant association with resistance.

**Figure 2.**
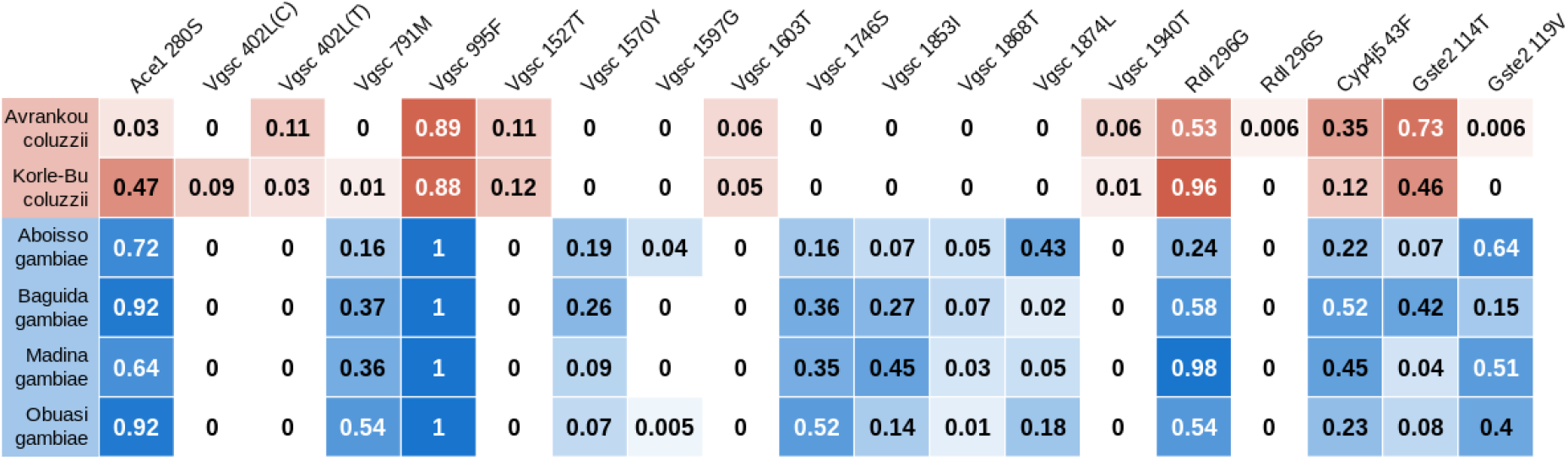
Allele frequencies for SNPs in known resistance loci. No wild type haplotypes were found in *Vgsc*, with all samples carrying either the 995F mutation (100% of *An. gambiae* samples) or the 402L / 1527T combination. Cells are colour-coded according to species (*An. gambiae* in blue, *An. coluzzii* in red) with darkness related to allele frequency. SNPs that were completely absent are not shown.

**Figure 3.**
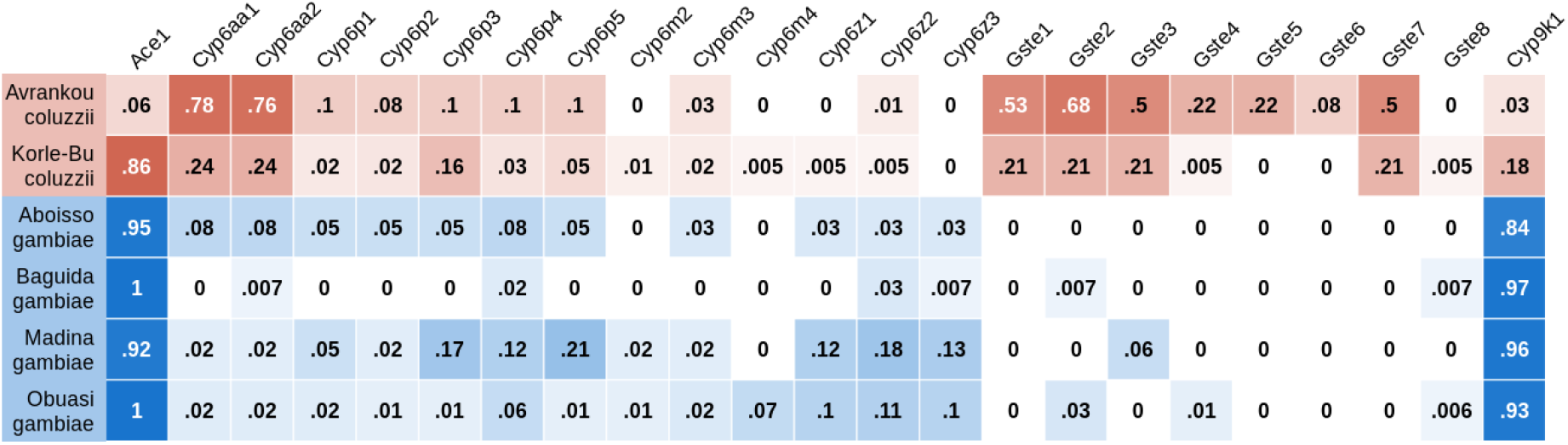
Proportion of samples with increased copy number in genes from four major resistance loci (*Ace1*, the *Cyp6aa* / *Cyp6p* cluster, the *Gste2* cluster and *Cyp9k1*). Cells are colour-coded according to species (*An. gambiae* in blue, *An. coluzzii* in red) with darkness related to allele frequency. SNPs that were completely absent are not shown.

When breaking down the *Cyp9k1* CNVs into distinct alleles (Supplementary Fig. S2), it is apparent that one allele (Cyp9k1_Dup10) is shared by both *An. gambiae* and *An. coluzzii*. Haplotype clustering analysis showed that the CNV in both species was present on the same genetic background (Supplementary Fig. S3), which was nested within the *An. coluzzii* part of the haplotype tree, indicating that the mutation spread through introgression from *An. coluzzii* to *An. gambiae*. Despite this introgression of a *Cyp9k1* CNV between species, there was no association between *Cyp9k1* copy number and resistance to either deltamethrin or PM.

The CNV in *Ace1*, known to be associated with PM resistance (Grau-Bové et al., 2021), was at high frequency (>85%) in all populations except *An. coluzzii* from Avrankou (Fig. 3, Supplementary Fig S2), and *Ace1* copy number was strongly associated with resistance to PM (P < 10^-5^ in all cases) in all populations except Baguida. No other genes showed a significant association between copy number and resistance.

### 2.3. Windowed measures of differentiation / selection to identify genomic regions associated with resistance

We identified regions of the genome associated with phenotype within each sample set using three different metrics (F_ST_, PBS and difference in H_12_ signal between resistant and susceptible subsets) calculated in 1000 SNP windows (Supplementary Data S2,S4,S5, Supplementary Fig. S4-S7). F_ST_ is a measure of genetic differentiation between two groups of samples but does not indicate whether either group displays signals of selection, which is expected if the genetic difference is associated with resistance. Peaks of F_ST_ were therefore investigated further to identify high-frequency haplotypes significantly associated with resistance (Supplementary Data S3). PBS (Yi et al., 2010) is a measure of selection that is particularly effective at identifying recent selection from standing genetic variation. PBS uses F_ST_ to identify genomic regions showing greater evolutionary change in one group (here, the resistant samples) relative to a closely related group (susceptible samples) and an outgroup. While originally designed to detect positive selection, it has also been used to detect phenotypic association (Grau-Bové et al., 2021). H_12_ is a measure used to detect genomic regions undergoing selective sweeps (Anopheles gambiae 1000 Genomes Consortium, 2017; Garud et al., 2015). To identify regions in which swept haplotypes are more frequent in resistant compared to susceptible individuals, we calculated the difference in H_12_ value between groups, which we refer to as ΔH_12_.

Based on these metrics, the cytochrome P450 *Cyp9k1* was frequently associated with resistance to both deltamethrin and PM, but curiously the association signal never localised to the gene itself (Supplementary Fig. S4, S8). We found signals on the telomeric side (Madina deltamethrin, Madina PM, Obuasi PM) and the centromic side (Avrankou deltamethrin, Obuasi deltamethrin, Baguida PM, Korle-Bu PM) or both (Korle-Bu deltamethrin), with distances from *Cyp9k1* ranging from 200 to 1200 Kbp. This was not due to an absence of windows closer to *Cyp9k1* itself. We also found recurrent, cross-insecticide signals in Thioester-containing protein (*Tep*) genes, with peaks in *Tep1* (AGAP010815) for Obuasi PM, and *Tep4* (AGAP010812) for Madina deltamethrin (Fig. S4).

For deltamethrin, the most consistent signal was at the *Cyp6aa* / *Cyp6p* gene cluster. At this locus, we found PBS and F_ST_ peaks in *An. gambiae* from Obuasi and Madina, and a peak in ΔH_12_ in *An. coluzzii* from Korle-Bu. The strongest of these signals came from the Obuasi sample set, where the F_ST_ peak spanned 17 windows spread over nearly 250 Kbp. The window covering the *Cyp6aa* / *Cyp6p* cluster contained a haplotype cluster that accounted for over a third of the sample set (56 out of 156 haplotypes) which was positively associated with resistance to deltamethrin (*P* = 0.005). In Madina, the peak consisted of two significant windows, the nearest one being around 6 Kbp away from the start of the cluster. This window contained a haplotype group accounting for over half (109 out of 202) of the haplotypes in the sample set and was positively associated with resistance to deltamethrin (*P* = 0.002).

To determine whether the haplotypes driving the F_ST_ signal in Obuasi and Madina were the same, we combined the deltamethrin-phenotyped samples from these two locations and re-analysed the significant window covering *Cyp6aa1*. The main haplotype cluster was indeed the same in Obuasi and Madina (Fig. 4) and the significance of association for the combined analysis was increased (*P =* 0.001). The haplotype cluster contained many SNPs (Fig. 4, Supplementary Data S2), of which four were non-synonymous in *Cyp6aa1* (2R positions 28480960:N501I, 28480993 + 28480994 (combine to make S490I) and 28482335:Y77F) and one in *Cyp6aa2* (28483525: M428V), but none of these SNPs were completely absent from the non-cluster haplotypes, suggesting that they may not be driving the sweep.

**Figure 4.**
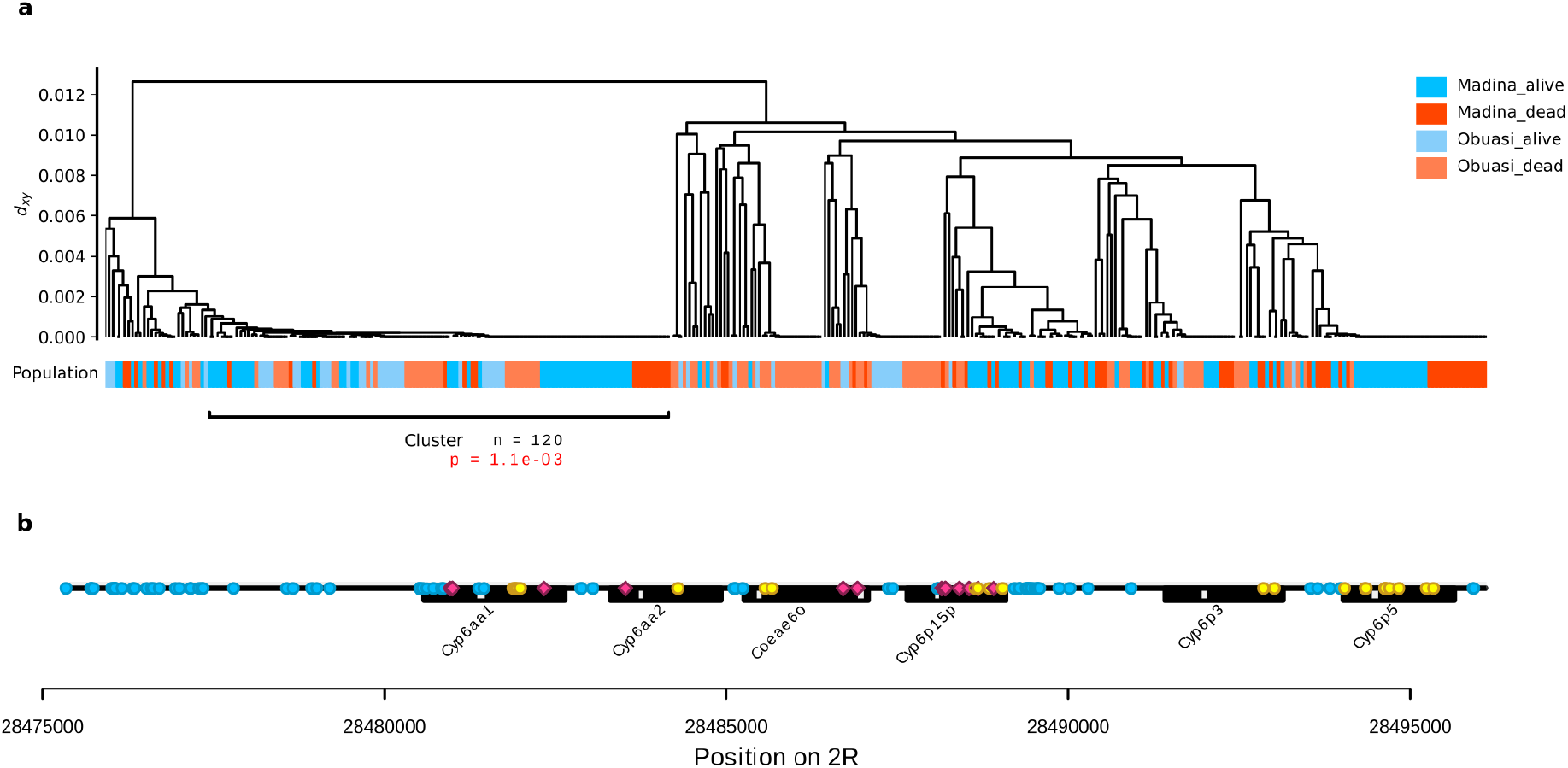
A shared haplotype cluster around *Cyp6aap* cluster at high frequency in *An. gambiae* from Madina and Obuasi is positively associated with resistance to deltamethrin. (a) hierarchical clustering dendrogram of haplotypes with leaves labelled by population and phenotype. The tree was cut at a height of d_xy_ = 0.001 to obtain haplotype clusters. (b) SNPs that were significantly more frequent on the haplotype than in the rest of the population (Fisher test, alpha = 0.001) are shown and labelled as non-synonymous (red), synonymous (yellow) and non-coding (blue, including UTRs).

Several peaks were also associated with genes from the superfamilies of metabolic genes typically associated with resistance (P450s, GSTs and carboxylesterases). We found PBS signals at *Cyp6m2* (Obuasi), *Cyp6ag1/2* (Avrankou) and *Cyp6aj1* (Madina) and an F_ST_ signal near the carboxylesterase *Coe22933* in *An. coluzzii* from Korle-Bu (Fig. S4). In *An. coluzzii* from Avrankou, one F_ST_ peak was adjacent to the P450 redox partner gene NADPH cytochrome P450 reductase (*Cpr*, AGAP000500) and contained a haplotype group positively associated with resistance (*P* = 0.048). There were also several independent signals in NADH dehydrogenase (ubiquinone) 1, with significant PBS peaks near beta subunit 2 (AGAP002630) in both *An. gambiae* from Madina and *An. coluzzii* from Korle-Bu. Furthermore, the PBS peak near *Cyp9k1* in Korle-Bu covers NADH dehydrogenase beta subunit 1 (AGAP000849), but the proximity to *Cyp9k1* makes it hard to determine whether this is coincidental.

For PM, we found strong peaks in the *Ace1* region in all populations except *An. gambiae* from Baguida (Supplementary Data S2). Additionally, in *An. coluzzii* from Korle-Bu, a PBS and F_ST_ peak that included the *Gste* gene family contained a haplotype cluster negatively associated with PM resistance (*P* = 0.003) and a small cluster (n = 30 haplotypes out of 210) positively associated with resistance (*P* = 0.03). There was also a significant peak in ΔH_12_ between resistant and susceptible *An. gambiae* from Madina at the *Gste* locus (Fig. S4, S6).

In *An. gambiae* from Obuasi, the broadest peak consisted of 19 windows on chromosome 3L ranging from 30477135 to 30697180, which includes methuselah-like protein 7 *(Gprmthl7*, AGAP011643) and is around 23 kbp away from an odorant receptor *Or11* (AGAP011631). However, the haplotype cluster in these windows is negatively associated with PM resistance (*P* < 0.0006). Interestingly, there was also a F_ST_ / PBS peak of a single window on chromosome 3R which contained the gene coding for the beta-2 subunit of nicotinic acetylcholine receptor (AGAP010057), the target site of neonicotinoid insecticides. The main haplotype in this region was positively associated with PM resistance (*P* = 0.0008) and contained a non-synonymous SNP (N418Y).

Some sample sets (Avrankou deltamethrin, Korle-Bu PM, Madina PM) also showed phenotypic associations for many SNPs within the 2La chromosomal inversion. Strong linkage disequilibrium and the large size of this inversion makes identification of the region responsible for the association unfeasible.

### 2.4. GWAS analysis

As well as the windowed approach above, we implemented a SNP-wise association analysis across the whole genome in each sample set. Preliminary analysis identified significant associations that were later revealed to be an artefact of bacterial sequence data; we therefore repeated the analysis after removing SNPs associated with this contamination (Supplementary Methods).

After false-discovery rate control, no SNPs remained significant for deltamethrin resistance in any of the populations, while the PM sample sets either contained no significant SNPs (Baguida and Obuasi) or predominantly SNPs in the *Ace1* region (Korle-Bu and Madina). This likely reflects a combination of the effect size of resistance mutations, the sample sizes in our study, and the stringent false discovery rate control when such a large number of tests are conducted. We reasoned that if mutations in a given region or cluster of genes can be associated with resistance, several mutations in the same region may all contribute separately to resistance, or may lead to association of nearby variants through selective sweeps. We would then expect to see groups of clustered variants all associated with resistance. We therefore adopted an alternative approach in which we took the 1000 most highly significant SNPs in each sample set and looked for regions of the genome in which these SNPs were concentrated (100,000 bp windows that contained at least 10 SNPs from among the top 1000, Supplementary Data S6). For convenience, we refer to these as candidate SNPs.

For deltamethrin, in *An. coluzzii* from Avrankou, the majority of the candidate SNPs were found across the 2La chromosomal inversion, mirroring what was found in the F_ST_ / PBS analysis. Aside from this, there were no candidate SNPs within gene sequences. In *An. gambiae* from Baguida, candidate SNPs were found in several regions of the genome, including a cluster of *Tep* genes *(Tep2* (AGAP008366), *Tep14* (AGAP008368) and *Tep15* (AGAP008364)), with a non-synonymous mutation in *Tep14*. Interestingly, we also found a window of candidate SNPs in *An. coluzzii* from Korle-Bu that was around 50 kbp from a different cluster of *Tep* genes, including *Tep1* (AGAP010815). We also found candidate SNPs around the cytochrome P450 *Cyp306a1*, including one within a splice site, and around the carboxylesterase *Coe22933*. In *An. gambiae* from Obuasi, the main signal comes from the region around *Cyp6aa1*, as was found with the F_st_ analysis.

For PM, in *An. coluzzii* from Korle-Bu and *An. gambiae* from Madina, all clusters of candidate SNPs were found in the *Ace1* region. In other populations, candidate SNPs were found in and around several clusters of possible detoxification genes (the carboxylesterases *Coeae2g-6g* in *An. gambiae* from Obuasi, and the P450s *Cyp12f1-f4, Cyp4d15-17, Cyp4k2, Cyp4ar1, Cyp4h19* and *Cyp4h24* in *An. gambiae* from Baguida). In Baguida, there was also a cluster of candidate SNPs covering a region around 39.2Mb on chromosome 2L, containing many cuticular proteins (RR1 family AGAP006838-AGAP006867). In *An. gambiae* from Obuasi, as well as a group in *Ace1*, candidate SNPs were found near gustatory receptors *(Gr26* (AGAP006717) & *Gr27* (AGAP006716)) and a cluster of *Tep* genes (including five non-synonymous mutations in *Tep1)*, as well as around *Gprmthl7* (including three non-synonymous mutations), as found in the F_ST_ analysis.

### 2.5 GWAS sample sizes

Although we found significant associations of individual SNPs in our analysis of established resistance markers for deltamethrin resistance, our agnostic SNP-level GWAS returned no markers passing FDR control. We therefore explored what sample size would have been required in order for these significant established markers to have been detected in the agnostic genome-wide analysis. Over 500 simulations, we found that a sample size of 300 (150 of each phenotype) would only have detected 11% or 25% of associated SNPs, depending on whether we modelled the observed allele frequencies of *Vgsc*_1527T or *Cyp4j5_43F*, respectively, in Avrankou. However, with a sample size of 500, this rose to 54% and 67%.

## 3. Discussion

Overall, our results show different genomic signals of resistance to deltamethrin and PM, but with clear points of overlap. The pyrethroid deltamethrin is a common constituent of ITNs and has been a mainstay of vector control since the end of the 20th century (Choi et al., 1995). The first genetic marker of resistance against pyrethroids in *An. gambiae*, discovered almost 25 years ago, was the target site mutation *kdr-1014F*, now referred to by Anopheles numbering as *Vgsc*-995F (Martinez-Torres et al., 1998). Since then, this target site resistance has spread and massively increased in frequency (Hancock et al., 2022), with at least five independent origins of the mutation in Africa, leaving the wild-type allele completely absent in our current dataset (collected in 2017). A recent decline in the frequency of *Vgsc*-995F in *An. coluzzii* is associated with the rise of an alternative resistance haplotype in the *Vgsc* gene, carrying the *Vgsc-*402L and *Vgsc*-1527T mutations instead (Clarkson et al., 2021). While there is debate as to the relative benefits of *Vgsc*-995F vs *Vgsc-*402L/152T, both provide target site resistance (Williams et al., 2022). It is in this context of ubiquitous target site resistance to deltamethrin that our study was conducted, making it poised to address three questions: Is there a difference in the level of resistance conferred by *Vgsc*-995F and the *Vgsc*-402L/1527T haplotype? Do other SNPs around *Vgsc*-995F provide additional resistance to deltamethrin? And what other mutations in the genome help to explain the residual variation in resistance?

We found that *Vgsc*-402L/1527T was associated with significantly higher levels of resistance to deltamethrin than *Vgsc*-995F. This is opposite to what has previously been found in laboratory colonies (Williams et al., 2022). This difference may be explained by the effect of additional non-synonymous mutations on the *Vgsc*-995F haplotype background. Such mutations were not exhaustively investigated in the laboratory colonies, and the relatively increased resistance of the *Vgsc*-995F haplotype in the colonies could thus be explained if this mutation was backed up by supporting SNPs elsewhere, or by differences in advantage between field and laboratory conditions.

Outside of the *Vgsc*, the main deltamethrin resistance-associated signals that we found were in and around the *Cypaa* / *Cyp6p* cluster of cytochrome P450 genes. The importance of *Cyp6aa1* for deltamethrin resistance has been previously demonstrated in *An. gambiae* from East and Central Africa, where the CNV Cyp6aap_Dup1 was found to be associated with resistance and has spread rapidly (Njoroge et al., 2022). Similarly in our data, we found that copy number of *Cyp6aa1* was associated with resistance in at least one population of *An. coluzzii*. Unlike East Africa, where a single CNV allele dominates, we found at least 6 CNV alleles in our sample sets of *An. coluzzii*, and more are known from other data (Lucas, Miles, et al., 2019; The Anopheles Gambiae 1000 Genomes Consortium, 2021).

We also found signals of association with deltamethrin resistance around *Cyp6aa1* in two populations of *An. gambiae* (Madina and Obuasi), leaving Baguida as the only population in which no evidence of *Cyp6aa1* association was found. Interestingly, the signal in Madina and Obuasi was not associated with any CNV, suggesting that an alternative mechanism of resistance has appeared in these populations. This putative non-CNV resistance haplotype, along with the large number of CNVs in the region, suggests that metabolic resistance associated with *Cyp6aa1* is likely to be highly multi-allelic.

Other signals of association with deltamethrin resistance were found around the carboxylesterase *Coe22933*, the P450s *Cyp9k1, Cyp6m2, Cyp6ag1/2, Cyp6aj1, Cyp306a1*, as well as *Cpr* (the obligatory cytochrome P450 electron donor, whose knockdown increases susceptibility to permethrin (Lycett et al., 2006)), pointing to the existence of further metabolic resistance mechanisms. There was also a suggestive signal of association of deltamethrin resistance with the beta subcomplex of NADH dehydrogenase (ubiquinone), a large mitochondrial complex of the respiratory chain. We found such signals in populations from two species (*An. gambiae* from Madina and *An. coluzzii* from Korle-Bu), and possibly two regions of the genome (subunit 2 on chromosome 2R and subunit 1 on chromosome X), although the proximity of subunit 1 to *Cyp9k1* makes it hard to establish whether this is genuinely an independent signal of association. A previous study of the loss and re-establishment of deltamethrin resistance in a laboratory colony of *An. coluzzii* from Burkina Faso found that resistance was associated with elevated expression of genes with NADH dehydrogenase activity or involved in cellular respiration (Ingham et al., 2021). Our results suggest that this putative mechanism of resistance might be present in other populations and species.

In contrast to deltamethrin, PM has been introduced more recently as a tool for mosquito control (Oxborough, 2016), although previous use of organophosphates as agricultural insecticides is likely to have led to prior exposure of at least some mosquito populations (Essandoh et al., 2013). Furthermore, while *Vgsc* target site resistance to deltamethrin consists of presence / absence of a SNP, target site resistance to PM, through the *Ace1*-280S SNP, is moderated by the copy number of the CNV in *Ace1* (Grau-Bové et al., 2021). This copy number is variable, and is likely to be highly mutable as recombination between CNV haplotypes leads to changes in copy number. For these reasons, variation in target site resistance persists, and our results show that this locus continues to dominate the association analyses in several populations. An exception is *An. gambiae* from Baguida, where there was no significant association with either the CNV or the *Ace1*-280S SNP, despite the frequencies of both being similar to that seen in Obuasi, where the association was significant.

Beyond *Ace1*, we found signals of association with PM resistance around cytochrome P450s (Cyp4k1, Cyp12f, Cyp4h, Cyp9j5) and carboxylesterases *(Coeae2g-6g)*, as well as at the glutathione S-transferase epsilon (*Gste*) genes. Expression and sequence mutation in *Gste2* has been associated with resistance, primarily to DDT (Adolfi et al., 2019; Mitchell et al., 2014; Riveron et al., 2014), but also to pyrethroids (Lucas, Rockett, et al., 2019; Riveron et al., 2013, 2014). While direct evidence of an association with resistance to PM is lacking, GSTs are known to detoxify organophosphates and have frequently been associated with resistance to them (Enayati et al., 2005). This includes studies in the mosquito *Aedes aegypti*, where there is evidence of association between *Gste2* (both expression and gene sequence) and resistance to the larvicide temephos (Saavedra-Rodriguez et al., 2014), and *An. gambiae*, where induced over-expression of *Gste2* led to elevated resistance to fenitrothion (Adolfi et al., 2019).

There was also a broad signal of resistance association around the *Gprmthl7*, named after the *Drosophila* gene *Mth*, which is associated with extended lifespan (Lin et al., 1998). However, *Mth* itself is restricted to *Drosophila*, and the function of *Mth*-like genes reported in other insects is not well understood (Li et al., 2014). Another intriguing signal was found at the beta-2 subunit of nicotinic acetylcholine receptor, where we found a haplotype with relatively strong association with PM resistance (*P* = 0.0008). This receptor is the target site for neonicotinoid insecticides (Bass et al., 2015) and is not known to be involved in resistance to PM, but its function is to respond to acetylcholine, the neurotransmitter that is broken down by *Ace1*. We can speculate that mutations in this receptor might in some way protect against the toxic accumulation of acetylcholine that occurs when *Ace1* is inhibited by PM.

Curiously, we also found marginally significant signals of association between PM resistance and SNPs in the *Vgsc* region (*Vgsc*-1570Y positively associated in Baguida and *Vgsc*-995F negatively associated in Korle-Bu. The negative association could be explained if the physiological cost of carrying the *Vgsc*-995F mutation compared to *Vgsc-*402L/1527T (Grigoraki et al., 2021; Williams et al., 2022) contributed to weakness in the face of stress. The effects of the *Vgsc*-1570Y mutation on fitness costs is unknown. If part of its effect is to reduce the fitness cost of *Vgsc*-995F, this may similarly explain its effect on PM resistance. These remain speculative arguments that require further investigation.

Two groups of signals were consistently found associated with resistance to both deltamethrin and PM. Given the importance of cross-resistance in the management of insecticide-based mosquito control strategies, these regions warrant further discussion. The first such signal was around *Cyp9k1*, a cytochrome P450 found in a region of strong selective sweep (Anopheles gambiae 1000 Genomes Consortium, 2017), and that has been implicated in resistance to pyrethroids (Main et al., 2015), and shown to metabolise deltamethrin *in vitro (Hearn et al., 2022; Vontas et al., 2018)*. We found repeated signals of association of *Cyp9k1* with resistance to both deltamethrin and PM, although the signal was always several hundred, or even a thousand, Kbp away from *Cyp9k1* either side. This makes it difficult to confirm whether *Cyp9k1* is driving these signals. On the one hand, the known importance of *Cyp9k1* and the presence of significant windows on both sides, suggests that we may be seeing a broad signal of association that for some reason is weaker at the locus of the gene itself. Indeed, there are no known non-synonymous mutations associated with resistance in *Cyp9k1*, with the only evidence being haplotypic (Main et al., 2015) and metabolic, and regulatory regions may be located far from the gene itself. On the other hand, genes such as NADH dehydrogenase and *Cyp4g17* were often closest to the significant window, or even within the windows themselves. *Cyp4g17* has been found to be over-expressed in *An. coluzzii* from the Sahel resistant to deltamethrin (Ibrahim et al., 2022), and the potential role of NADH dehydrogenase has been discussed above. The large number of CNV alleles found in *Cyp9k1*, and the introgression of one of these alleles from *An. coluzzii* to *An. gambiae*, also suggests an important role for this protein in resistance, but we have yet to find a significant association between *Cyp9k1* copy number and resistance. The exact role that *Cyp9k1* and the genes around it play in resistance to both deltamethrin and PM remains poorly understood.

The other signals of association that were found for both insecticides were in *Tep* genes, key components of the arthropod innate immune system, with *Tep1* being known for its role in immunity against *Plasmodium* (Blandin et al., 2004; Cirimotich et al., 2010). Our results lend further weight to a recent study of *An. coluzzii* in several regions of the Sahel, which found that *Tep1* was consistently over-expressed in mosquitoes resistant to deltamethrin (Ibrahim et al., 2022). It is intriguing to see *Tep1*, and other members of the *Tep* gene family, consistently associated with genomic signals of resistance to two insecticides, in both *An. gambiae* and *An. coluzzii*. While there is evidence that exposure to *Plasmodium* increases susceptibility to insecticides (Saddler et al., 2015), inviting speculation that improving immunity could improve resistance, that study found that it is exposure to the parasite, rather than infection, that causes susceptibility. The mechanisms for this link between the immunity pathway and resistance requires further investigation.

When performing SNP-wise GWAS on the data, we found very few significant SNPs after false discovery rate control. This prompted us to turn to a windowed approach, which in effect makes use of the non-independence of adjacent SNPs, particularly in regions of positive selection, to greatly reduce the number of loci being considered without overly diluting the association signals. The drawback is that this approach reduces the power to home in on the precise region most closely linked to the phenotype. In our targeted analyses of loci with *a priori* expectations of effect (SNPs and CNVs in known resistance genes), the number of tests being conducted was considerably smaller and the range of *P* values that we found to be significant, with the exception of *Ace1*, would not come close to significance if FDR control is applied on a genomic scale such as in the GWAS. We conclude that the scale of the effect size for mutations associated with residual resistance (after the major loci such as *Ace1* and *Vgsc* are accounted for) are such that larger sample sizes than those used here (or in any insect GWAS study) are required in order for their *P* values to stand out in a GWAS. Our simulations suggest that a sample size of 500, with a dataset of 7 million SNPS, of which 10 are associated with resistance, would detect 54% of SNPs that have the effect size that we observed for *Vgsc*_1527T in Avrankou, and 67% with the effect size observed for *Cyp4j5*_43F. This is an encouraging figure, and a reasonable one to achieve given current sequencing costs and throughput.

A striking result from our findings is that signals of non-target site resistance showed little consistency between our study sites. Apart from the *Cyp6aa1* haplotype associated with deltamethrin resistance in *An. gambiae* from Madina and Obuasi, few other signals were shared between populations, despite several repeated themes (*Cyp6aa1* resistance also present in *An. coluzzii*, *Tep* genes, P450s, carboxylesterases) and despite several documented instances of introgression between *An. gambiae* and *An. coluzzii* (*Vgsc* (Clarkson et al., 2014), *Ace1* (Grau-Bové et al., 2021), Rdl (Grau-Bové et al., 2020), *Cyp9k1* (this study)). These findings, along with the sheer number *Cyp6aa1* CNV alleles discovered so far (at least 30, (The Anopheles Gambiae 1000 Genomes Consortium, 2021)), suggest that metabolic insecticide resistance in *An. gambiae* and *An. coluzzii* is highly multiallelelic, with even nearby populations displaying different resistance mutations. This is in contrast to what has been observed with target site resistance, where various arrangements of a single SNP / CNV combination in *Ace1* dominate for PM resistance, while a few different mutations in *Vgsc* seem to be involved in deltamethrin resistance. At the *Vgsc-995* locus for example, while at least 10 different origins of resistance have been identified across Africa (Anopheles gambiae 1000 Genomes Consortium, 2017; Clarkson et al., 2021), nine of which are mixing in the Democratic Republic of Congo (Lucas, Rockett, et al., 2019), these are parallel origins which together account for only two different changes in the same amino acid. It is also in contrast to *An. funestus*, where target site resistance has yet to be found in any populations, but where a few mutations associated with metabolic resistance have spread extensively in Africa (Mugenzi et al., 2019; Mulamba et al., 2014; Riveron et al., 2014; Weedall et al., 2019).

This multi-allelic nature of metabolic resistance presents a twofold challenge for genetic monitoring programs. First, if the mutations that cause resistance, or can be used to track it, are different in every population, methods that target individual SNPs are unlikely to be generalisable. This includes probe-based technologies such as MassArray iPlex, LNA and Taqman. This challenge can be overcome by turning to more general methods, that target broader loci, such as amplicon sequencing, where regions the size of one or several amplicons can be investigated, thus remaining agnostic as to the specific change that could occur within that region.

Second, it is more challenging to build predictive models of resistance if each population has its own resistance markers and thus requires its own model. Metabolic resistance is often driven by increased expression of certain genes, and part of the reason for its multi-allelic nature is that there are many ways by which expression can be increased (copy number variants, mutation of cis-regulatory region, modified transposable element activity). To capture all of these in a single assay, it may be necessary to begin thinking about the possibility of transcriptomic or even proteomic, rather than simply genomic surveillance (Mavridis et al., 2019). This brings its own challenges. While the genetic sequence of an organism is largely stable, gene expression and protein synthesis are variable across tissues, life stages and environments. Collection methodologies therefore need to be refined in order to achieve comparable results across studies. Also, RNA degrades much more quickly than DNA after death, and collecting samples for gene expression analyses therefore requires greater care. It is however becoming increasingly apparent that these challenges should be addressed and these avenues explored.

We have conducted a large-scale GWAS of insecticide resistance, with data from 827 (after QC and removal of males and siblings) individuals from five locations and two insecticides. The presence of full siblings in our data reflects our sample collection from larval pools. Since females lay eggs in batches, and larval breeding habitats offer little scope for dispersal, collection of individuals from the same egg batch cannot be avoided with any certainty. Larval collections are typical for insecticide resistance bioassays, because females of known age are needed to maintain a standard process (World Health Organization, 2016). We therefore expect that this problem would be frequent in phenotypic association experiments, yet relatedness is rarely investigated since it requires larger scale genotyping data that would be obtained when only a few markers are investigated.

## 4. Methods

### 4.1 Sample collection and phenotyping

Mosquitoes were collected in 2017 as larvae from six locations in West Africa (Benin: Avrankou, Côte d’Ivoire: Aboisso, Ghana: Madina, Korle-Bu and Obuasi; Togo: Baguida, Fig. 1). Larvae were raised to adulthood in the laboratory and females were phenotyped for either deltamethrin or PM using a custom dose-response assay with WHO standard tubes, which was designed to identify the most resistant and susceptible individuals, while removing those of intermediate resistance (Supplementary Methods).

DNA was extracted from individual mosquitoes using nexttec extraction kits. Species identity was determined using two molecular methods designed to discriminate between *An. gambiae, An. coluzzii* and *An. arabiensis:* a PCR of species-specific SINE insertion polymorphisms as described in (Santolamazza et al., 2008), and a melt curve analysis (Chabi et al., 2019). The complete list of specimens, sampling times and locations, and species assignments are available in Supplementary Data S1.

Sample sets (mosquitoes of a given species, from a given location, exposed to a given insecticides) to send for sequencing were chosen on the basis of sample size and to obtain a balance of deltamethrin / PM data (Supplementary Methods). Final sample sizes for each sample set, after QC filtering of sequencing data, are shown in Table 1.

### 4.2 Whole genome sequencing and bioinformatic analysis

Overall, 1258 samples from this study were whole-genome sequenced as part of the *Anopheles gambiae* 1000 genomes project (Ag1000G) release v3.2. Full details of library preparation, sequencing, alignment, SNP calling, CNV calling and phasing are detailed on the Ag1000G website (https://malariagen.github.io/vector-data/ag3/methods.html).

Briefly, individuals were sequenced to a target coverage of 30x on an Illumina HiSeq X, generating 150 bp paired-end reads. Reads were aligned to the AgamP4 reference genome using BWA, and indel realignment was performed using GATK version 3.7-0. Genotypes were called for each sample independently using GATK version 3.7-0 UnifiedGenotyper in genotyping mode, given all possible alleles at all genomic sites where the reference base was not “N”. Sample QC removed 62 samples for low coverage (< 10x), 215 samples for cross-contamination (*alpha* > 4.5%, (Jun et al., 2012) and 8 samples as apparent technical replicates (genetic distance below 0.006). 973 samples passed QC filtering. Sex was called using the modal coverage ratio between chromosomes X and 3R (ratio between 0.4-0.6 = male, ratio between 0.8-1.2 = female, other ratios would lead to sample exclusion).

Known CNVs in specific genes of interest (Cyp6aa1–Cyp6p2, Gstu4–Gste3, Cyp6m2–Cyp6m4, Cyp6z3–Cyp6z1, Cyp9k1, Ace1) were detected using discordant reads associated with CNV alleles previously identified in Ag1000G release 3.0. Agnostic CNV detection was performed using normalised sequencing coverage calculated in 300 bp windows and a hidden markov model (HMM) to estimate the copy number state at each window. This allows the detection of CNVs genome-wide, and of novel CNV alleles in the regions of interest. Gene copy number was calculated as the modal value of the HMM along each gene. A novel CNV in the regions of interest was identified if there was increased copy number according to modal coverage, but no discordant reads supporting the presence of known CNV alleles.

### 4.3 Kinship analysis

We calculated pairwise kinship between all samples using the KING statistic (Manichaikul et al., 2010) implemented in NGSRelate (Hanghøj et al., 2019) using SNP data across the whole genome. Whole genome SNPs were used because the recombination rate on such a small genome can lead to large disparity in kinship values between chromosomes. Results indicated a slight positive bias in kinship, with the mode of the distribution slightly above 0 (Supplementary Methods). We identified full siblings as any pair of individuals with a kinship value greater than 0.195 (Supplementary Methods) and obtained full sib groups by considering that any siblings of siblings were themselves siblings. No full siblings were found between populations.

### 4.5 Candidate marker and CNV association analysis

We investigated the association between phenotype and a range of known SNPs in candidate resistance genes (Ace1-280S all *Vgsc* SNPs reported in (Clarkson et al., 2021), Rdl-296G, Rdl-296S, Cyp4j5-43F, Gste2-114T and Gste2-119V) using generalised linear models in R, with binomial error and a logit link function, with phenotype as the dependent variable and SNP genotypes as independent variables, coded numerically as the number of mutant alleles (possible values of 0, 1 and 2). Within each sample set, we included all SNPs with an allele frequency of at least 10% in the analysis. We used a forward step-wise procedure, calculating the significance of adding each marker to the current model using the *anova* function. Starting from the null model, we added the most significant marker to the model and then repeated the process until no remaining markers provided a significant improvement. We used the same procedure to investigate the phenotypic association of gene copy number.

### 4.6 Windowed measures of differentiation (F_ST_, PBS, H_12_)

FST in 1000 SNP windows was calculated between resistant and susceptible samples within each sample set using the *moving_patterson_fst* function in *scikit-allel*, after filtering SNPs for missing data and accessibility (The *Anopheles gambiae* 1000 Genomes Consortium, 2021) and removing singletons. In order to take full advantage of the full sample set despite the non-independence of full siblings, we performed up to 100 permutations in which one randomly-chosen individual per sib group was used in the calculation of F_ST_, and took the mean of all permutations. The set of possible permutations was sampled without replacement, such that in some sample sets fewer than 100 permutations were possible. Provisional windows of interest (“peaks”) were identified as ones with positive F_ST_ values three times further from the mode than the smallest negative value (Supplementary Methods).

The existence of extended haplotype homozygosity in a region (due to a selective sweep) could cause a peak in windowed F_ST_ even if the swept haplotype is unrelated to the phenotype, because non-independence of SNPs in the window would lead to increased variance in F_ST_ compared to other genomic regions. This led, for example, to spurious provisional peaks in the *Ace1* region in sample sets phenotyped against deltamethrin (Fig. S5). To filter out these peaks, we performed 200 simulations in which the phenotype labels were randomly permuted and F_ST_ recalculated as above. Provisional windows of interest were retained if their observed F_ST_ was higher than the 99th centile of the simulations.

H_12_ was calculated using phased biallelic SNPs in 1000 SNP windows, using the *garuds_h* function in scikit-allel. 200 phenotype permutations were performed as above. PBS was calculated using segregating SNPs in 1000 SNP windows using the *pbs* function in scikit-allel, after filtering SNPs for accessibility using the Ag1000G phase 3 gamb_colu site mask. For the outgroup for the PBS calculation, we used conspecific samples from Mali collected in 2004, available as part of the Ag1000G phase3 data release. For both H_12_ and PBS, phenotype permutations were performed as for F_ST_ to filter out false positives caused by the presence of extended swept haplotypes.

### 4.7 Haplotype association

Within each window of interest identified through the F_ST_ analysis, we explored the presence of swept haplotypes that could be associated with phenotype. Haplotype clusters were determined by hierarchical clustering on pairwise genetic distance (Dxy) between haplotypes, and cutting the tree at a height of 0.001. Clusters larger than 20 haplotypes were tested for association with phenotype using a generalised linear model with binomial error and logit link function, with phenotype as the response and sample genotype (number of copies of the haplotype) as a numerical independent variable.

### 4.8 Genome-wide association analysis

In the GWAS, a single permutation of sibling removal was randomly chosen for the analysis, since averaging over permutations would not produce interpretable *P*-values.

In each sample set, all SNPs passing accessibility filters (The Anopheles gambiae 1000 Genomes Consortium, 2021) with no missing data and a minor allele count of at least five were included in the analysis, including sites that failed accessibility filters. Preliminary investigation revealed that some of our samples contained contamination by *Asaia* bacteria which interfered with SNP calling at some sites (Supplementary Methods). We used Bracken (Lu et al., 2017) to estimate the amount of *Asaia* contamination in each sample and excluded SNP loci where genotype was correlated with *Asaia* levels (*P* < 0.05).

*P*-values for each SNP were obtained by generalised linear modelling with binomial error and logit link function, with phenotype as the response variable and genotype (number of non-reference alleles) as a numeric response variable. False discovery rate correction was applied using the *fdrtool* package in R (Klaus & Strimmer, 2015). In several sample sets, no significant *P* values remained after false discovery rate correction. Since recent selection on a SNP should lead to a broad selection signal, we took the 1000 most significant SNPs in each sample set and looked for 100,000bp windows that contained at least 10 SNPs among the top 1000. Effects of individual SNPs were determined using SNPeff (Cingolani et al., 2012).

### GWAS sample size analysis

Although we found significant associations of individual SNPs in our analysis of established resistance markers for deltamethrin resistance, our agnostic SNP-level GWAS returned no markers passing FDR control. We therefore asked what sample size would have been required in order for these significant established markers to have been detected in the agnostic genome-wide analysis. Taking the *Vgsc*_1527T and *Cyp4j5_43F* loci in Avrankou, we used the observed allele frequencies in each of the resistant and susceptible subsamples to randomly generate new sample sets of a given size, split into 50/50 resistant/susceptible samples. We thus maintained the observed effect sizes for these loci, projected onto new sample sizes. For each random sample set, we calculated *P* values following the same procedure as for our GWAS. FDR correction depends on the number of truly associated and unassociated genes in the analysis; we therefore simulated 10 such loci that were truly associated with resistance across the genome (each drawn independently) and 7 million non-associated SNPs (approximately 7 million SNPs were used in the Avrankou GWAS), whose *P*-values were drawn from a random uniform distribution. This simulation was performed 500 times for each sample size.

## Availability of data and materials

Code used to analyse the data can be found in the github repository https://github.com/vigg-lstm/GAARD_work. All sequencing, alignment, SNP and CNV calling was carried out as part of the *Anopheles gambiae* 1000 genomes project v3.2 (https://www.malariagen.net/data).

## Acknowledgements

This work was supported by the National Institute of Allergy and Infectious Diseases ([NIAID] R01-AI116811) with additional support from the Medical Research Council (MR/P02520X/1). The latter grant is a UK-funded award and is part of the EDCTP2 programme supported by the European Union. MJD is supported by a Royal Society Wolfson Fellowship (RSWF\FT\180003). We thank the *Anopheles gambiae* 1000 genomes project for carrying out the sequencing, quality control, SNP calling and haplotype phasing the sequencing data, and Luciene Salas Jennings and Andrew Carey for providing administrative support to the project.

## Funding

Medical Research Council, Grant/Award. Number: MR/P02520X/1; National Institute of Allergy and Infectious Diseases, Grant/Award Number: R01AI116811; Royal Society, Grant/Award Number: RSWF\FT\180003

## Notes

https://github.com/vigg-lstm/GAARD_work

https://www.malariagen.net/data

### Competing Interest Statement

The authors have declared no competing interest.

## References

Adolfi, A., Poulton, B., Anthousi, A., Macilwee, S., Ranson, H., & Lycett, G. J. (2019). Functional genetic validation of key genes conferring insecticide resistance in the major *African malaria vector, Anopheles gambiae*. Proceedings of the National Academy of Sciences of the United States of America, 116(51), 25764–25772.

Anopheles gambiae 1000 Genomes Consortium. (2017). Genetic diversity of the African malaria vector *Anopheles gambiae*. Nature, 552(7683), 96–100.

Bass, C., Denholm, I., Williamson, M. S., & Nauen, R. (2015). The global status of insect *resistance to neonicotinoid insecticides*. Pesticide Biochemistry and Physiology, 121, 78–87.

Bhatt, S., Weiss, D. J., Cameron, E., Bisanzio, D., Mappin, B., Dalrymple, U., Battle, K. E., Moyes, C. L., Henry, A., Eckhoff, P. A., Wenger, E. A., Briët, O., Penny, M. A., Smith, T. A., Bennett, A., Yukich, J., Eisele, T. P., Griffin, J. T., Fergus, C. A., … Gething, P. W. (2015). The effect of malaria control on *Plasmodium falciparum* in Africa between 2000 and 2015. Nature, 526(7572), 207–211.

Blandin, S., Shiao, S.-H., Moita, L. F., Janse, C. J., Waters, A. P., Kafatos, F. C., & Levashina, E. A. (2004). Complement-like protein TEP1 is a determinant of vectorial capacity in the malaria vector Anopheles gambiae. Cell, 116(5), 661–670.

Chabi, J., Van’t Hof, A., N’dri, L. K., Datsomor, A., Okyere, D., Njoroge, H., Pipini, D., Hadi, M. P., de Souza, D. K., Suzuki, T., Dadzie, S. K., & Jamet, H. P. (2019). Rapid high throughput SYBR green assay for identifying the malaria vectors Anopheles arabiensis, Anopheles coluzzii and Anopheles gambiae s.s. Giles. PloS One, 14(4), e0215669.

Choi, H. W., Breman, J. G., Teutsch, S. M., Liu, S., Hightower, A. W., & Sexton, J. D. (1995). The effectiveness of insecticide-impregnated bed nets in reducing cases of malaria *infection: a meta-analysis of published results*. The American Journal of Tropical Medicine and Hygiene, 52(5), 377–382.

Cingolani, P., Platts, A., Wang, L. L., Coon, M., Nguyen, T., Wang, L., Land, S. J., Lu, X., & Ruden, D. M. (2012). A program for annotating and predicting the effects of single nucleotide polymorphisms, SnpEff: SNPs in the genome of Drosophila melanogaster strain w1118; iso-2; iso-3. Fly, 6(2), 80–92.

Cirimotich, C. M., Dong, Y., Garver, L. S., Sim, S., & Dimopoulos, G. (2010). Mosquito *immune defenses against Plasmodium infection*. Developmental and Comparative Immunology, 34(4), 387–395.

Clarkson, C. S., Miles, A., Harding, N. J., O’Reilly, A. O., Weetman, D., Kwiatkowski, D., Donnelly, M. J., & Anopheles gambiae 1000 Genomes Consortium. (2021). The genetic architecture of target-site resistance to pyrethroid insecticides in the African malaria vectors Anopheles gambiae and Anopheles coluzzii. Molecular Ecology, 30(21), 5303–5317.

Clarkson, C. S., Weetman, D., Essandoh, J., Yawson, A. E., Maslen, G., Manske, M., Field, S. G., Webster, M., Antão, T., MacInnis, B., Kwiatkowski, D., & Donnelly, M. J. (2014). Adaptive introgression between *Anopheles* sibling species eliminates a major genomic island but not reproductive isolation. Nature Communications, 5, 4248.

Donnelly, M. J., Isaacs, A. T., & Weetman, D. (2016). Identification, Validation, and Application of Molecular Diagnostics for Insecticide Resistance in Malaria Vectors. Trends in Parasitology, 32(3), 197–206.

Enayati, A. A., Ranson, H., & Hemingway, J. (2005). Insect glutathione transferases and insecticide resistance. Insect Molecular Biology, 14(1), 3–8.

Essandoh, J., Yawson, A. E., & Weetman, D. (2013). Acetylcholinesterase (Ace-1) target site mutation 119S is strongly diagnostic of carbamate and organophosphate resistance in *Anopheles gambiae ss and Anopheles coluzzii across southern Ghana*. Malaria Journal, 12(1), 1–10.

Garud, N. R., Messer, P. W., Buzbas, E. O., & Petrov, D. A. (2015). Recent selective sweeps *in North American Drosophila melanogaster show signatures of soft sweeps*. PLoS Genetics, 11(2), e1005004.

Grau-Bové, X., Lucas, E., Pipini, D., Rippon, E., van ‘t Hof, A. E., Constant, E., Dadzie, S., Egyir-Yawson, A., Essandoh, J., Chabi, J., Djogbénou, L., Harding, N. J., Miles, A., Kwiatkowski, D., Donnelly, M. J., Weetman, D., & Anopheles gambiae 1000 Genomes Consortium. (2021). Resistance to pirimiphos-methyl in West African Anopheles is spreading via duplication and introgression of the Ace1 locus. PLoS Genetics, 17(1), e1009253.

Grau-Bové, X., Tomlinson, S., O’Reilly, A. O., Harding, N. J., Miles, A., Kwiatkowski, D., Donnelly, M. J., Weetman, D., & 1000 Genomes Consortium, A. G. (2020). Evolution of the insecticide target Rdl in African Anopheles is driven by interspecific and interkaryotypic introgression. Molecular Biology and Evolution, 37(10), 2900–2917.

Grigoraki, L., Cowlishaw, R., Nolan, T., Donnelly, M., Lycett, G., & Ranson, H. (2021). CRISPR/Cas9 modified An. gambiae carrying kdr mutation L1014F functionally validate its contribution in insecticide resistance and combined effect with metabolic enzymes. PLoS Genetics, 17(7), e1009556.

Hancock, P. A., Hendriks, C. J. M., Tangena, J.-A., Gibson, H., Hemingway, J., Coleman, M., Gething, P. W., Cameron, E., Bhatt, S., & Moyes, C. L. (2020). Mapping trends in insecticide resistance phenotypes in African malaria vectors. PLoS Biology, 18(6), e3000633.

Hancock, P. A., Lynd, A., Wiebe, A., Devine, M., Essandoh, J., Wat’senga, F., Manzambi, E. Z., Agossa, F., Donnelly, M. J., Weetman, D., & Moyes, C. L. (2022). Modelling spatiotemporal trends in the frequency of genetic mutations conferring insecticide target-site resistance in African mosquito malaria vector species. BMC Biology, 20(1), 46.

Hanghøj, K., Moltke, I., Andersen, P. A., Manica, A., & Korneliussen, T. S. (2019). Fast and accurate relatedness estimation from high-throughput sequencing data in the presence of inbreeding. GigaScience, 8(5). https://doi.org/10.1093/gigascience/giz034

Hearn, J., Djoko Tagne, C. S., Ibrahim, S. S., Tene-Fossog, B., Mugenzi, L. M. J., Irving, H., Riveron, J. M., Weedall, G. D., & Wondji, C. S. (2022). Multi-omics analysis identifies a CYP9K1 haplotype conferring pyrethroid resistance in the malaria vector Anopheles funestus in East Africa. Molecular Ecology, 31(13), 3642–3657.

Ibrahim, S. S., Muhammad, A., Hearn, J., Weedall, G. D., Nagi, S. C., Mukhtar, M. M., Fadel, A. N., Mugenzi, L. J., Patterson, E. I., Irving, H., & Wondji, C. S. (2022). Molecular drivers of insecticide resistance in the Sahelo-Sudanian populations of a major malaria vector. In bioRxiv (p. 2022.03.21.485146). https://doi.org/10.1101/2022.03.21.485146

Ingham, V. A., Tennessen, J. A., Lucas, E. R., Elg, S., Yates, H. C., Carson, J., Guelbeogo, W. M., Sagnon, N. ‘fale, Hughes, G. L., Heinz, E., Neafsey, D. E., & Ranson, H. (2021). Integration of whole genome sequencing and transcriptomics reveals a complex picture of the reestablishment of insecticide resistance in the major malaria vector Anopheles coluzzii. PLoS Genetics, 17(12), e1009970.

Jun, G., Flickinger, M., Hetrick, K. N., Romm, J. M., Doheny, K. F., Abecasis, G. R., Boehnke, M., & Kang, H. M. (2012). Detecting and estimating contamination of human *DNA samples in sequencing and array-based genotype data*. American Journal of Human Genetics, 91(5), 839–848.

Klaus, B., & Strimmer, K. (2015). fdrtool: Estimation of (local) false discovery rates and higher Criticism. http://CRAN.R-project.org/package=fdrtool

Li, C., Zhang, Y., Yun, X., Wang, Y., Sang, M., Liu, X., Hu, X., & Li, B. (2014). Methuselah-like genes affect development, stress resistance, lifespan and reproduction in Tribolium castaneum. Insect Molecular Biology, 23(5), 587–597.

Lindsay, S. W., Thomas, M. B., & Kleinschmidt, I. (2021). Threats to the effectiveness of insecticide-treated bednets for malaria control: thinking beyond insecticide resistance. The Lancet. Global Health, 9(9), e1325–e1331.

Lin, Y. J., Seroude, L., & Benzer, S. (1998). Extended life-span and stress resistance in the Drosophila mutant methuselah. Science, 282(5390), 943–946.

Lucas, E. R., Miles, A., Harding, N. J., Clarkson, C. S., Lawniczak, M. K. N., Kwiatkowski, D. P., Weetman, D., Donnelly, M. J., & The Anopheles gambiae 1000 Genomes Consortium. (2019). Whole genome sequencing reveals high complexity of copy *number variation at insecticide resistance loci in malaria mosquitoes*. Genome Research, 29, 1250–1261.

Lucas, E. R., Rockett, K. A., Lynd, A., Essandoh, J., Grisales, N., Kemei, B., Njoroge, H., Hubbart, C., Rippon, E. J., Morgan, J., Van’t Hof, A. E., Ochomo, E. O., Kwiatkowski, D. P., Weetman, D., & Donnelly, M. J. (2019). A high throughput multi-locus insecticide *resistance marker panel for tracking resistance emergence and spread in Anopheles gambiae*. Scientific Reports, 9, 13335.

Lu, J., Breitwieser, F. P., Thielen, P., & Salzberg, S. L. (2017). Bracken: estimating species abundance in metagenomics data. PeerJ Computer Science. https://peerj.com/articles/cs-104/

Lycett, G. J., McLaughlin, L. A., Ranson, H., Hemingway, J., Kafatos, F. C., Loukeris, T. G., & Paine, M. J. I. (2006). Anopheles gambiae P450 reductase is highly expressed in *oenocytes and in vivo knockdown increases permethrin susceptibility*. Insect Molecular Biology, 15(3), 321–327.

Main, B. J., Lee, Y., Collier, T. C., Norris, L. C., Brisco, K., Fofana, A., Cornel, A. J., & Lanzaro, G. C. (2015). Complex genome evolution in *Anopheles coluzzii* associated with increased insecticide usage in Mali. Molecular Ecology, 24(20), 5145–5157.

Manichaikul, A., Mychaleckyj, J. C., Rich, S. S., Daly, K., Sale, M., & Chen, W. (2010). Robust relationship inference in genome-wide association studies. Bioinformatics, 26(22), 2867–2873.

Martinez-Torres, D., Chandre, F., Williamson, M. S., Darriet, F., Berge, J. B., Devonshire, A. L., Guillet, P., Pasteur, N., & Pauron, D. (1998). Molecular characterization of pyrethroid *knockdown resistance* (*kdr*) in the major malaria vector *Anopheles gambiae* ss. Insect Molecular Biology, 7(2), 179–184.

Mavridis, K., Wipf, N., Medves, S., Erquiaga, I., Müller, P., & Vontas, J. (2019). Rapid multiplex gene expression assays for monitoring metabolic resistance in the major malaria vector Anopheles gambiae. Parasites & Vectors, 12(1), 9.

Mitchell, S. N., Rigden, D. J., Dowd, A. J., Lu, F., Wilding, C. S., Weetman, D., Dadzie, S., Jenkins, A. M., Regna, K., Boko, P., Djogbenou, L., Muskavitch, M. A. T., Ranson, H., Paine, M. J. I., Mayans, O., & Donnelly, M. J. (2014). Metabolic and target-site *mechanisms combine to confer strong DDT resistance* in *Anopheles gambiae*. PloS One, 9(3), e92662.

Mugenzi, L. M. J., Menze, B. D., Tchouakui, M., Wondji, M. J., Irving, H., Tchoupo, M., Hearn, J., Weedall, G. D., Riveron, J. M., & Wondji, C. S. (2019). Cis-regulatory CYP6P9b P450 variants associated with loss of insecticide-treated bed net efficacy against Anopheles funestus. Nature Communications, 10(1), 4652.

Mulamba, C., Riveron, J. M., Ibrahim, S. S., Irving, H., Barnes, K. G., Mukwaya, L. G., Birungi, J., & Wondji, C. S. (2014). Widespread pyrethroid and DDT resistance in the major malaria vector *Anopheles funestus* in East Africa is driven by metabolic resistance mechanisms. PloS One, 9(10), e110058.

Njoroge, H., Van’t Hof, A., Oruni, A., Pipini, D., Nagi, S. C., Lynd, A., Lucas, E. R., Tomlinson, S., Grau-Bove, X., McDermott, D., Wat’senga, F. T., Manzambi, E. Z., Agossa, F. R., Mokuba, A., Irish, S., Kabula, B., Mbogo, C., Bargul, J., Paine, M. J. I., … Donnelly, M. J. (2022). Identification of a rapidly-spreading triple mutant for high-level metabolic insecticide resistance in Anopheles gambiae provides a real-time molecular diagnostic for antimalarial intervention deployment. Molecular Ecology, 31(16), 4307–4318.

Oxborough, R. M. (2016). Trends in US President’s Malaria Initiative-funded indoor residual spray coverage and insecticide choice in sub-Saharan Africa (2008-2015): urgent need for affordable, long-lasting insecticides. Malaria Journal, 15(1), 146.

Riveron, J. M., Irving, H., Ndula, M., Barnes, K. G., Ibrahim, S. S., Paine, M. J. I., & Wondji, C. S. (2013). Directionally selected cytochrome P450 alleles are driving the spread of *pyrethroid resistance in the major malaria vector Anopheles funestus*. Proceedings of the National Academy of Sciences, 110(1), 252–257.

Riveron, J. M., Yunta, C., Ibrahim, S. S., Djouaka, R., Irving, H., Menze, B. D., Ismail, H. M., Hemingway, J., Ranson, H., Albert, A., & Wondji, C. S. (2014). A single mutation in the *GSTe2* gene allows tracking of metabolically based insecticide resistance in a major malaria vector. Genome Biology, 15, R27.

Roth, G. A., Abate, D., Abate, K. H., Abay, S. M., Abbafati, C., Abbasi, N., Abbastabar, H., Abd-Allah, F., Abdela, J., Abdelalim, A., & Others. (2018). Global, regional, and national age-sex-specific mortality for 282 causes of death in 195 countries and territories, *1980-2017: a systematic analysis for the Global Burden of Disease Study 2017*. The Lancet, 392(10159), 1736–1788.

Saavedra-Rodriguez, K., Strode, C., Flores, A. E., Garcia-Luna, S., Reyes-Solis, G., Ranson, H., Hemingway, J., & Black, W. C., 4th. (2014). Differential transcription profiles *in Aedes aegypti detoxification genes after temephos selection*. Insect Molecular Biology, 23(2), 199–215.

Saddler, A., Burda, P.-C., & Koella, J. C. (2015). Resisting infection by Plasmodium berghei *increases the sensitivity of the malaria vector Anopheles gambiae to DDT*. Malaria Journal, 14, 134.

Santolamazza, F., Mancini, E., Simard, F., Qi, Y., Tu, Z., & della Torre, A. (2008). Insertion *polymorphisms of SINE200 retrotransposons within speciation islands of Anopheles gambiae molecular forms*. Malaria Journal, 7(1), 163.

The Anopheles Gambiae 1000 Genomes Consortium. (2021). Ag1000G phase 3 CNV data release. https://www.malariagen.net/data/ag1000g-phase3-cnv

The Anopheles gambiae 1000 Genomes Consortium. (2021). Ag1000G phase 3 SNP data release. MalariaGEN, 2021. https://www.malariagen.net/data/ag1000g-phase3-snp

Vontas, J., Grigoraki, L., Morgan, J., Tsakireli, D., Fuseini, G., Segura, L., Niemczura de Carvalho, J., Nguema, R., Weetman, D., Slotman, M. A., & Hemingway, J. (2018). Rapid selection of a pyrethroid metabolic enzyme CYP9K1 by operational malaria control activities. Proceedings of the National Academy of Sciences, 115(18), 4619–4624.

Weedall, G. D., Mugenzi, L. M. J., Menze, B. D., Tchouakui, M., Ibrahim, S. S., Amvongo-Adjia, N., Irving, H., Wondji, M. J., Tchoupo, M., Djouaka, R., Riveron, J. M., & Wondji, C. S. (2019). A cytochrome P450 allele confers pyrethroid resistance on a major *African malaria vector, reducing insecticide-treated bednet efficacy*. Science Translational Medicine, 11(484), eaat7386.

Weetman, D., Djogbenou, L. S., & Lucas, E. (2018). Copy number variation (CNV) and insecticide resistance in mosquitoes: Evolving knowledge or an evolving problem? Current Opinion in Insect Science, 27, 82–88.

Weetman, D., Wilding, C. S., Steen, K., Pinto, J., & Donnelly, M. J. (2012). Gene Flow–Dependent Genomic Divergence between Anopheles gambiae M and S Forms. Molecular Biology and Evolution, 29(1), 279–291.

Williams, J., Cowlishaw, R., Sanou, A., Ranson, H., & Grigoraki, L. (2022). In vivo functional validation of the V402L voltage gated sodium channel mutation in the malaria vector An. gambiae. Pest Management Science, 78(3), 1155–1163.

World Health Organization. (2016). Test procedures for insecticide resistance monitoring in malaria vector mosquitoes.

Yi, X., Liang, Y., Huerta-Sanchez, E., Jin, X., Cuo, Z. X. P., Pool, J. E., Xu, X., Jiang, H., Vinckenbosch, N., Korneliussen, T. S., Zheng, H., Liu, T., He, W., Li, K., Luo, R., Nie, X., Wu, H., Zhao, M., Cao, H., … Wang, J. (2010). Sequencing of 50 human exomes reveals adaptation to high altitude. Science, 329(5987), 75–78.

