## Supplementary Figures for "Genome-wide association studies reveal novel loci associated with pyrethroid and organophosphate resistance in *Anopheles gambiae* s.l."

### Electronic Supplementary Material

### Supplementary figures and tables

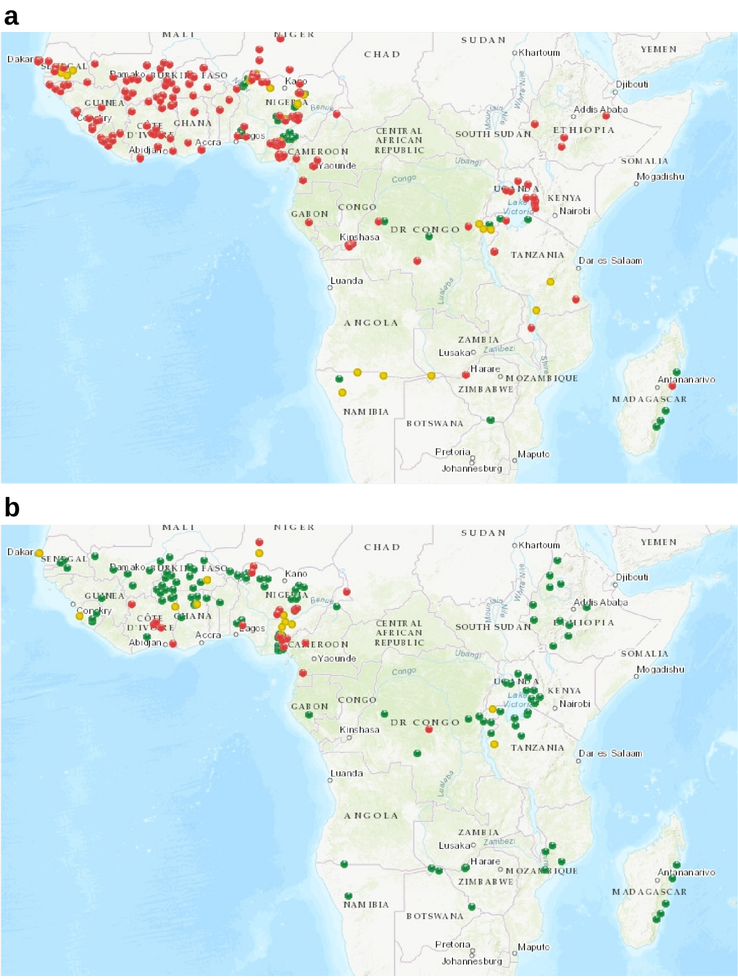


**Fig. S1**: Distribution of insecticide resistance detected by studies in *An. gambiae* sl over the last 5 years (2017-2022) to deltamethrin (a) and PM (b), indicating confirmed resistance (red), possible resistance (yellow) and susceptibility (green). Data obtained from IR-Mapper v2.0 https://anopheles.irmapper.com/ on 25th of November 2022.

| | Ace1_Dup0 | Ace1_Dup1 | Ace1_Dup2 | Ace1_Del1 | Ace1_Del2 | Ace1_Del3 | Ace1_Del4 |
|---|---|---|---|---|---|---|---|
| Avrankou coluzzii | .05 | .05 | 0 | .01 | 0 | 0 | 0 |
| Korle-Bu coluzzii | .85 | .87 | 0 | .44 | 0 | 0 | .06 |
| Aboisso gambiae | .95 | .95 | 0 | .95 | 0 | 0 | 0 |
| Baguida gambiae | .99 | 1 | 0 | .95 | 0 | 0 | 0 |
| Madina gambiae | .9 | .92 | 0 | .77 | 0 | 0 | 0 |
| Obuasi gambiae | 1 | 1 | 0 | .98 | 0 | 0 | 0 |

| | Cyp6aap_Dup0 | Cyp6aap_Dup1 | Cyp6aap_Dup1a | Cyp6aap_Dup1b | Cyp6aap_Dup2 | Cyp6aap_Dup3 | Cyp6aap_Dup4 | Cyp6aap_Dup5 | Cyp6aap_Dup6 | Cyp6aap_Dup7 | Cyp6aap_Dup8 | Cyp6aap_Dup9 | Cyp6aap_Dup10 | Cyp6aap_Dup11 | Cyp6aap_Dup12 | Cyp6aap_Dup13 | Cyp6aap_Dup14 | Cyp6aap_Dup15 | Cyp6aap_Dup16 | Cyp6aap_Dup17 | Cyp6aap_Dup18 | Cyp6aap_Dup19 | Cyp6aap_Dup20 | Cyp6aap_Dup21 | Cyp6aap_Dup22 | Cyp6aap_Dup23 | Cyp6aap_Dup24 | Cyp6aap_Dup25 | Cyp6aap_Dup26 | Cyp6aap_Dup27 | Cyp6aap_Dup28 | Cyp6aap_Dup29 | Cyp6aap_Dup30 |
|---|---|---|---|---|---|---|---|---|---|---|---|---|---|---|---|---|---|---|---|---|---|---|---|---|---|---|---|---|---|---|---|---|---|
| Avrankou coluzzii | .77 | 0 | 0 | .01 | 0 | 0 | 0 | 0 | 0 | .1 | 0 | 0 | .43 | 0 | 0 | 0 | 0 | 0 | 0 | .01 | 0 | 0 | 0 | 0 | 0 | 0 | 0 | 0 | 0 | 0 | 0 | 0 | 0 |
| Korle-Bu coluzzii | .3 | 0 | 0 | 0 | 0 | 0 | 0 | 0 | 0 | .004 | 0 | 0 | .06 | .004 | 0 | 0 | 0 | 0 | 0 | .02 | .16 | 0 | 0 | 0 | 0 | 0 | 0 | 0 | 0 | 0 | 0 | 0 | 0 |
| Aboisso gambiae | .11 | 0 | 0 | 0 | 0 | 0 | 0 | 0 | 0 | 0 | .03 | 0 | 0 | 0 | 0 | 0 | 0 | 0 | 0 | 0 | 0 | .03 | .03 | 0 | 0 | 0 | 0 | 0 | 0 | 0 | 0 | 0 | 0 |
| Baguida gambiae | .03 | 0 | 0 | 0 | 0 | 0 | 0 | 0 | 0 | 0 | 0 | 0 | 0 | 0 | 0 | 0 | 0 | 0 | 0 | 0 | 0 | 0 | 0 | 0 | 0 | 0 | 0 | 0 | 0 | 0 | 0 | 0 | 0 |
| Madina gambiae | .34 | 0 | 0 | 0 | 0 | 0 | 0 | 0 | 0 | 0 | .02 | 0 | 0 | 0 | 0 | 0 | 0 | 0 | 0 | 0 | 0 | 0 | 0 | 0 | 0 | 0 | 0 | 0 | 0 | 0 | 0 | 0 | 0 |
| Obuasi gambiae | .1 | 0 | 0 | 0 | 0 | 0 | 0 | 0 | 0 | 0 | .005 | 0 | 0 | 0 | 0 | 0 | 0 | 0 | 0 | 0 | 0 | 0 | .02 | 0 | 0 | 0 | 0 | 0 | 0 | 0 | 0 | 0 | 0 |

| | Cyp6mz_Dup0 | Cyp6mz_Dupm1 | Cyp6mz_Dupm2 | Cyp6mz_Dupm3 | Cyp6mz_Dupm4 | Cyp6mz_Dupm5 | Cyp6mz_Dupz1 | Cyp6mz_Dupz2 | Cyp6mz_Dupz3 | Cyp6mz_Dupz4 | Cyp6mz_Dupz5 | Cyp6mz_Dupmz1 |
|---|---|---|---|---|---|---|---|---|---|---|---|---|
| Avrankou coluzzii | .04 | 0 | 0 | 0 | 0 | 0 | 0 | 0 | 0 | 0 | 0 | 0 |
| Korle-Bu coluzzii | .06 | 0 | 0 | 0 | 0 | 0 | 0 | 0 | 0 | 0 | 0 | 0 |
| Aboisso gambiae | .05 | 0 | 0 | 0 | 0 | 0 | .03 | 0 | 0 | 0 | 0 | 0 |
| Baguida gambiae | .05 | 0 | 0 | 0 | 0 | 0 | 0 | 0 | 0 | 0 | 0 | 0 |
| Madina gambiae | .24 | 0 | .006 | 0 | 0 | 0 | .11 | 0 | .02 | 0 | 0 | 0 |
| Obuasi gambiae | .2 | 0 | .06 | 0 | 0 | 0 | .09 | 0 | 0 | 0 | 0 | 0 |

| | Gstue_Dup0 | Gstue_Dup1 | Gstue_Dup2 | Gstue_Dup3 | Gstue_Dup4 | Gstue_Dup5 | Gstue_Dup6 | Gstue_Dup7 | Gstue_Dup8 | Gstue_Dup9 | Gstue_Dup10 | Gstue_Dup11 | Gstue_Dup12 | Gstue_Dup13 | Gstue_Dup14 | Gstue_Dup15 | Gstue_Dup16 |
|---|---|---|---|---|---|---|---|---|---|---|---|---|---|---|---|---|---|
| Avrankou coluzzii | .68 | .05 | 0 | 0 | 0 | 0 | 0 | .52 | 0 | 0 | 0 | 0 | 0 | 0 | 0 | 0 | 0 |
| Korle-Bu coluzzii | .23 | 0 | 0 | 0 | 0 | 0 | 0 | .22 | 0 | 0 | 0 | 0 | 0 | 0 | 0 | 0 | 0 |
| Aboisso gambiae | 0 | 0 | 0 | 0 | 0 | 0 | 0 | 0 | 0 | 0 | 0 | 0 | 0 | 0 | 0 | 0 | 0 |
| Baguida gambiae | .01 | 0 | 0 | 0 | 0 | 0 | 0 | 0 | 0 | 0 | 0 | 0 | 0 | 0 | 0 | 0 | 0 |
| Madina gambiae | .07 | 0 | 0 | 0 | 0 | 0 | 0 | 0 | 0 | 0 | 0 | 0 | 0 | 0 | 0 | 0 | 0 |
| Obuasi gambiae | .08 | 0 | 0 | 0 | 0 | 0 | 0 | 0 | 0 | 0 | 0 | 0 | 0 | 0 | 0 | 0 | 0 |

| | Cyp9k1_Dup0 | Cyp9k1_Dup1 | Cyp9k1_Dup2 | Cyp9k1_Dup3 | Cyp9k1_Dup4 | Cyp9k1_Dup5 | Cyp9k1_Dup6 | Cyp9k1_Dup7 | Cyp9k1_Dup8 | Cyp9k1_Dup9 | Cyp9k1_Dup10 | Cyp9k1_Dup11 | Cyp9k1_Dup12 | Cyp9k1_Dup13 | Cyp9k1_Dup14 | Cyp9k1_Dup15 | Cyp9k1_Dup16 | Cyp9k1_Dup17 | Cyp9k1_Dup18 | Cyp9k1_Dup19 | Cyp9k1_Dup20 | Cyp9k1_Dup21 | Cyp9k1_Dup22 | Cyp9k1_Dup23 | Cyp9k1_Dup24 | Cyp9k1_Dup25 | Cyp9k1_Dup26 | Cyp9k1_Dup27 | Cyp9k1_Dup28 |
|---|---|---|---|---|---|---|---|---|---|---|---|---|---|---|---|---|---|---|---|---|---|---|---|---|---|---|---|---|---|
| Avrankou coluzzii | .03 | 0 | 0 | 0 | 0 | 0 | 0 | 0 | 0 | 0 | .03 | 0 | 0 | 0 | 0 | 0 | 0 | 0 | 0 | 0 | 0 | 0 | 0 | 0 | 0 | 0 | 0 | 0 | 0 |
| Korle-Bu coluzzii | .16 | 0 | 0 | 0 | 0 | 0 | 0 | 0 | 0 | 0 | .17 | 0 | 0 | 0 | 0 | 0 | 0 | 0 | 0 | 0 | 0 | 0 | 0 | 0 | 0 | 0 | 0 | 0 | 0 |
| Aboisso gambiae | .84 | 0 | 0 | 0 | 0 | 0 | 0 | 0 | 0 | 0 | 0 | .16 | 0 | 0 | 0 | 0 | 0 | 0 | 0 | 0 | 0 | 0 | 0 | 0 | 0 | 0 | 0 | 0 | .73 |
| Baguida gambiae | .96 | 0 | 0 | 0 | .16 | 0 | 0 | .16 | 0 | 0 | .73 | .01 | 0 | 0 | 0 | 0 | 0 | 0 | 0 | 0 | 0 | 0 | .02 | 0 | 0 | 0 | 0 | 0 | .19 |
| Madina gambiae | .93 | 0 | 0 | 0 | .1 | 0 | 0 | .006 | 0 | 0 | .006 | .13 | 0 | .006 | 0 | 0 | 0 | 0 | 0 | 0 | 0 | 0 | .006 | 0 | 0 | 0 | 0 | 0 | .86 |
| Obuasi gambiae | .93 | 0 | 0 | 0 | .02 | 0 | 0 | .02 | 0 | 0 | 0 | .02 | 0 | 0 | 0 | 0 | 0 | 0 | 0 | 0 | 0 | 0 | .01 | 0 | 0 | 0 | 0 | 0 | .91 |

**Fig. S2**: Frequency (proportion of samples carrying at least one copy) of known CNV alleles detected using diagnostic reads around *Ace1*, the *Cyp6aa / Cyp6p* cluster, the *Cyp6m / Cyp6z* cluster, the *Gste* cluster and *Cyp9k1*. Darkness of blue (*An. gambiae*) and red (*An. coluzzii*) provided as a visual aid for the magnitude of the value in each cell. The genomic coordinates of each CNV allele can be found in Supplementary Data @@. In each cluster, the "Dup0" column indicates the presence of increased copy number in any of the genes in the cluster. Where this is larger than the sum of known alleles, it suggests the presence of CNV alleles not present in the Ag1000G database. "Del" alleles in Ace1 represent secondary deletions within the Ace1-Dup1 CNV.

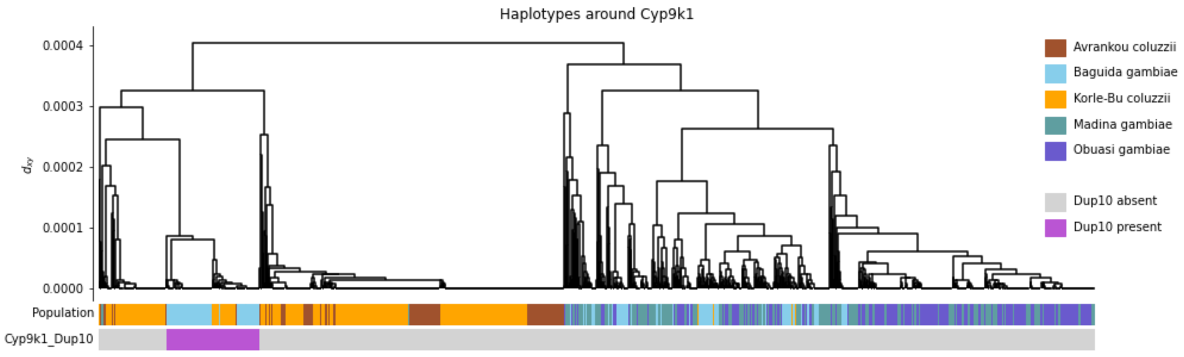


**Fig. S3**: Haplotypes bearing the CNV allele Cyp9k1_Dup10 in *An. gambiae* and *An. coluzzii* form a single cluster nested within other *An. coluzzii* haplotypes, indicating that the haplotype is of *An. coluzzii* origin and thus that introgession of the allele was from *An. coluzzii* to *An. gambiae*. Haplotype clustering was performed using 500 SNPs around *Cyp9k1*. The Cyp9k1_Dup10 CNV allele was phased by identifying SNPs perfectly correlated with its presence / absence level. Full workings to reproduce this analysis can be found at https://github.com/vigg-lstm/GAARD_work/blob/main/CNV_analysis/Cyp9k1_Dup10_haplotypes/Cyp9k1_CNV_haplotypes.ipynb.

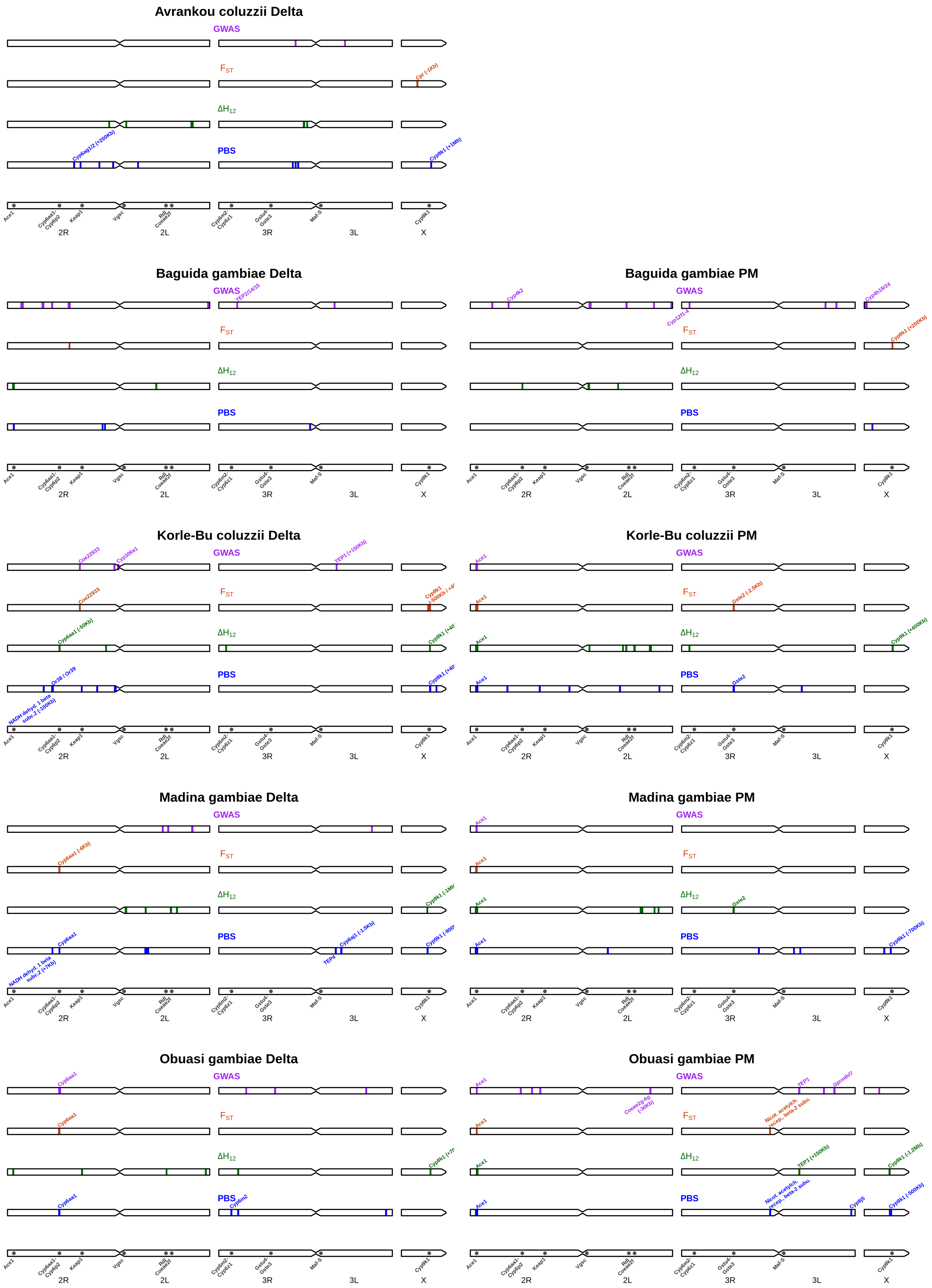


**Fig. S4**: Genomic regions implicated in insecticide resistance by each of our four approaches. For the global GWAS method, these are 100,000 bp windows which contained at least 10 of the top 1000 significant SNPs. For $F_{ST}$, these are significant peaks which contained at least one haplotype significantly positively associated with resistance (Supplementary Data S2). For $\Delta H_{12}$ and PBS, these are significant positive peaks (ie: indicating stronger signals of selection in resistant compared to susceptible samples). Regions are annotated with genes discussed in the manuscript as possibly causing the signal. Genomic distances in brackets indicate the distance of the peak either to the left (-) or right (+) of the gene in question.

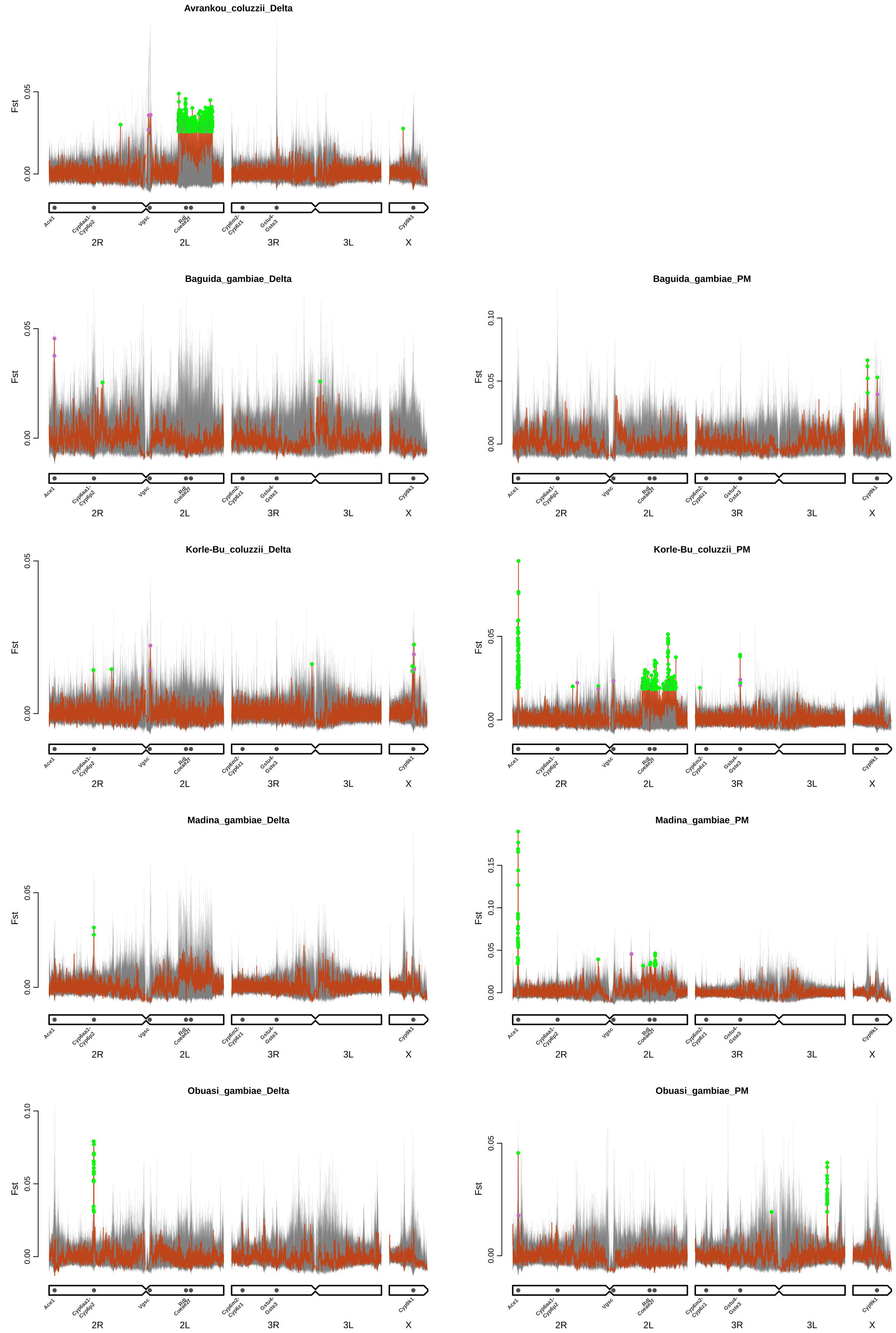


**Fig. S5**: $F_{ST}$ between resistant and susceptible individuals in each sample set, calculated in 1000 SNP windows. Red line indicates $F_{ST}$, grey lines in background show results from 200 randomisations in which phenotype labels were permuted. Regions of extended haplotype homozygosity can cause spurious peaks in $F_{ST}$, which are captured by the randomisations (eg., peaks around Ace1 in Deltamethrin sample sets). Windows identified as peaks were considered "significant" (green points) if their $F_{ST}$ value fell above the 99th centile of the randomisations for that window (ie: $P < 0.01$) and non-significant (purple points) otherwise.

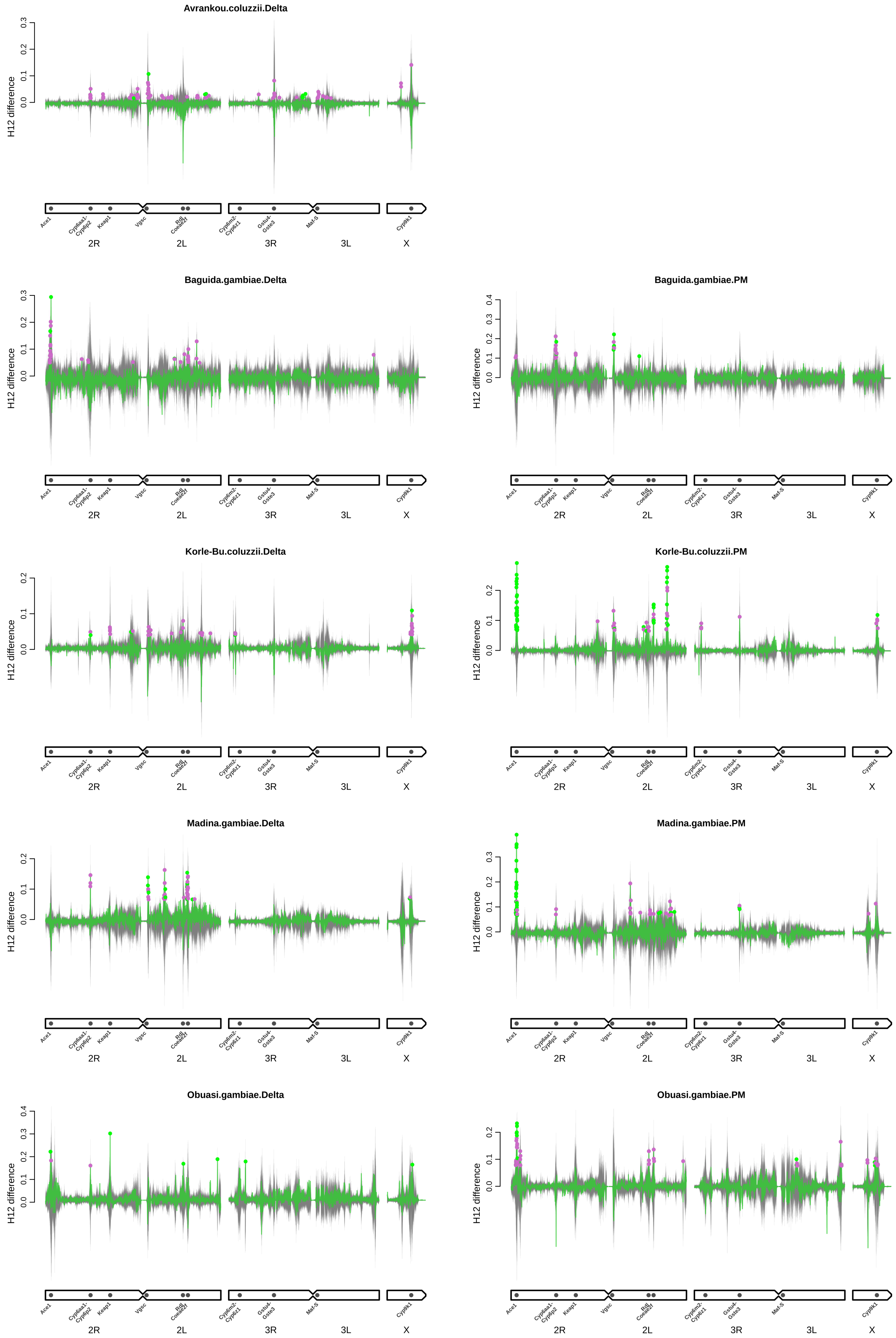


**Fig. S6**: Difference in $H_{12}$ between resistant and susceptible sub-samples ($\Delta H_{12}$) in each sample set, calculated in 1000 SNP windows. Green line indicates $\Delta H_{12}$, grey lines in background show results from 200 randomisations in which phenotype labels were permuted. Regions of extended haplotype homozygosity can cause spurious peaks in $\Delta H_{12}$, which are captured by the randomisations. Windows identified as positive peaks were considered "significant" (green points) if their $\Delta H_{12}$ value fell above the 99th centile of the randomisations for that window (ie: $P < 0.01$) and non-significant (purple points) otherwise.

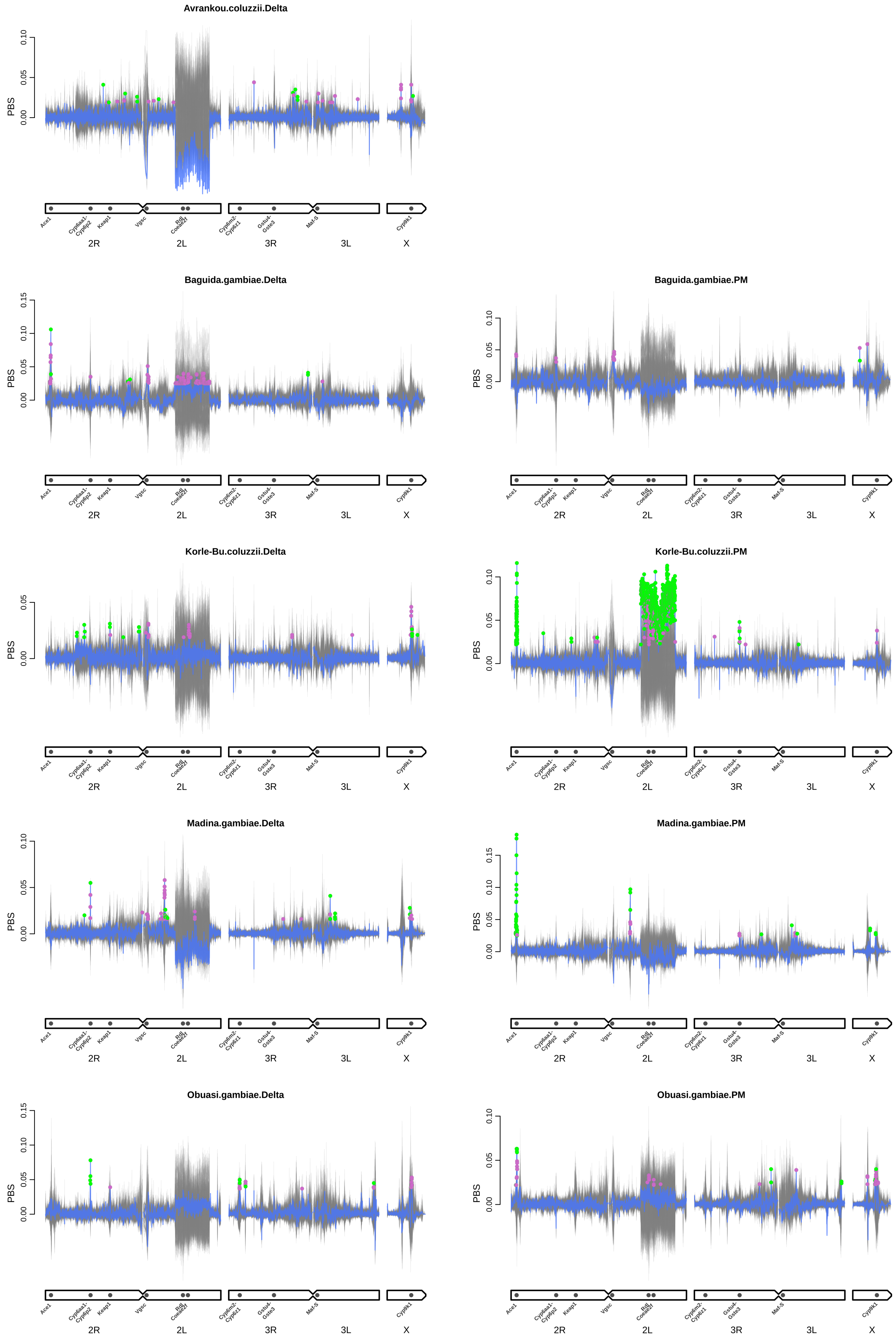


**Fig. S7**: PBS between resistant and susceptible individuals in each sample set, calculated in 1000 SNP windows. Orange line indicates PBS, grey lines in background show results from 200 randomisations in which phenotype labels were permuted. Regions of extended haplotype homozygosity can cause spurious peaks in PBS which are captured by the randomisations. Windows identified as positive peaks were considered "significant" (green points) if their PBS value fell above the 99th centile of the randomisations for that window (ie: $P < 0.01$) and non-significant (purple points) otherwise.

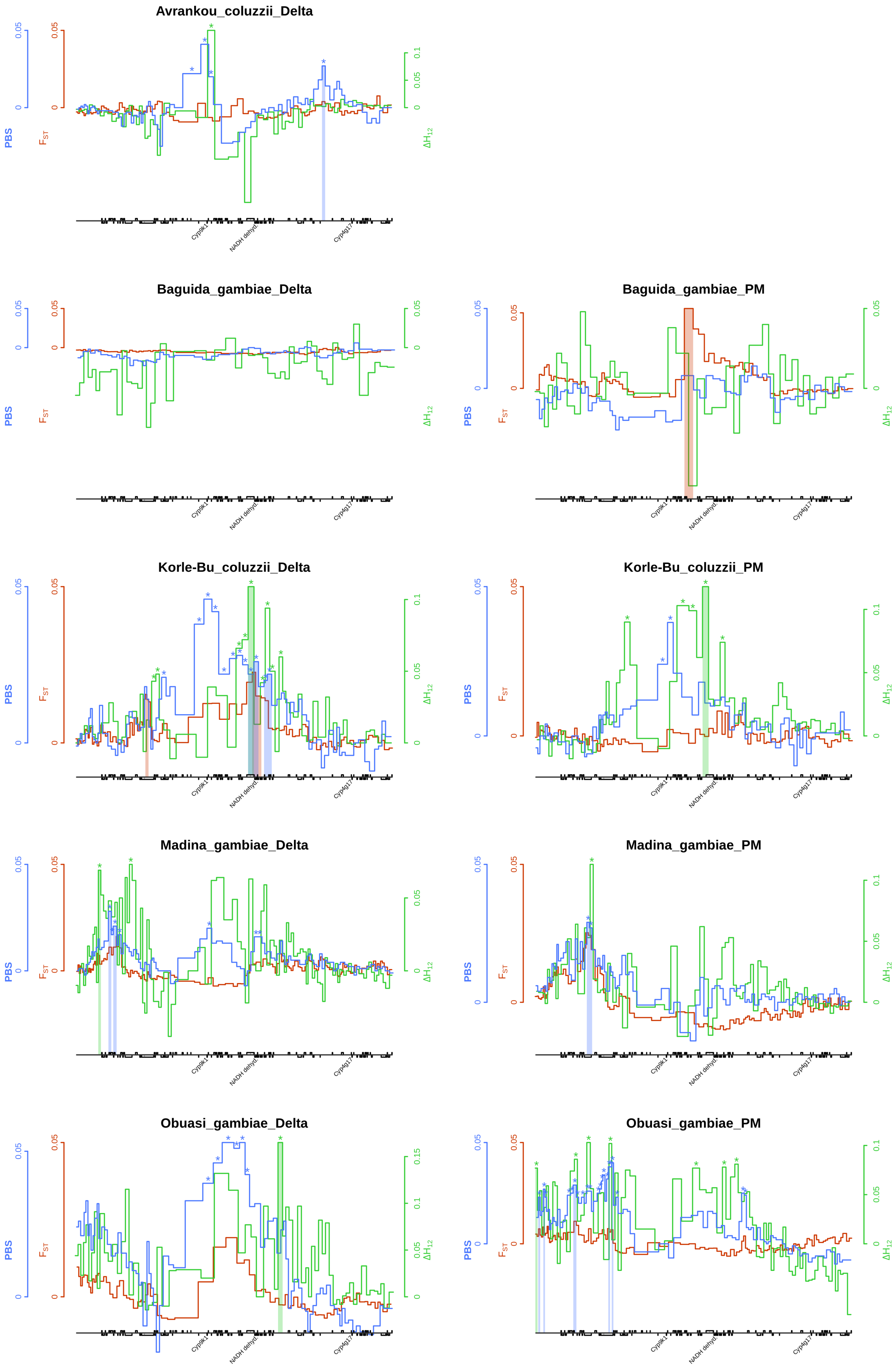


**Fig. S8**: Caption on next page

**Fig. S8** caption: Genomic windows of phenotypic association around *Cyp9k1* were never at the *Cyp9k1* locus itself. Plot shows $F_{ST}$ (red), $\Delta H_{12}$ (green) and PBS (blue), with shaded rectangles extending to the bead plot below indicating peaks that were significantly associated with resistance phenotype based on phenotype randomisations (there are the peaks summarised genome-wide in figure S4). Asterisks denote $\Delta H_{12}$ and PBS peaks determined by outlier analysis, with many being non-significant according to phenotype randomisations. These peaks may be caused by the presence of a selective sweep in the region, regardless of whether the sweep is associated with the phenotype. The positions of *Cyp9k1*, NADH dehydrogenase (ubquinone) 1 $\beta$ subcomponent 1 ("NADH dehyd.") and *Cyp4g17* (discussed in the main text) are shown on the bead plot.
