## Supplementary Methods for "Genome-wide association studies reveal novel loci associated with pyrethroid and organophosphate resistance in *Anopheles gambiae* s.l."

#### Electronic Supplementary Material

#### Supplementary methods

### 1 Sample collection and phenotyping

Mosquitoes were collected in 2017 from six locations in West Africa (Benin: Avrankou [6.550, 2.667], Côte d'Ivoire: Aboisso [5.467, -3.200], Ghana: Madina [5.683, -0.166], Korle-Bu [5.537, -0.240] and Obuasi [6.200, -1.683]; Togo: Baguida [6.161, 1.314]). Larvae were collected by dipping from multiple habitats, pooled, and raised to adults in the laboratory. Non-blood-fed adult females were separated and at 3-5 days old bioassayed using WHO tubes in replicates of 25 with either deltamethrin or pirimiphos methyl (PM) papers. Initially, to determine appropriate doses to produce well-separated phenotypic groups, which eliminated those of intermediate phenotype (Fig. **M1**), 1-2 replicate tubes were run (for 60 min, with mortality assessed 24h later) at a range of concentrations reflecting X-fold the standard WHO diagnostic dose of 0.05% for deltamethrin (0.5X, 1X, 2X, 5X and 10X) and of 0.25% for PM (0.5X, 1X, 2X). From these preliminary data appropriate "lower" and "higher" doses (Fig. **M1**) were determined to achieve the desired phenotypic separation. Our initial plan to use different concentrations with a fixed (60 min) time was followed in all cases with the exception of Obuasi and Baguida phenotyped for PM, for which survivorship with 1X papers at 60 min was very low ($\leq$ 2%, indicating susceptibility according to the WHO criterion). In these cases we obtained separated phenotypic groups by using a single concentration (0.5X) but varying exposure times. Bioassay doses used are shown in Table M1, and results from bioassays in Fig. **M2**. Note that with the exception of PM phenotypes in the two sites above, all populations tested conform to WHO defined resistance (Fig. **M2**, Table M1), and in most cases for deltamethrin resistance would be classified as substantial ($<$90% at *geq* 5X).

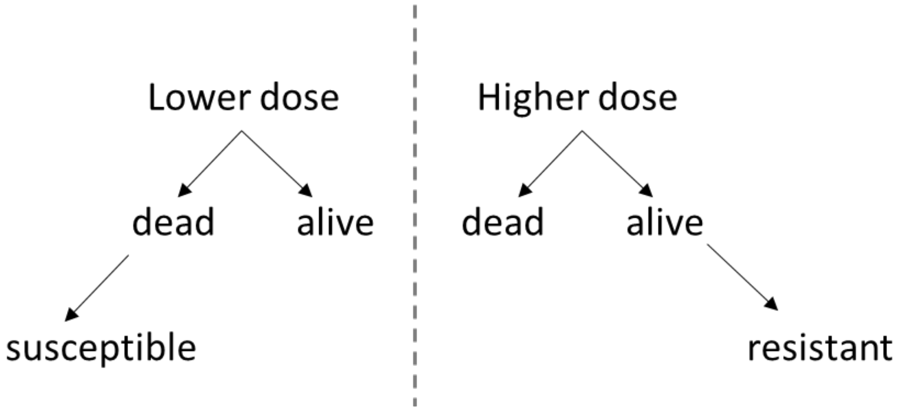


**Fig. M1**: Schematic diagram illustrating the bioassay strategy to obtain well-separated susceptible and resistant phenotypic groups for whole genome sequencing. Our initial target was to use a lower dose resulting in 20% mortality, and a higher dose resulting in 80% mortality with N=100 samples preserved from each. Though this level of separation was seldom feasible, good separation was generally achieved (see Fig. **M2**).

DNA was extracted from samples within each phenotype group using nexttec kits (Biotechnologie GmbH) according to manufacturer's instructions. Species identity of sam-

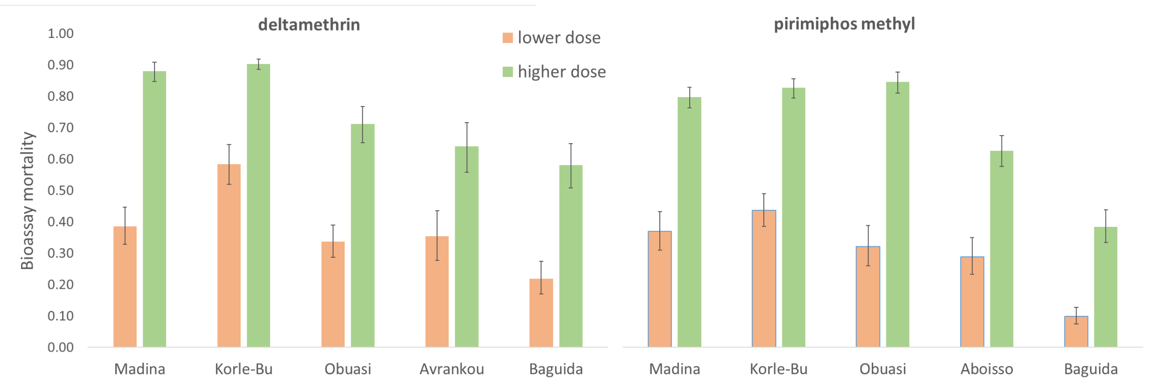


**Fig. M2**: Mean bioassay mortalities (with 95% binomial confidence intervals) of *An. gambiae* s.l. for the lower and higher doses obtained for each insecticide in the different West African sites sampled. Doses used for each site are shown in Table M1, with details of the sibling species composition within each phenotype group in each site.

**Table M1**: Bioassays to segregate females defined as susceptible (dead at lower dose) from resistant (alive at higher dose). All bioassays were performed on *An. gambiae* s.l. with subsequent molecular identification of species within the tested samples as shown. Numbers highlighted in bold show the groups chosen for sequencing. Final numbers for each sample set, after QC filtering of sequencing data, are shown in Table 1 (main text). Delta = deltmethrin; PM = pirimiphos-methyl.

| | | | Lower dose details | | Higher dose details | | *An. gambiae s.l.* | | *An. gambiae* | | *An. coluzzii* | |
|---|---|---|---|---|---|---|---|---|---|---|---|---|
| **Country** | **Site** | **Insecticide** | **Conc.** | **Time (min)** | **Conc.** | **Time (min)** | **Dead N** | **Alive N** | **Dead N** | **Alive N** | **Dead N** | **Alive N** |
| Benin | Avrankou | Delta | 0.5X | 60 | 2X | 60 | 58 | 50 | 1 | 1 | 49 | 57 |
| Ghana | Korle-Bu | Delta | 0.5X | 60 | 5X | 60 | 58 | 43 | 0 | 0 | 43 | 58 |
| | Korle-Bu | PM | 0.5X | 60 | 1X | 60 | 46 | 68 | 0 | 0 | 68 | 46 |
| | Madina | Delta | 1X | 60 | 10X | 60 | 47 | 141 | 76 | 45 | 65 | 2 |
| | Madina | PM | 0.5X | 60 | 1X | 60 | 52 | 91 | 63 | 43 | 28 | 9 |
| | Obuasi | Delta | 2X | 60 | 5X | 60 | 68 | 112 | 111 | 68 | 1 | 0 |
| | Obuasi | PM | 0.5X | 15, 30 | 0.5X | 45, 60 | 71 | 69 | 69 | 71 | 0 | 0 |
| | | | | | | 75, 90 | | | | | 0 | 0 |
| Ivory Coast | Aboisso | PM | 0.5X | 60 | 1X | 60 | 83 | 64 | 13 | 75 | 51 | 8 |
| Togo | Baguida | Delta | 2X | 60 | 10X | 60 | 82 | 57 | 56 | 81 | 1 | 1 |
| | Baguida | PM | 0.5X | 45 | 0.5X | 90 | 59 | 50 | 44 | 58 | 6 | 1 |

ples was determined using either of two methods designed to discriminate between *An. gambiae*, *An. coluzzii* and *An. arabiensis*: a PCR of species-specific SINE insertion polymorphisms as described in Santolamazza et al. 2008, and a high-resolution melt curve analysis Chabi et al. 2019. Results are shown in Table M1. In most cases, samples were dominated by a single species, meaning that bioassay results for the majority species will closely reflect those for *An. gambiae* s.l. (Fig. **M2**). In two sites (Madina and Aboisso) *An. gambiae* and *An. coluzzii* were both present in the collections in substantial proportions and in each case *An. gambiae* were over-represented relative to *An. coluzzii* in resistant compared to susceptible groups (Madina, deltamethrin odds ratio = 19.2; PM odds ratio = 2.1; Aboisso, PM odds ratio = 36.8). This indicates that bioassay mortalities will be overestimated for *An. gambiae* in each case, compared to the results for *An. gambiae* s.l., this will apply to both the lower and higher doses. Therefore, pronounced phenotypic segregation between susceptible and resistant groups should still remain, though potentially quantitatively different from the *An. gambiae* s.l. data shown in Fig. **M2**.

#### 2 Choice of kinship treshold

We calculated pairwise kinship between all samples using the KING statistic (Manichaikul et al. 2010) implemented in NGSRelate (Korneliussen and Moltke 2015) using SNP data across the whole genome. Whole genome SNPs were used because the recombination rate on such a small genome can lead to large disparity in kinship values between chromosomes (Fig. M3). Results indicated a slight positive bias in kinship, with the mode of the distribution slightly above 0 (Fig. M4). Because of this positive bias in kinship values, we sought to empirically establish the most parsimonious threshold to identify full siblings in our data, instead of the threshold of 0.177 suggested in the manual (https://www.kingrelatedness.com/manual.shtml). For all possible threshhold between 0.15 and 0.35, in increments of 0.005, we identified all full siblings and counted the proportion of full sib groups that contained inconsistencies (where siblings of siblings were not themselves classed as siblings). The script to carry out this analysis is available at . We chose the threshold 0.195 as that which produced the smallest proportion of inconsistent sib groups.

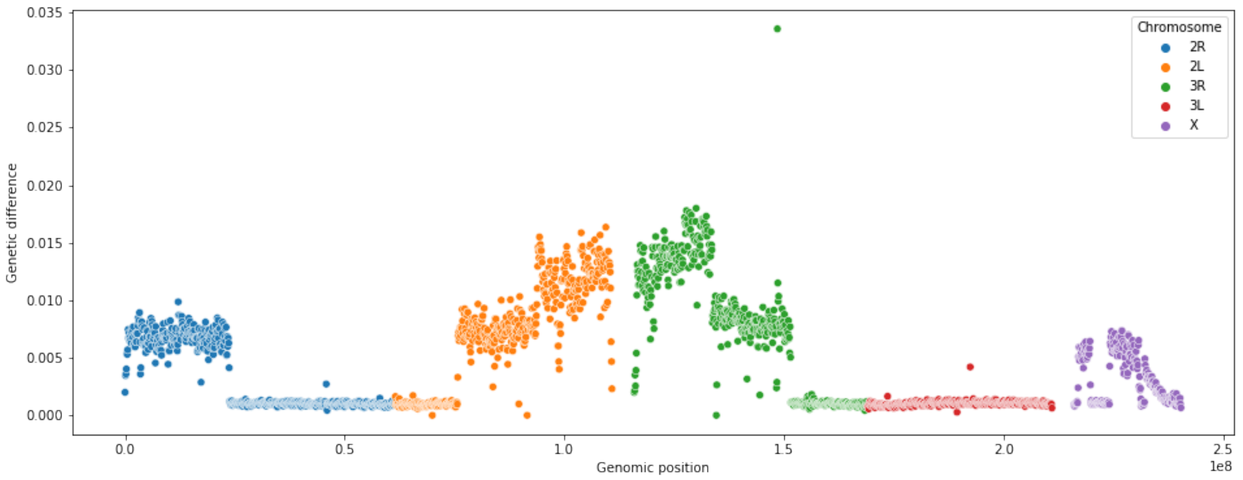


**Fig. M3**: Genetic difference (calculated as the mean proportion of nucleotide differences across all sites) between a single pair of siblings (WA-0997 and WA-1288, both *An. coluzzii* from Korle-Bu in Ghana) across the genome. Aside from the centromeres, the values of genetic difference occupy one of three states: 0 (genetically identical, where the two individuals inherited the same allele from both their mother and their father, for example the start of Chromosome 2L), around 0.007 (intermediate: where the two individuals inherited the same allele from one of their parents, but not the other, for example the middle of chromosome 2L) and around 0.014 (low: where the two individuals inherited different alleles from both their mother and their father, for example the end of chromosome 2L). These states are invariant over large stretches of the genome (eg: the whole of chromosome 3L is identical between these two individuals) due to the recombination rate per site per generation (usually assumed to be $10-8$ for modelling purposes) relative to the size of a chromosome (on the order of $5x10^{-7}$). Thus, a single chromosome can give an inaccurate picture of relatedness (using chromosome 3L for this pair would make them appear to be clones).

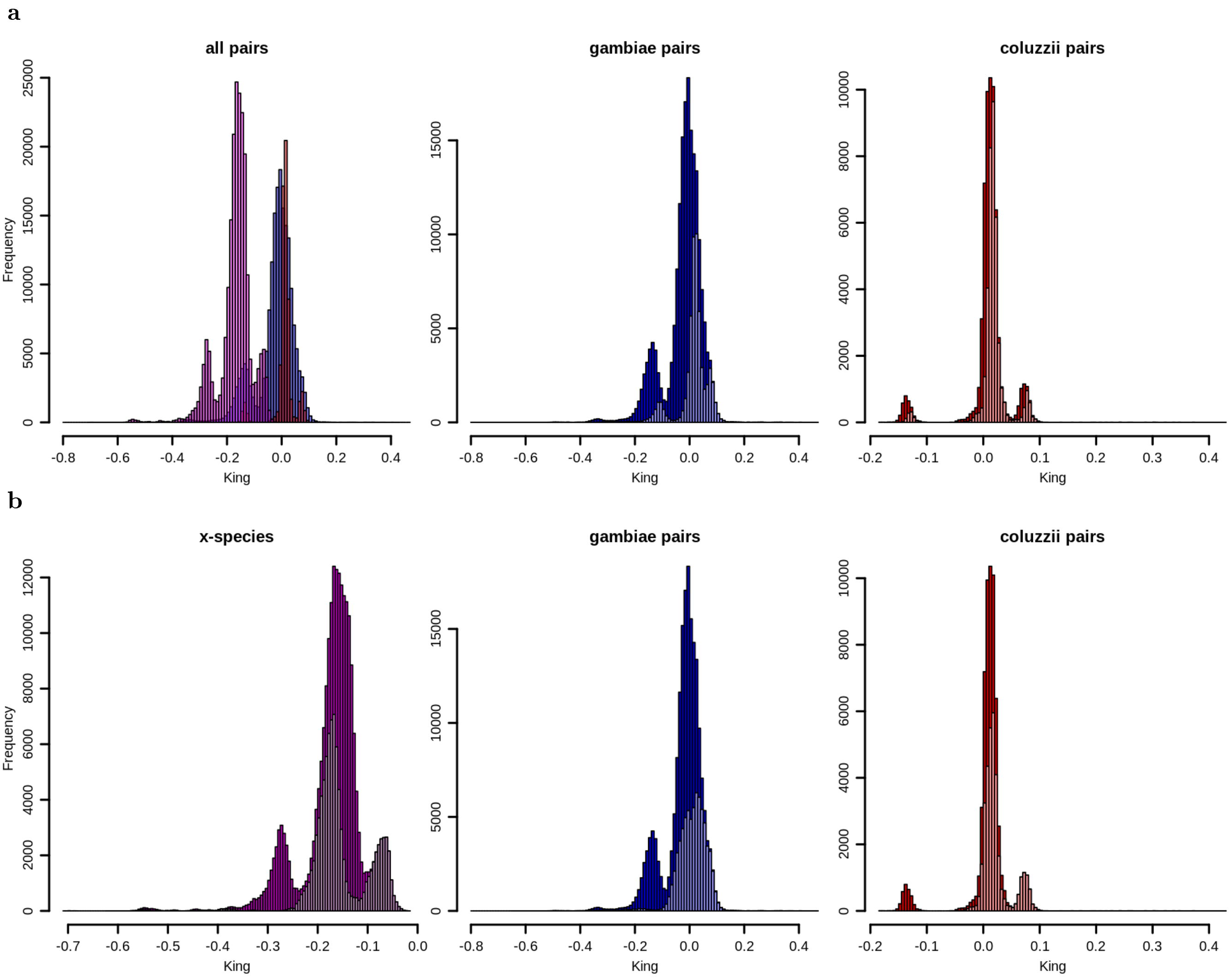


**Fig. M4**: Histograms of kinship values (KING scores, expected score between full sibs = 0.25, expected score between unrelated individuals = 0) across all sample pairs. **a** Left: Coloured by species (blue = gambiae, red = coluzzii, magenta = cross-species pair). Middle and right: stacked and coloured by whether the pair is of individuals from the same (light colours) or different (dark colours) sampling locations. **b** Stacked and coloured by whether the pair is of individuals with the same (light) or different (dark) 2La karyotype. The three modes of the distribution correspond to the 2La karyotype of the pair, with the right hand mode consisting of pairs of 2La heterozygotes (shared heterozygote genotype produces the largest relatedness values), the main central mode consisting of pairs with the same homozygous karyotype (in dark) or where one individual is heterozygous and the other homozygous (in light), and the left hand mode consisting of pairs with opposite homozygous karyotypes.

#### 3 $F_{ST}$ outlier analysis

We calculcated $F_{ST}$ between phenotype classes (dead / alive after exposure to insecticides) in windows of 1000 SNPs across the genome, and wanted to identify peaks in the data. We used the fact that true $F_{ST}$ peaks in these data should only be positive, meaning that the left hand side of the distribution should be largely unaffected by the number of peaks in the data. We therefore used the left hand part of the $F_{ST}$ distribution to determine what the limits of the right hand part would typically look like in the absence of peaks, and identified outliers as positive windows beyond these limits. To do this, we took the difference between the smallest $F_{ST}$ value and the mode of the distribution, and considered an outlier to be any value more than three times this distance away from the mode (Fig. **M5**).

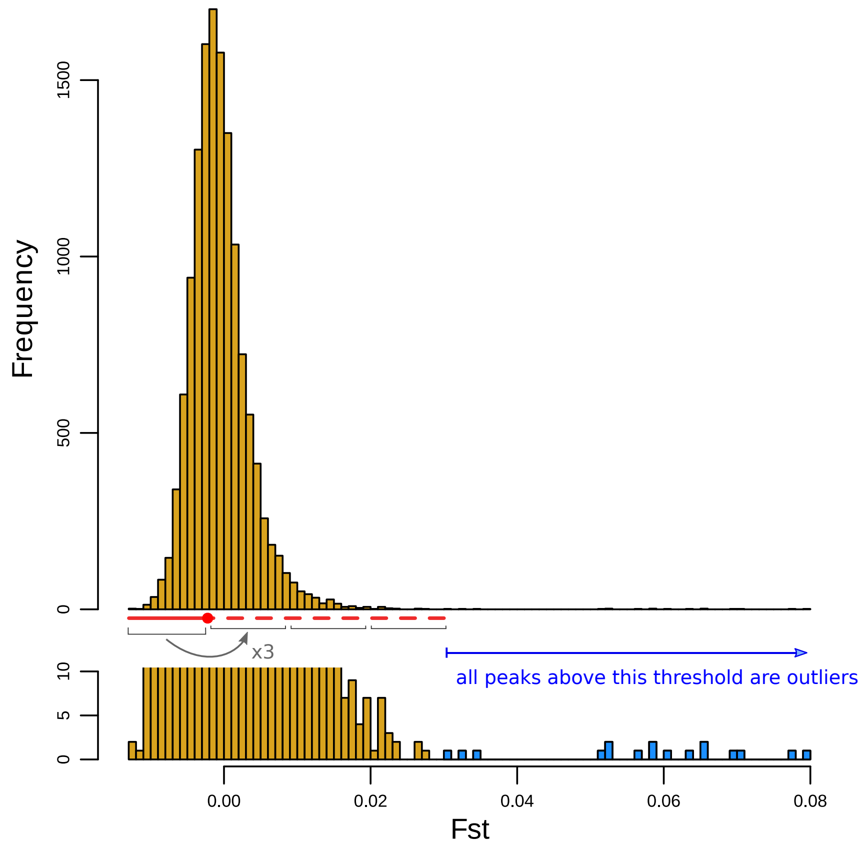


**Fig. M5**: Historgram of $F_{ST}$ values between phenotype groups in*An. gambiae* from Obuasi exposed to deltamethrin (top) and zoomed in on at the origin of the Y axis to show small outlier peaks (bottom). We first identified the mode of the distribution (red point) and then calculated its distance from the minimum of the distribution (solid red line). We then conservatively took three times this distance to the right of the mode (dashed red line) as our threshold. Any windows with $F_{ST}$ values larger than this (in blue) were considered outliers and thus provisional windows of interest.

#### 4 Removal of *Asaia* contamination

In an original pass of the GWAS analysis, we found SNPs significantly correlated with phenotype, but which showed strong allelic imbalance and, in some cases, heterozygote excess. Looking into these SNPs, we found that they were the result of mis-alignment of non-*Anopheles* reads. Assembling and BLASTing the reads in question indicated that they likely originated from *Asaia* / *Acetobacter* bacteria. There are two possible explanations for this. The first is that of a metagenomic association with resistance. The second is of sample contamination differentially affecting the two phenotypes. We investigated this possibility by examining the distribution of the levels of *Asaia* reads with respect to the position of the sample on the plates in which DNA was stored. Fig. M6 shows that *Asaia* reads are non-randomly distributed across plates, indicating that they are likely to be the result of contamination, but creating the appearance of correlation with phenotype.

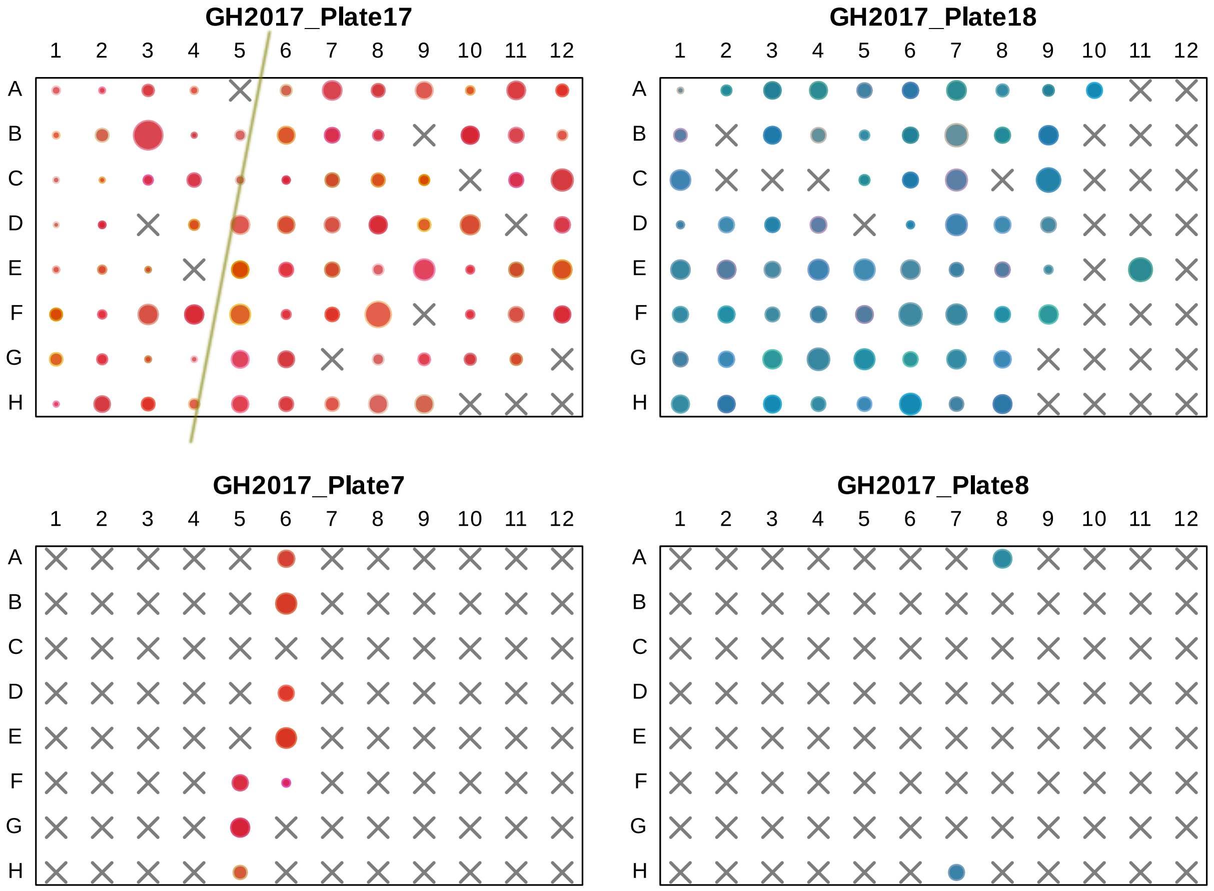


**Fig. M6**: Layout of levels of *Asaia* reads across DNA storage plates for deltamethrin-exposed samples from Korle-Bu. Size of circle indicates normalised number of Asaia reads. Colour indicates phenotype of sample (blue = alive, red = dead). Crosses indicate wells not used for sequencing. Diagonal line on top left plate separates wells with broadly less *Asaia* signal on the left and more signal on the right.

To control for this in the GWAS, we used Bracken (Lu et al. 2017) to estimate the amount of *Asaia* signal in each sample, and then calculated the correlation between *Asaia* levels and each SNP in the analysis. SNPs that were correlated with *Asaia* (uncorrected $P < 0.05$) were removed from the GWAS.

### References


Chabi, J. et al. (2019). Rapid high throughput SYBR green assay for identifying the malaria vectors Anopheles arabiensis, Anopheles coluzzii and Anopheles gambiae s. s. Giles. *PLoS One* 14(4):e0215669.

Korneliussen, T. S. and I. Moltke (2015). NgsRelate: a software tool for estimating pairwise relatedness from next-generation sequencing data. *Bioinformatics* 31(24):4009–4011.

Lu, J. et al. (2017). Bracken: estimating species abundance in metagenomics data. *PeerJ Computer Science* 3:e104.

Manichaikul, A. et al. (2010). Robust relationship inference in genome-wide association studies. *Bioinformatics* 26(22):2867–2873.

Santolamazza, F. et al. (2008). Insertion polymorphisms of SINE200 retrotransposons within speciation islands of *Anopheles gambiae* molecular forms. *Malaria journal* 7(1):163.
