## Supplementary Data S2 for "Genome-wide association studies reveal novel loci associated with pyrethroid and organophosphate resistance in *Anopheles gambiae* s.l.": Supplementary_Data_S2.html

window\_Fst\_haplotype\_summary


### F_ST_ windows of interest

For each sample set, we first provide a summary plot of
F_ST_ bewteen resistant and susceptible samples across the
genome, with F_ST_ shown in red and the results of the 200
randomisations shown behind in grey. Windows identified as peaks are
highlighted by points, colour-coded by whether they are significantly
higher than expected based on the simulations (green) or not (purple).
For each significant window (green points), we then provide four
additional figures. The first shows the significant (*P* <
0.01) SNPs found in the region of that window, and their -log10(Pvalue)
of association with phenotype. Red points indicate non-synonymous SNPs,
blue points indicate all other SNPs. Point shape indicates whether the
mutant allele at that SNP is associated with increased (circle) or
decreased (triangle) resistance. Dark points in the centre of the plot
show SNPs within the significant window, light points on the sides show
SNPs in region 10,000 bp either side of the window. The second plot
shows the hierarchical cluster of haplotypes in the window, with
brackets indicating large haplotype clusters. The third plot shows, each
haplotype cluster, indicating the number of haplotypes comprising the
cluster (n), the *P* value of association of the cluster with
resistance phenotype, and the SNPs that differentiate that cluster from
other haplotypes in the region, colour-coded by type (red =
non-synonymous, yellow = synonymous, blue = non-coding). The final
figure shows the contingency tables of phenotype association for each
haplotype cluster.

Legend  
Avrankou\_*coluzzii*\_Delta  
Baguida\_*gambiae*\_Delta  
Korle-Bu\_*coluzzii*\_Delta  
Madina\_*gambiae*\_Delta  
Obuasi\_*gambiae*\_Delta  
Baguida\_*gambiae*\_PM  
Korle-Bu\_*coluzzii*\_PM  
Madina\_*gambiae*\_PM  
Obuasi\_*gambiae*\_PM

---

#### Plot legend

---

#### Avrankou\_*coluzzii*\_Delta

Avrankou\_coluzzii\_Delta\_2R:45413683-45433265  
Avrankou\_coluzzii\_Delta\_X:8745589-8778998

##### Avrankou\_coluzzii\_Delta\_2R\_45413683-45433265

##### Avrankou\_coluzzii\_Delta\_X\_8745589-8778998

###### Contingency table for cluster\_1:

| phenotype | wt | het | hom |
| --- | --- | --- | --- |
| alive | 11 | 22 | 12 |
| dead | 14 | 16 | 4 |

---

#### Baguida\_*gambiae*\_Delta

Baguida\_gambiae\_Delta\_2R:33967209-33981960  
Baguida\_gambiae\_Delta\_3L:3059554-3112922

##### Baguida\_gambiae\_Delta\_2R\_33967209-33981960

###### Contingency table for cluster\_1:

| phenotype | wt | het | hom |
| --- | --- | --- | --- |
| alive | 31 | 21 | 2 |
| dead | 31 | 3 | 0 |

##### Baguida\_gambiae\_Delta\_3L\_3059554-3112922

###### Contingency table for cluster\_1:

| phenotype | wt | het | hom |
| --- | --- | --- | --- |
| alive | 37 | 8 | 9 |
| dead | 11 | 21 | 2 |

###### Contingency table for cluster\_2:

| phenotype | wt | het | hom |
| --- | --- | --- | --- |
| alive | 39 | 14 | 1 |
| dead | 28 | 6 | 0 |

---

#### Korle-Bu\_*coluzzii*\_Delta

Korle-Bu\_coluzzii\_Delta\_2R:28145243-28168331  
Korle-Bu\_coluzzii\_Delta\_2R:39682306-39698081  
Korle-Bu\_coluzzii\_Delta\_3R:50971406-51021318  
Korle-Bu\_coluzzii\_Delta\_X:14653714-14668824  
Korle-Bu\_coluzzii\_Delta\_X:14668824-14684264  
Korle-Bu\_coluzzii\_Delta\_X:15670884-15702974  
Korle-Bu\_coluzzii\_Delta\_X:15702974-15756306

##### Korle-Bu\_coluzzii\_Delta\_2R\_28145243-28168331

###### Contingency table for cluster\_1:

| phenotype | wt | het | hom |
| --- | --- | --- | --- |
| alive | 52 | 7 | 0 |
| dead | 52 | 30 | 1 |

###### Contingency table for cluster\_2:

| phenotype | wt | het | hom |
| --- | --- | --- | --- |
| alive | 46 | 12 | 1 |
| dead | 69 | 13 | 1 |

###### Contingency table for cluster\_3:

| phenotype | wt | het | hom |
| --- | --- | --- | --- |
| alive | 50 | 9 | 0 |
| dead | 67 | 16 | 0 |

###### Contingency table for cluster\_4:

| phenotype | wt | het | hom |
| --- | --- | --- | --- |
| alive | 48 | 11 | 0 |
| dead | 72 | 11 | 0 |

##### Korle-Bu\_coluzzii\_Delta\_2R\_39682306-39698081

###### Contingency table for cluster\_1:

| phenotype | wt | het | hom |
| --- | --- | --- | --- |
| alive | 47 | 12 | 0 |
| dead | 65 | 16 | 2 |

###### Contingency table for cluster\_2:

| phenotype | wt | het | hom |
| --- | --- | --- | --- |
| alive | 42 | 16 | 1 |
| dead | 77 | 6 | 0 |

##### Korle-Bu\_coluzzii\_Delta\_3R\_50971406-51021318

###### Contingency table for cluster\_1:

| phenotype | wt | het | hom |
| --- | --- | --- | --- |
| alive | 44 | 13 | 2 |
| dead | 53 | 26 | 4 |

###### Contingency table for cluster\_2:

| phenotype | wt | het | hom |
| --- | --- | --- | --- |
| alive | 45 | 14 | 0 |
| dead | 75 | 7 | 1 |

###### Contingency table for cluster\_3:

| phenotype | wt | het | hom |
| --- | --- | --- | --- |
| alive | 49 | 10 | 0 |
| dead | 72 | 10 | 1 |

##### Korle-Bu\_coluzzii\_Delta\_X\_14653714-14668824

###### Contingency table for cluster\_1:

| phenotype | wt | het | hom |
| --- | --- | --- | --- |
| alive | 30 | 26 | 3 |
| dead | 58 | 24 | 1 |

###### Contingency table for cluster\_2:

| phenotype | wt | het | hom |
| --- | --- | --- | --- |
| alive | 49 | 9 | 1 |
| dead | 69 | 12 | 2 |

###### Contingency table for cluster\_3:

| phenotype | wt | het | hom |
| --- | --- | --- | --- |
| alive | 51 | 8 | 0 |
| dead | 70 | 13 | 0 |

##### Korle-Bu\_coluzzii\_Delta\_X\_14668824-14684264

###### Contingency table for cluster\_1:

| phenotype | wt | het | hom |
| --- | --- | --- | --- |
| alive | 31 | 25 | 3 |
| dead | 58 | 24 | 1 |

###### Contingency table for cluster\_2:

| phenotype | wt | het | hom |
| --- | --- | --- | --- |
| alive | 47 | 11 | 1 |
| dead | 69 | 12 | 2 |

###### Contingency table for cluster\_3:

| phenotype | wt | het | hom |
| --- | --- | --- | --- |
| alive | 51 | 8 | 0 |
| dead | 70 | 13 | 0 |

##### Korle-Bu\_coluzzii\_Delta\_X\_15670884-15702974

###### Contingency table for cluster\_1:

| phenotype | wt | het | hom |
| --- | --- | --- | --- |
| alive | 13 | 32 | 14 |
| dead | 37 | 35 | 11 |

###### Contingency table for cluster\_2:

| phenotype | wt | het | hom |
| --- | --- | --- | --- |
| alive | 52 | 7 | 0 |
| dead | 69 | 14 | 0 |

###### Contingency table for cluster\_3:

| phenotype | wt | het | hom |
| --- | --- | --- | --- |
| alive | 53 | 6 | 0 |
| dead | 68 | 15 | 0 |

##### Korle-Bu\_coluzzii\_Delta\_X\_15702974-15756306

###### Contingency table for cluster\_1:

| phenotype | wt | het | hom |
| --- | --- | --- | --- |
| alive | 12 | 38 | 9 |
| dead | 38 | 35 | 10 |

###### Contingency table for cluster\_2:

| phenotype | wt | het | hom |
| --- | --- | --- | --- |
| alive | 52 | 7 | 0 |
| dead | 68 | 14 | 1 |

###### Contingency table for cluster\_3:

| phenotype | wt | het | hom |
| --- | --- | --- | --- |
| alive | 52 | 7 | 0 |
| dead | 69 | 14 | 0 |

---

#### Madina\_*gambiae*\_Delta

Madina\_gambiae\_Delta\_2R:28401363-28416200  
Madina\_gambiae\_Delta\_2R:28454838-28474602

##### Madina\_gambiae\_Delta\_2R\_28401363-28416200

###### Contingency table for cluster\_1:

| phenotype | wt | het | hom |
| --- | --- | --- | --- |
| alive | 13 | 33 | 18 |
| dead | 14 | 21 | 2 |

##### Madina\_gambiae\_Delta\_2R\_28454838-28474602

###### Contingency table for cluster\_1:

| phenotype | wt | het | hom |
| --- | --- | --- | --- |
| alive | 8 | 33 | 23 |
| dead | 12 | 20 | 5 |

###### Contingency table for cluster\_2:

| phenotype | wt | het | hom |
| --- | --- | --- | --- |
| alive | 50 | 13 | 1 |
| dead | 26 | 10 | 1 |

---

#### Obuasi\_*gambiae*\_Delta

Obuasi\_gambiae\_Delta\_2R:28250033-28262588  
Obuasi\_gambiae\_Delta\_2R:28262588-28274893  
Obuasi\_gambiae\_Delta\_2R:28274893-28289873  
Obuasi\_gambiae\_Delta\_2R:28289873-28306606  
Obuasi\_gambiae\_Delta\_2R:28306605-28322681  
Obuasi\_gambiae\_Delta\_2R:28322681-28337951  
Obuasi\_gambiae\_Delta\_2R:28337951-28354693  
Obuasi\_gambiae\_Delta\_2R:28354692-28369784  
Obuasi\_gambiae\_Delta\_2R:28369784-28391434  
Obuasi\_gambiae\_Delta\_2R:28391433-28405240  
Obuasi\_gambiae\_Delta\_2R:28405239-28421116  
Obuasi\_gambiae\_Delta\_2R:28421116-28439220  
Obuasi\_gambiae\_Delta\_2R:28439219-28456407  
Obuasi\_gambiae\_Delta\_2R:28456407-28475225  
Obuasi\_gambiae\_Delta\_2R:28475225-28496115  
Obuasi\_gambiae\_Delta\_2R:28496114-28515482  
Obuasi\_gambiae\_Delta\_2R:28515481-28531112

##### Obuasi\_gambiae\_Delta\_2R\_28250033-28262588

###### Contingency table for cluster\_1:

| phenotype | wt | het | hom |
| --- | --- | --- | --- |
| alive | 11 | 16 | 0 |
| dead | 39 | 12 | 0 |

###### Contingency table for cluster\_2:

| phenotype | wt | het | hom |
| --- | --- | --- | --- |
| alive | 23 | 4 | 0 |
| dead | 33 | 16 | 2 |

##### Obuasi\_gambiae\_Delta\_2R\_28262588-28274893

###### Contingency table for cluster\_1:

| phenotype | wt | het | hom |
| --- | --- | --- | --- |
| alive | 10 | 15 | 2 |
| dead | 39 | 12 | 0 |

###### Contingency table for cluster\_2:

| phenotype | wt | het | hom |
| --- | --- | --- | --- |
| alive | 23 | 4 | 0 |
| dead | 33 | 16 | 2 |

##### Obuasi\_gambiae\_Delta\_2R\_28274893-28289873

###### Contingency table for cluster\_1:

| phenotype | wt | het | hom |
| --- | --- | --- | --- |
| alive | 10 | 14 | 3 |
| dead | 39 | 12 | 0 |

###### Contingency table for cluster\_2:

| phenotype | wt | het | hom |
| --- | --- | --- | --- |
| alive | 23 | 4 | 0 |
| dead | 33 | 16 | 2 |

##### Obuasi\_gambiae\_Delta\_2R\_28289873-28306606

###### Contingency table for cluster\_1:

| phenotype | wt | het | hom |
| --- | --- | --- | --- |
| alive | 9 | 14 | 4 |
| dead | 39 | 12 | 0 |

###### Contingency table for cluster\_2:

| phenotype | wt | het | hom |
| --- | --- | --- | --- |
| alive | 23 | 4 | 0 |
| dead | 33 | 17 | 1 |

##### Obuasi\_gambiae\_Delta\_2R\_28306605-28322681

###### Contingency table for cluster\_1:

| phenotype | wt | het | hom |
| --- | --- | --- | --- |
| alive | 10 | 14 | 3 |
| dead | 38 | 13 | 0 |

###### Contingency table for cluster\_2:

| phenotype | wt | het | hom |
| --- | --- | --- | --- |
| alive | 23 | 4 | 0 |
| dead | 33 | 17 | 1 |

##### Obuasi\_gambiae\_Delta\_2R\_28322681-28337951

###### Contingency table for cluster\_1:

| phenotype | wt | het | hom |
| --- | --- | --- | --- |
| alive | 9 | 14 | 4 |
| dead | 37 | 14 | 0 |

###### Contingency table for cluster\_2:

| phenotype | wt | het | hom |
| --- | --- | --- | --- |
| alive | 23 | 4 | 0 |
| dead | 34 | 16 | 1 |

###### Contingency table for cluster\_3:

| phenotype | wt | het | hom |
| --- | --- | --- | --- |
| alive | 24 | 3 | 0 |
| dead | 34 | 16 | 1 |

##### Obuasi\_gambiae\_Delta\_2R\_28337951-28354693

###### Contingency table for cluster\_1:

| phenotype | wt | het | hom |
| --- | --- | --- | --- |
| alive | 9 | 14 | 4 |
| dead | 36 | 15 | 0 |

###### Contingency table for cluster\_2:

| phenotype | wt | het | hom |
| --- | --- | --- | --- |
| alive | 24 | 3 | 0 |
| dead | 33 | 17 | 1 |

###### Contingency table for cluster\_3:

| phenotype | wt | het | hom |
| --- | --- | --- | --- |
| alive | 22 | 5 | 0 |
| dead | 36 | 14 | 1 |

##### Obuasi\_gambiae\_Delta\_2R\_28354692-28369784

###### Contingency table for cluster\_1:

| phenotype | wt | het | hom |
| --- | --- | --- | --- |
| alive | 8 | 15 | 4 |
| dead | 35 | 16 | 0 |

###### Contingency table for cluster\_2:

| phenotype | wt | het | hom |
| --- | --- | --- | --- |
| alive | 24 | 3 | 0 |
| dead | 32 | 17 | 2 |

###### Contingency table for cluster\_3:

| phenotype | wt | het | hom |
| --- | --- | --- | --- |
| alive | 21 | 6 | 0 |
| dead | 36 | 14 | 1 |

##### Obuasi\_gambiae\_Delta\_2R\_28369784-28391434

###### Contingency table for cluster\_1:

| phenotype | wt | het | hom |
| --- | --- | --- | --- |
| alive | 8 | 15 | 4 |
| dead | 35 | 15 | 1 |

###### Contingency table for cluster\_2:

| phenotype | wt | het | hom |
| --- | --- | --- | --- |
| alive | 24 | 3 | 0 |
| dead | 32 | 17 | 2 |

###### Contingency table for cluster\_3:

| phenotype | wt | het | hom |
| --- | --- | --- | --- |
| alive | 21 | 6 | 0 |
| dead | 35 | 15 | 1 |

##### Obuasi\_gambiae\_Delta\_2R\_28391433-28405240

###### Contingency table for cluster\_1:

| phenotype | wt | het | hom |
| --- | --- | --- | --- |
| alive | 7 | 16 | 4 |
| dead | 34 | 15 | 2 |

###### Contingency table for cluster\_2:

| phenotype | wt | het | hom |
| --- | --- | --- | --- |
| alive | 24 | 3 | 0 |
| dead | 32 | 17 | 2 |

###### Contingency table for cluster\_3:

| phenotype | wt | het | hom |
| --- | --- | --- | --- |
| alive | 21 | 6 | 0 |
| dead | 35 | 15 | 1 |

##### Obuasi\_gambiae\_Delta\_2R\_28405239-28421116

###### Contingency table for cluster\_1:

| phenotype | wt | het | hom |
| --- | --- | --- | --- |
| alive | 9 | 11 | 7 |
| dead | 32 | 17 | 2 |

###### Contingency table for cluster\_2:

| phenotype | wt | het | hom |
| --- | --- | --- | --- |
| alive | 21 | 6 | 0 |
| dead | 35 | 15 | 1 |

###### Contingency table for cluster\_3:

| phenotype | wt | het | hom |
| --- | --- | --- | --- |
| alive | 24 | 3 | 0 |
| dead | 33 | 16 | 2 |

##### Obuasi\_gambiae\_Delta\_2R\_28421116-28439220

###### Contingency table for cluster\_1:

| phenotype | wt | het | hom |
| --- | --- | --- | --- |
| alive | 7 | 13 | 7 |
| dead | 29 | 18 | 4 |

###### Contingency table for cluster\_2:

| phenotype | wt | het | hom |
| --- | --- | --- | --- |
| alive | 24 | 3 | 0 |
| dead | 33 | 16 | 2 |

###### Contingency table for cluster\_3:

| phenotype | wt | het | hom |
| --- | --- | --- | --- |
| alive | 21 | 6 | 0 |
| dead | 36 | 14 | 1 |

##### Obuasi\_gambiae\_Delta\_2R\_28439219-28456407

###### Contingency table for cluster\_1:

| phenotype | wt | het | hom |
| --- | --- | --- | --- |
| alive | 4 | 14 | 9 |
| dead | 27 | 17 | 7 |

###### Contingency table for cluster\_2:

| phenotype | wt | het | hom |
| --- | --- | --- | --- |
| alive | 23 | 4 | 0 |
| dead | 32 | 17 | 2 |

###### Contingency table for cluster\_3:

| phenotype | wt | het | hom |
| --- | --- | --- | --- |
| alive | 20 | 7 | 0 |
| dead | 36 | 14 | 1 |

##### Obuasi\_gambiae\_Delta\_2R\_28456407-28475225

###### Contingency table for cluster\_1:

| phenotype | wt | het | hom |
| --- | --- | --- | --- |
| alive | 3 | 15 | 9 |
| dead | 25 | 19 | 7 |

###### Contingency table for cluster\_2:

| phenotype | wt | het | hom |
| --- | --- | --- | --- |
| alive | 23 | 4 | 0 |
| dead | 32 | 17 | 2 |

###### Contingency table for cluster\_3:

| phenotype | wt | het | hom |
| --- | --- | --- | --- |
| alive | 20 | 7 | 0 |
| dead | 38 | 12 | 1 |

##### Obuasi\_gambiae\_Delta\_2R\_28475225-28496115

###### Contingency table for cluster\_1:

| phenotype | wt | het | hom |
| --- | --- | --- | --- |
| alive | 7 | 12 | 8 |
| dead | 28 | 18 | 5 |

###### Contingency table for cluster\_2:

| phenotype | wt | het | hom |
| --- | --- | --- | --- |
| alive | 19 | 8 | 0 |
| dead | 39 | 11 | 1 |

##### Obuasi\_gambiae\_Delta\_2R\_28496114-28515482

###### Contingency table for cluster\_1:

| phenotype | wt | het | hom |
| --- | --- | --- | --- |
| alive | 11 | 9 | 7 |
| dead | 30 | 16 | 5 |

###### Contingency table for cluster\_2:

| phenotype | wt | het | hom |
| --- | --- | --- | --- |
| alive | 17 | 10 | 0 |
| dead | 38 | 12 | 1 |

##### Obuasi\_gambiae\_Delta\_2R\_28515481-28531112

###### Contingency table for cluster\_1:

| phenotype | wt | het | hom |
| --- | --- | --- | --- |
| alive | 13 | 11 | 3 |
| dead | 35 | 12 | 4 |

###### Contingency table for cluster\_2:

| phenotype | wt | het | hom |
| --- | --- | --- | --- |
| alive | 14 | 12 | 1 |
| dead | 38 | 12 | 1 |

---

#### Baguida\_*gambiae*\_PM

Baguida\_gambiae\_PM\_X:15411876-15495650  
Baguida\_gambiae\_PM\_X:9165646-9222299  
Baguida\_gambiae\_PM\_X:9222298-9273975  
Baguida\_gambiae\_PM\_X:9273975-9317640  
Baguida\_gambiae\_PM\_X:9317639-9346605

##### Baguida\_gambiae\_PM\_X\_15411876-15495650

###### Contingency table for cluster\_1:

| phenotype | wt | het | hom |
| --- | --- | --- | --- |
| alive | 17 | 14 | 4 |
| dead | 13 | 12 | 5 |

###### Contingency table for cluster\_2:

| phenotype | wt | het | hom |
| --- | --- | --- | --- |
| alive | 24 | 11 | 0 |
| dead | 12 | 16 | 2 |

###### Contingency table for cluster\_3:

| phenotype | wt | het | hom |
| --- | --- | --- | --- |
| alive | 20 | 11 | 4 |
| dead | 24 | 5 | 1 |

##### Baguida\_gambiae\_PM\_X\_9165646-9222299

###### Contingency table for cluster\_1:

| phenotype | wt | het | hom |
| --- | --- | --- | --- |
| alive | 6 | 13 | 16 |
| dead | 9 | 13 | 8 |

###### Contingency table for cluster\_2:

| phenotype | wt | het | hom |
| --- | --- | --- | --- |
| alive | 24 | 10 | 1 |
| dead | 10 | 14 | 6 |

##### Baguida\_gambiae\_PM\_X\_9222298-9273975

###### Contingency table for cluster\_1:

| phenotype | wt | het | hom |
| --- | --- | --- | --- |
| alive | 16 | 13 | 6 |
| dead | 13 | 15 | 2 |

###### Contingency table for cluster\_2:

| phenotype | wt | het | hom |
| --- | --- | --- | --- |
| alive | 24 | 10 | 1 |
| dead | 10 | 14 | 6 |

##### Baguida\_gambiae\_PM\_X\_9273975-9317640

###### Contingency table for cluster\_1:

| phenotype | wt | het | hom |
| --- | --- | --- | --- |
| alive | 24 | 10 | 1 |
| dead | 10 | 14 | 6 |

###### Contingency table for cluster\_2:

| phenotype | wt | het | hom |
| --- | --- | --- | --- |
| alive | 20 | 11 | 4 |
| dead | 19 | 11 | 0 |

##### Baguida\_gambiae\_PM\_X\_9317639-9346605

###### Contingency table for cluster\_1:

| phenotype | wt | het | hom |
| --- | --- | --- | --- |
| alive | 24 | 10 | 1 |
| dead | 11 | 14 | 5 |

###### Contingency table for cluster\_2:

| phenotype | wt | het | hom |
| --- | --- | --- | --- |
| alive | 19 | 14 | 2 |
| dead | 22 | 8 | 0 |

---

#### Korle-Bu\_*coluzzii*\_PM

Korle-Bu\_coluzzii\_PM\_2R:38097626-38111826  
Korle-Bu\_coluzzii\_PM\_2R:54371673-54397846  
Korle-Bu\_coluzzii\_PM\_3R:28482794-28505651  
Korle-Bu\_coluzzii\_PM\_3R:28505651-28533744  
Korle-Bu\_coluzzii\_PM\_3R:28563392-28595112  
Korle-Bu\_coluzzii\_PM\_3R:2929236-2946500

##### Korle-Bu\_coluzzii\_PM\_2R\_38097626-38111826

##### Korle-Bu\_coluzzii\_PM\_2R\_54371673-54397846

###### Contingency table for cluster\_1:

| phenotype | wt | het | hom |
| --- | --- | --- | --- |
| alive | 30 | 22 | 5 |
| dead | 34 | 9 | 5 |

##### Korle-Bu\_coluzzii\_PM\_3R\_28482794-28505651

###### Contingency table for cluster\_1:

| phenotype | wt | het | hom |
| --- | --- | --- | --- |
| alive | 34 | 20 | 3 |
| dead | 14 | 28 | 6 |

###### Contingency table for cluster\_2:

| phenotype | wt | het | hom |
| --- | --- | --- | --- |
| alive | 34 | 19 | 4 |
| dead | 40 | 8 | 0 |

###### Contingency table for cluster\_3:

| phenotype | wt | het | hom |
| --- | --- | --- | --- |
| alive | 39 | 17 | 1 |
| dead | 41 | 6 | 1 |

##### Korle-Bu\_coluzzii\_PM\_3R\_28505651-28533744

###### Contingency table for cluster\_1:

| phenotype | wt | het | hom |
| --- | --- | --- | --- |
| alive | 37 | 17 | 3 |
| dead | 15 | 29 | 4 |

###### Contingency table for cluster\_2:

| phenotype | wt | het | hom |
| --- | --- | --- | --- |
| alive | 26 | 24 | 7 |
| dead | 31 | 16 | 1 |

###### Contingency table for cluster\_3:

| phenotype | wt | het | hom |
| --- | --- | --- | --- |
| alive | 39 | 15 | 3 |
| dead | 40 | 7 | 1 |

##### Korle-Bu\_coluzzii\_PM\_3R\_28563392-28595112

###### Contingency table for cluster\_1:

| phenotype | wt | het | hom |
| --- | --- | --- | --- |
| alive | 20 | 27 | 10 |
| dead | 20 | 25 | 3 |

###### Contingency table for cluster\_2:

| phenotype | wt | het | hom |
| --- | --- | --- | --- |
| alive | 38 | 17 | 2 |
| dead | 18 | 25 | 5 |

###### Contingency table for cluster\_3:

| phenotype | wt | het | hom |
| --- | --- | --- | --- |
| alive | 38 | 16 | 3 |
| dead | 41 | 6 | 1 |

##### Korle-Bu\_coluzzii\_PM\_3R\_2929236-2946500

###### Contingency table for cluster\_1:

| phenotype | wt | het | hom |
| --- | --- | --- | --- |
| alive | 24 | 26 | 7 |
| dead | 7 | 27 | 14 |

---

#### Madina\_*gambiae*\_PM

Madina\_gambiae\_PM\_2R:54255612-54278008

##### Madina\_gambiae\_PM\_2R\_54255612-54278008

###### Contingency table for cluster\_1:

| phenotype | wt | het | hom |
| --- | --- | --- | --- |
| alive | 31 | 7 | 0 |
| dead | 14 | 12 | 1 |

---

#### Obuasi\_*gambiae*\_PM

Obuasi\_gambiae\_PM\_2R:3474360-3509400  
Obuasi\_gambiae\_PM\_3L:30477135-30487616  
Obuasi\_gambiae\_PM\_3L:30487615-30497759  
Obuasi\_gambiae\_PM\_3L:30497758-30506894  
Obuasi\_gambiae\_PM\_3L:30506893-30516797  
Obuasi\_gambiae\_PM\_3L:30534828-30545854  
Obuasi\_gambiae\_PM\_3L:30554459-30562573  
Obuasi\_gambiae\_PM\_3L:30562572-30578511  
Obuasi\_gambiae\_PM\_3L:30578511-30587941  
Obuasi\_gambiae\_PM\_3L:30587941-30597762  
Obuasi\_gambiae\_PM\_3L:30597761-30607138  
Obuasi\_gambiae\_PM\_3L:30607137-30617085  
Obuasi\_gambiae\_PM\_3L:30617085-30624950  
Obuasi\_gambiae\_PM\_3L:30624949-30635698  
Obuasi\_gambiae\_PM\_3L:30635698-30646602  
Obuasi\_gambiae\_PM\_3L:30646601-30657763  
Obuasi\_gambiae\_PM\_3L:30657762-30667355  
Obuasi\_gambiae\_PM\_3L:30667355-30678119  
Obuasi\_gambiae\_PM\_3L:30678119-30687769  
Obuasi\_gambiae\_PM\_3L:30687769-30697180  
Obuasi\_gambiae\_PM\_3R:48454456-48494640

##### Obuasi\_gambiae\_PM\_2R\_3474360-3509400

###### Contingency table for cluster\_1:

| phenotype | wt | het | hom |
| --- | --- | --- | --- |
| alive | 1 | 2 | 51 |
| dead | 1 | 16 | 33 |

##### Obuasi\_gambiae\_PM\_3L\_30477135-30487616

###### Contingency table for cluster\_1:

| phenotype | wt | het | hom |
| --- | --- | --- | --- |
| alive | 37 | 16 | 1 |
| dead | 17 | 28 | 5 |

##### Obuasi\_gambiae\_PM\_3L\_30487615-30497759

###### Contingency table for cluster\_1:

| phenotype | wt | het | hom |
| --- | --- | --- | --- |
| alive | 33 | 20 | 1 |
| dead | 14 | 29 | 7 |

##### Obuasi\_gambiae\_PM\_3L\_30497758-30506894

###### Contingency table for cluster\_1:

| phenotype | wt | het | hom |
| --- | --- | --- | --- |
| alive | 36 | 17 | 1 |
| dead | 13 | 31 | 6 |

##### Obuasi\_gambiae\_PM\_3L\_30506893-30516797

###### Contingency table for cluster\_1:

| phenotype | wt | het | hom |
| --- | --- | --- | --- |
| alive | 35 | 19 | 0 |
| dead | 15 | 29 | 6 |

##### Obuasi\_gambiae\_PM\_3L\_30534828-30545854

###### Contingency table for cluster\_1:

| phenotype | wt | het | hom |
| --- | --- | --- | --- |
| alive | 35 | 18 | 1 |
| dead | 15 | 31 | 4 |

##### Obuasi\_gambiae\_PM\_3L\_30554459-30562573

###### Contingency table for cluster\_1:

| phenotype | wt | het | hom |
| --- | --- | --- | --- |
| alive | 36 | 17 | 1 |
| dead | 16 | 29 | 5 |

##### Obuasi\_gambiae\_PM\_3L\_30562572-30578511

###### Contingency table for cluster\_1:

| phenotype | wt | het | hom |
| --- | --- | --- | --- |
| alive | 36 | 16 | 2 |
| dead | 16 | 29 | 5 |

##### Obuasi\_gambiae\_PM\_3L\_30578511-30587941

###### Contingency table for cluster\_1:

| phenotype | wt | het | hom |
| --- | --- | --- | --- |
| alive | 35 | 17 | 2 |
| dead | 15 | 28 | 7 |

##### Obuasi\_gambiae\_PM\_3L\_30587941-30597762

###### Contingency table for cluster\_1:

| phenotype | wt | het | hom |
| --- | --- | --- | --- |
| alive | 35 | 17 | 2 |
| dead | 15 | 28 | 7 |

##### Obuasi\_gambiae\_PM\_3L\_30597761-30607138

###### Contingency table for cluster\_1:

| phenotype | wt | het | hom |
| --- | --- | --- | --- |
| alive | 35 | 17 | 2 |
| dead | 15 | 28 | 7 |

##### Obuasi\_gambiae\_PM\_3L\_30607137-30617085

###### Contingency table for cluster\_1:

| phenotype | wt | het | hom |
| --- | --- | --- | --- |
| alive | 34 | 18 | 2 |
| dead | 15 | 27 | 8 |

##### Obuasi\_gambiae\_PM\_3L\_30617085-30624950

###### Contingency table for cluster\_1:

| phenotype | wt | het | hom |
| --- | --- | --- | --- |
| alive | 33 | 18 | 3 |
| dead | 15 | 25 | 10 |

##### Obuasi\_gambiae\_PM\_3L\_30624949-30635698

###### Contingency table for cluster\_1:

| phenotype | wt | het | hom |
| --- | --- | --- | --- |
| alive | 30 | 21 | 3 |
| dead | 10 | 28 | 12 |

##### Obuasi\_gambiae\_PM\_3L\_30635698-30646602

###### Contingency table for cluster\_1:

| phenotype | wt | het | hom |
| --- | --- | --- | --- |
| alive | 30 | 21 | 3 |
| dead | 10 | 28 | 12 |

##### Obuasi\_gambiae\_PM\_3L\_30646601-30657763

###### Contingency table for cluster\_1:

| phenotype | wt | het | hom |
| --- | --- | --- | --- |
| alive | 28 | 24 | 2 |
| dead | 11 | 29 | 10 |

##### Obuasi\_gambiae\_PM\_3L\_30657762-30667355

###### Contingency table for cluster\_1:

| phenotype | wt | het | hom |
| --- | --- | --- | --- |
| alive | 28 | 24 | 2 |
| dead | 11 | 30 | 9 |

##### Obuasi\_gambiae\_PM\_3L\_30667355-30678119

###### Contingency table for cluster\_1:

| phenotype | wt | het | hom |
| --- | --- | --- | --- |
| alive | 27 | 25 | 2 |
| dead | 10 | 31 | 9 |

##### Obuasi\_gambiae\_PM\_3L\_30678119-30687769

###### Contingency table for cluster\_1:

| phenotype | wt | het | hom |
| --- | --- | --- | --- |
| alive | 26 | 26 | 2 |
| dead | 11 | 29 | 10 |

##### Obuasi\_gambiae\_PM\_3L\_30687769-30697180

###### Contingency table for cluster\_1:

| phenotype | wt | het | hom |
| --- | --- | --- | --- |
| alive | 26 | 26 | 2 |
| dead | 9 | 30 | 11 |

##### Obuasi\_gambiae\_PM\_3R\_48454456-48494640

###### Contingency table for cluster\_1:

| phenotype | wt | het | hom |
| --- | --- | --- | --- |
| alive | 24 | 24 | 6 |
| dead | 39 | 9 | 2 |

###### Contingency table for cluster\_2:

| phenotype | wt | het | hom |
| --- | --- | --- | --- |
| alive | 43 | 10 | 1 |
| dead | 38 | 9 | 3 |

###### Contingency table for cluster\_3:

| phenotype | wt | het | hom |
| --- | --- | --- | --- |
| alive | 44 | 10 | 0 |
| dead | 39 | 10 | 1 |

---
