## Supplementary Data S4 for "Genome-wide association studies reveal novel loci associated with pyrethroid and organophosphate resistance in *Anopheles gambiae* s.l.": Supplementary_Data_S4.html

window\_H12\_summary


### H12 windows of interest

For each sample set, we first provide a summary plot of *Δ*H_12_ (H_12_ in
susceptible samples subtracted from that in resistant samples) across
the genome, with *Δ*H_12_ shown in green and
the results of the 200 randomisations shown behind in grey. Windows
identified as peaks are highlighted by points, colour-coded by whether
they are significantly higher than expected based on the simulations
(green) or not (purple). For each significant window (green points), we
then provide its own plot showing the significant (*P* < 0.01)
SNPs found in the region of that window, and their -log10(Pvalue) of
association with phenotype. Red points indicate non-synonymous SNPs,
blue points indicate all other SNPs. Point shape indicates whether the
mutant allele at that SNP is associated with increased (circle) or
decreased (triangle) resistance. Dark points in the centre of the plot
show SNPs within the significant window, light points on the sides show
SNPs in the region 10,000 bp either side of the window.

Legend  
Avrankou\_*coluzzii*\_Delta  
Baguida\_*gambiae*\_Delta  
Korle-Bu\_*coluzzii*\_Delta  
Madina\_*gambiae*\_Delta  
Obuasi\_*gambiae*\_Delta  
Baguida\_*gambiae*\_PM  
Korle-Bu\_*coluzzii*\_PM  
Madina\_*gambiae*\_PM  
Obuasi\_*gambiae*\_PM

---

#### Plot legend

---

#### Avrankou\_*coluzzii*\_Delta

Avrankou\_coluzzii\_Delta\_2L:3627850  
Avrankou\_coluzzii\_Delta\_2L:39336219  
Avrankou\_coluzzii\_Delta\_2L:40020139  
Avrankou\_coluzzii\_Delta\_2R:55812794  
Avrankou\_coluzzii\_Delta\_3R:46530542  
Avrankou\_coluzzii\_Delta\_3R:46606399  
Avrankou\_coluzzii\_Delta\_3R:46697535  
Avrankou\_coluzzii\_Delta\_3R:46784988  
Avrankou\_coluzzii\_Delta\_3R:46831369  
Avrankou\_coluzzii\_Delta\_3R:46856221  
Avrankou\_coluzzii\_Delta\_3R:48501462

##### Avrankou\_coluzzii\_Delta\_2L\_3627850

##### Avrankou\_coluzzii\_Delta\_2L\_39336219

##### Avrankou\_coluzzii\_Delta\_2L\_40020139

##### Avrankou\_coluzzii\_Delta\_2R\_55812794

##### Avrankou\_coluzzii\_Delta\_3R\_46530542

##### Avrankou\_coluzzii\_Delta\_3R\_46606399

##### Avrankou\_coluzzii\_Delta\_3R\_46697535

##### Avrankou\_coluzzii\_Delta\_3R\_46784988

##### Avrankou\_coluzzii\_Delta\_3R\_46831369

##### Avrankou\_coluzzii\_Delta\_3R\_46856221

##### Avrankou\_coluzzii\_Delta\_3R\_48501462

---

#### Baguida\_*gambiae*\_Delta

Baguida\_gambiae\_Delta\_2L:20037401  
Baguida\_gambiae\_Delta\_2R:3042129  
Baguida\_gambiae\_Delta\_2R:3519707

##### Baguida\_gambiae\_Delta\_2L\_20037401

##### Baguida\_gambiae\_Delta\_2R\_3042129

##### Baguida\_gambiae\_Delta\_2R\_3519707

---

#### Korle-Bu\_*coluzzii*\_Delta

Korle-Bu\_coluzzii\_Delta\_2R:28546290  
Korle-Bu\_coluzzii\_Delta\_2R:54064272  
Korle-Bu\_coluzzii\_Delta\_3R:4030785  
Korle-Bu\_coluzzii\_Delta\_X:15661082

##### Korle-Bu\_coluzzii\_Delta\_2R\_28546290

##### Korle-Bu\_coluzzii\_Delta\_2R\_54064272

##### Korle-Bu\_coluzzii\_Delta\_3R\_4030785

##### Korle-Bu\_coluzzii\_Delta\_X\_15661082

---

#### Madina\_*gambiae*\_Delta

Madina\_gambiae\_Delta\_2L:14217696  
Madina\_gambiae\_Delta\_2L:14284997  
Madina\_gambiae\_Delta\_2L:27986348  
Madina\_gambiae\_Delta\_2L:28023911  
Madina\_gambiae\_Delta\_2L:28093738  
Madina\_gambiae\_Delta\_2L:28110739  
Madina\_gambiae\_Delta\_2L:28130562  
Madina\_gambiae\_Delta\_2L:28174232  
Madina\_gambiae\_Delta\_2L:31419497  
Madina\_gambiae\_Delta\_2L:3246748  
Madina\_gambiae\_Delta\_2L:3287434  
Madina\_gambiae\_Delta\_2L:3626021  
Madina\_gambiae\_Delta\_X:14215545

##### Madina\_gambiae\_Delta\_2L\_14217696

##### Madina\_gambiae\_Delta\_2L\_14284997

##### Madina\_gambiae\_Delta\_2L\_27986348

##### Madina\_gambiae\_Delta\_2L\_28023911

##### Madina\_gambiae\_Delta\_2L\_28093738

##### Madina\_gambiae\_Delta\_2L\_28110739

##### Madina\_gambiae\_Delta\_2L\_28130562

##### Madina\_gambiae\_Delta\_2L\_28174232

##### Madina\_gambiae\_Delta\_2L\_31419497

##### Madina\_gambiae\_Delta\_2L\_3246748

##### Madina\_gambiae\_Delta\_2L\_3287434

##### Madina\_gambiae\_Delta\_2L\_3626021

##### Madina\_gambiae\_Delta\_X\_14215545

---

#### Obuasi\_*gambiae*\_Delta

Obuasi\_gambiae\_Delta\_2L:25666039  
Obuasi\_gambiae\_Delta\_2L:47246666  
Obuasi\_gambiae\_Delta\_2R:3175305  
Obuasi\_gambiae\_Delta\_2R:40905141  
Obuasi\_gambiae\_Delta\_3R:10633692  
Obuasi\_gambiae\_Delta\_X:15939854

##### Obuasi\_gambiae\_Delta\_2L\_25666039

##### Obuasi\_gambiae\_Delta\_2L\_47246666

##### Obuasi\_gambiae\_Delta\_2R\_3175305

##### Obuasi\_gambiae\_Delta\_2R\_40905141

##### Obuasi\_gambiae\_Delta\_3R\_10633692

##### Obuasi\_gambiae\_Delta\_X\_15939854

---

#### Baguida\_*gambiae*\_PM

Baguida\_gambiae\_PM\_2L:19523767  
Baguida\_gambiae\_PM\_2L:3266679  
Baguida\_gambiae\_PM\_2L:3471571  
Baguida\_gambiae\_PM\_2L:3517468  
Baguida\_gambiae\_PM\_2L:3578649  
Baguida\_gambiae\_PM\_2R:28578188  
Baguida\_gambiae\_PM\_2R:28622655

##### Baguida\_gambiae\_PM\_2L\_19523767

##### Baguida\_gambiae\_PM\_2L\_3266679

##### Baguida\_gambiae\_PM\_2L\_3471571

##### Baguida\_gambiae\_PM\_2L\_3517468

##### Baguida\_gambiae\_PM\_2L\_3578649

##### Baguida\_gambiae\_PM\_2R\_28578188

##### Baguida\_gambiae\_PM\_2R\_28622655

---

#### Korle-Bu\_*coluzzii*\_PM

Korle-Bu\_coluzzii\_PM\_2L:22237788  
Korle-Bu\_coluzzii\_PM\_2L:23989307  
Korle-Bu\_coluzzii\_PM\_2L:28414771  
Korle-Bu\_coluzzii\_PM\_2L:28438932  
Korle-Bu\_coluzzii\_PM\_2L:28561906  
Korle-Bu\_coluzzii\_PM\_2L:28615047  
Korle-Bu\_coluzzii\_PM\_2L:28641468  
Korle-Bu\_coluzzii\_PM\_2L:28664089  
Korle-Bu\_coluzzii\_PM\_2L:28689296  
Korle-Bu\_coluzzii\_PM\_2L:36913180  
Korle-Bu\_coluzzii\_PM\_2L:36939137  
Korle-Bu\_coluzzii\_PM\_2L:36963255  
Korle-Bu\_coluzzii\_PM\_2L:37011181  
Korle-Bu\_coluzzii\_PM\_2L:37103650  
Korle-Bu\_coluzzii\_PM\_2L:37142451  
Korle-Bu\_coluzzii\_PM\_2L:37176411  
Korle-Bu\_coluzzii\_PM\_2L:37570900  
Korle-Bu\_coluzzii\_PM\_2L:3821991  
Korle-Bu\_coluzzii\_PM\_3R:4253878  
Korle-Bu\_coluzzii\_PM\_X:15614925

##### Korle-Bu\_coluzzii\_PM\_2L\_22237788

##### Korle-Bu\_coluzzii\_PM\_2L\_23989307

##### Korle-Bu\_coluzzii\_PM\_2L\_28414771

##### Korle-Bu\_coluzzii\_PM\_2L\_28438932

##### Korle-Bu\_coluzzii\_PM\_2L\_28561906

##### Korle-Bu\_coluzzii\_PM\_2L\_28615047

##### Korle-Bu\_coluzzii\_PM\_2L\_28641468

##### Korle-Bu\_coluzzii\_PM\_2L\_28664089

##### Korle-Bu\_coluzzii\_PM\_2L\_28689296

##### Korle-Bu\_coluzzii\_PM\_2L\_36913180

##### Korle-Bu\_coluzzii\_PM\_2L\_36939137

##### Korle-Bu\_coluzzii\_PM\_2L\_36963255

##### Korle-Bu\_coluzzii\_PM\_2L\_37011181

##### Korle-Bu\_coluzzii\_PM\_2L\_37103650

##### Korle-Bu\_coluzzii\_PM\_2L\_37142451

##### Korle-Bu\_coluzzii\_PM\_2L\_37176411

##### Korle-Bu\_coluzzii\_PM\_2L\_37570900

##### Korle-Bu\_coluzzii\_PM\_2L\_3821991

##### Korle-Bu\_coluzzii\_PM\_3R\_4253878

##### Korle-Bu\_coluzzii\_PM\_X\_15614925

---

#### Madina\_*gambiae*\_PM

Madina\_gambiae\_PM\_2L:31835378  
Madina\_gambiae\_PM\_2L:32812183  
Madina\_gambiae\_PM\_2L:39511138  
Madina\_gambiae\_PM\_2L:41724046  
Madina\_gambiae\_PM\_2R:3004025  
Madina\_gambiae\_PM\_2R:3037081  
Madina\_gambiae\_PM\_3R:28433993  
Madina\_gambiae\_PM\_3R:28591533

##### Madina\_gambiae\_PM\_2L\_31835378

##### Madina\_gambiae\_PM\_2L\_32812183

##### Madina\_gambiae\_PM\_2L\_39511138

##### Madina\_gambiae\_PM\_2L\_41724046

##### Madina\_gambiae\_PM\_2R\_3004025

##### Madina\_gambiae\_PM\_2R\_3037081

##### Madina\_gambiae\_PM\_3R\_28433993

##### Madina\_gambiae\_PM\_3R\_28591533

---

#### Obuasi\_*gambiae*\_PM

Obuasi\_gambiae\_PM\_2R:3979039  
Obuasi\_gambiae\_PM\_3L:11361643  
Obuasi\_gambiae\_PM\_3L:11384773  
Obuasi\_gambiae\_PM\_X:13850824  
Obuasi\_gambiae\_PM\_X:14004692

##### Obuasi\_gambiae\_PM\_2R\_3979039

##### Obuasi\_gambiae\_PM\_3L\_11361643

##### Obuasi\_gambiae\_PM\_3L\_11384773

##### Obuasi\_gambiae\_PM\_X\_13850824

##### Obuasi\_gambiae\_PM\_X\_14004692

---
