## Supplementary Data S5 for "Genome-wide association studies reveal novel loci associated with pyrethroid and organophosphate resistance in *Anopheles gambiae* s.l.": Supplementary_Data_S5.html

window\_PBS\_summary


### PBS windows of interest

For each sample set, we first provide a summary plot of PBS across
the genome, with PBS shown in blue and the results of the 200
randomisations shown behind in grey. Windows identified as peaks are
highlighted by points, colour-coded by whether they are significantly
higher than expected based on the simulations (green) or not (purple).
For each significant window (green points), we then provide its own plot
showing the significant (*P* < 0.01) SNPs found in the region
of that window, and their -log10(Pvalue) of association with phenotype.
Red points indicate non-synonymous SNPs, blue points indicate all other
SNPs. Point shape indicates whether the mutant allele at that SNP is
associated with increased (circle) or decreased (triangle) resistance.
Dark points in the centre of the plot show SNPs within the significant
window, light points on the sides show SNPs in the region 10,000 bp
either side of the window.

Legend  
Avrankou\_*coluzzii*\_Delta  
Baguida\_*gambiae*\_Delta  
Korle-Bu\_*coluzzii*\_Delta  
Madina\_*gambiae*\_Delta  
Obuasi\_*gambiae*\_Delta  
Baguida\_*gambiae*\_PM  
Korle-Bu\_*coluzzii*\_PM  
Madina\_*gambiae*\_PM  
Obuasi\_*gambiae*\_PM

---

#### Plot legend

---

#### Avrankou\_*coluzzii*\_Delta

Avrankou\_coluzzii\_Delta\_2L:10075762  
Avrankou\_coluzzii\_Delta\_2R:36536888  
Avrankou\_coluzzii\_Delta\_2R:40059143  
Avrankou\_coluzzii\_Delta\_2R:50397087  
Avrankou\_coluzzii\_Delta\_2R:57936779  
Avrankou\_coluzzii\_Delta\_2R:58017231  
Avrankou\_coluzzii\_Delta\_3R:40583507  
Avrankou\_coluzzii\_Delta\_3R:42095383  
Avrankou\_coluzzii\_Delta\_3R:43486612  
Avrankou\_coluzzii\_Delta\_3R:43532647  
Avrankou\_coluzzii\_Delta\_X:16349957

##### Avrankou\_coluzzii\_Delta\_2L\_10075762

##### Avrankou\_coluzzii\_Delta\_2R\_36536888

##### Avrankou\_coluzzii\_Delta\_2R\_40059143

##### Avrankou\_coluzzii\_Delta\_2R\_50397087

##### Avrankou\_coluzzii\_Delta\_2R\_57936779

##### Avrankou\_coluzzii\_Delta\_2R\_58017231

##### Avrankou\_coluzzii\_Delta\_3R\_40583507

##### Avrankou\_coluzzii\_Delta\_3R\_42095383

##### Avrankou\_coluzzii\_Delta\_3R\_43486612

##### Avrankou\_coluzzii\_Delta\_3R\_43532647

##### Avrankou\_coluzzii\_Delta\_X\_16349957

---

#### Baguida\_*gambiae*\_Delta

Baguida\_gambiae\_Delta\_2R:3411049  
Baguida\_gambiae\_Delta\_2R:3453444  
Baguida\_gambiae\_Delta\_2R:52110329  
Baguida\_gambiae\_Delta\_2R:53504721  
Baguida\_gambiae\_Delta\_3R:50086331  
Baguida\_gambiae\_Delta\_3R:50126025

##### Baguida\_gambiae\_Delta\_2R\_3411049

##### Baguida\_gambiae\_Delta\_2R\_3453444

##### Baguida\_gambiae\_Delta\_2R\_52110329

##### Baguida\_gambiae\_Delta\_2R\_53504721

##### Baguida\_gambiae\_Delta\_3R\_50086331

##### Baguida\_gambiae\_Delta\_3R\_50126025

---

#### Korle-Bu\_*coluzzii*\_Delta

Korle-Bu\_coluzzii\_Delta\_2R:19795384  
Korle-Bu\_coluzzii\_Delta\_2R:19935256  
Korle-Bu\_coluzzii\_Delta\_2R:24507343  
Korle-Bu\_coluzzii\_Delta\_2R:24528127  
Korle-Bu\_coluzzii\_Delta\_2R:24867722  
Korle-Bu\_coluzzii\_Delta\_2R:40748811  
Korle-Bu\_coluzzii\_Delta\_2R:40775991  
Korle-Bu\_coluzzii\_Delta\_2R:49194797  
Korle-Bu\_coluzzii\_Delta\_2R:58948322  
Korle-Bu\_coluzzii\_Delta\_2R:59097397  
Korle-Bu\_coluzzii\_Delta\_2R:59368196  
Korle-Bu\_coluzzii\_Delta\_X:15660088  
Korle-Bu\_coluzzii\_Delta\_X:15703693  
Korle-Bu\_coluzzii\_Delta\_X:15798349  
Korle-Bu\_coluzzii\_Delta\_X:15836204  
Korle-Bu\_coluzzii\_Delta\_X:19194835

##### Korle-Bu\_coluzzii\_Delta\_2R\_19795384

##### Korle-Bu\_coluzzii\_Delta\_2R\_19935256

##### Korle-Bu\_coluzzii\_Delta\_2R\_24507343

##### Korle-Bu\_coluzzii\_Delta\_2R\_24528127

##### Korle-Bu\_coluzzii\_Delta\_2R\_24867722

##### Korle-Bu\_coluzzii\_Delta\_2R\_40748811

##### Korle-Bu\_coluzzii\_Delta\_2R\_40775991

##### Korle-Bu\_coluzzii\_Delta\_2R\_49194797

##### Korle-Bu\_coluzzii\_Delta\_2R\_58948322

##### Korle-Bu\_coluzzii\_Delta\_2R\_59097397

##### Korle-Bu\_coluzzii\_Delta\_2R\_59368196

##### Korle-Bu\_coluzzii\_Delta\_X\_15660088

##### Korle-Bu\_coluzzii\_Delta\_X\_15703693

##### Korle-Bu\_coluzzii\_Delta\_X\_15798349

##### Korle-Bu\_coluzzii\_Delta\_X\_15836204

##### Korle-Bu\_coluzzii\_Delta\_X\_19194835

---

#### Madina\_*gambiae*\_Delta

Madina\_gambiae\_Delta\_2L:14079208  
Madina\_gambiae\_Delta\_2L:14235728  
Madina\_gambiae\_Delta\_2L:14276791  
Madina\_gambiae\_Delta\_2L:15136081  
Madina\_gambiae\_Delta\_2L:15613228  
Madina\_gambiae\_Delta\_2R:24638486  
Madina\_gambiae\_Delta\_2R:28466877  
Madina\_gambiae\_Delta\_3L:10952426  
Madina\_gambiae\_Delta\_3L:10990152  
Madina\_gambiae\_Delta\_3L:13952703  
Madina\_gambiae\_Delta\_3L:14016877  
Madina\_gambiae\_Delta\_3L:14034923  
Madina\_gambiae\_Delta\_X:14317604  
Madina\_gambiae\_Delta\_X:14369513

##### Madina\_gambiae\_Delta\_2L\_14079208

##### Madina\_gambiae\_Delta\_2L\_14235728

##### Madina\_gambiae\_Delta\_2L\_14276791

##### Madina\_gambiae\_Delta\_2L\_15136081

##### Madina\_gambiae\_Delta\_2L\_15613228

##### Madina\_gambiae\_Delta\_2R\_24638486

##### Madina\_gambiae\_Delta\_2R\_28466877

##### Madina\_gambiae\_Delta\_3L\_10952426

##### Madina\_gambiae\_Delta\_3L\_10990152

##### Madina\_gambiae\_Delta\_3L\_13952703

##### Madina\_gambiae\_Delta\_3L\_14016877

##### Madina\_gambiae\_Delta\_3L\_14034923

##### Madina\_gambiae\_Delta\_X\_14317604

##### Madina\_gambiae\_Delta\_X\_14369513

---

#### Obuasi\_*gambiae*\_Delta

Obuasi\_gambiae\_Delta\_2R:28289136  
Obuasi\_gambiae\_Delta\_2R:28425075  
Obuasi\_gambiae\_Delta\_2R:28464858  
Obuasi\_gambiae\_Delta\_2R:28511074  
Obuasi\_gambiae\_Delta\_3L:38536401  
Obuasi\_gambiae\_Delta\_3R:10596787  
Obuasi\_gambiae\_Delta\_3R:10636722  
Obuasi\_gambiae\_Delta\_3R:6860221  
Obuasi\_gambiae\_Delta\_3R:6894288  
Obuasi\_gambiae\_Delta\_3R:6908903  
Obuasi\_gambiae\_Delta\_3R:6944074

##### Obuasi\_gambiae\_Delta\_2R\_28289136

##### Obuasi\_gambiae\_Delta\_2R\_28425075

##### Obuasi\_gambiae\_Delta\_2R\_28464858

##### Obuasi\_gambiae\_Delta\_2R\_28511074

##### Obuasi\_gambiae\_Delta\_3L\_38536401

##### Obuasi\_gambiae\_Delta\_3R\_10596787

##### Obuasi\_gambiae\_Delta\_3R\_10636722

##### Obuasi\_gambiae\_Delta\_3R\_6860221

##### Obuasi\_gambiae\_Delta\_3R\_6894288

##### Obuasi\_gambiae\_Delta\_3R\_6908903

##### Obuasi\_gambiae\_Delta\_3R\_6944074

---

#### Baguida\_*gambiae*\_PM

Baguida\_gambiae\_PM\_X:4481600

##### Baguida\_gambiae\_PM\_X\_4481600

---

#### Korle-Bu\_*coluzzii*\_PM

Korle-Bu\_coluzzii\_PM\_2L:20512784  
Korle-Bu\_coluzzii\_PM\_2L:42166138  
Korle-Bu\_coluzzii\_PM\_2R:20360304  
Korle-Bu\_coluzzii\_PM\_2R:38099416  
Korle-Bu\_coluzzii\_PM\_2R:38131522  
Korle-Bu\_coluzzii\_PM\_2R:54401839  
Korle-Bu\_coluzzii\_PM\_3L:12686869  
Korle-Bu\_coluzzii\_PM\_3R:28478380  
Korle-Bu\_coluzzii\_PM\_3R:28503198  
Korle-Bu\_coluzzii\_PM\_3R:28531750  
Korle-Bu\_coluzzii\_PM\_3R:28562261  
Korle-Bu\_coluzzii\_PM\_3R:28648828

##### Korle-Bu\_coluzzii\_PM\_2L\_20512784

##### Korle-Bu\_coluzzii\_PM\_2L\_42166138

##### Korle-Bu\_coluzzii\_PM\_2R\_20360304

##### Korle-Bu\_coluzzii\_PM\_2R\_38099416

##### Korle-Bu\_coluzzii\_PM\_2R\_38131522

##### Korle-Bu\_coluzzii\_PM\_2R\_54401839

##### Korle-Bu\_coluzzii\_PM\_3L\_12686869

##### Korle-Bu\_coluzzii\_PM\_3R\_28478380

##### Korle-Bu\_coluzzii\_PM\_3R\_28503198

##### Korle-Bu\_coluzzii\_PM\_3R\_28531750

##### Korle-Bu\_coluzzii\_PM\_3R\_28562261

##### Korle-Bu\_coluzzii\_PM\_3R\_28648828

---

#### Madina\_*gambiae*\_PM

Madina\_gambiae\_PM\_2L:13810050  
Madina\_gambiae\_PM\_2L:13835438  
Madina\_gambiae\_PM\_2L:13857697  
Madina\_gambiae\_PM\_2R:2909619  
Madina\_gambiae\_PM\_2R:2924284  
Madina\_gambiae\_PM\_2R:2937977  
Madina\_gambiae\_PM\_2R:2952746  
Madina\_gambiae\_PM\_2R:2966523  
Madina\_gambiae\_PM\_2R:3022442  
Madina\_gambiae\_PM\_2R:3035126  
Madina\_gambiae\_PM\_2R:3048496  
Madina\_gambiae\_PM\_2R:3066659  
Madina\_gambiae\_PM\_2R:3082612  
Madina\_gambiae\_PM\_2R:3098810  
Madina\_gambiae\_PM\_3L:11845057  
Madina\_gambiae\_PM\_3L:8379569  
Madina\_gambiae\_PM\_3R:42366567  
Madina\_gambiae\_PM\_X:10930275  
Madina\_gambiae\_PM\_X:10958263  
Madina\_gambiae\_PM\_X:14496426  
Madina\_gambiae\_PM\_X:14521612

##### Madina\_gambiae\_PM\_2L\_13810050

##### Madina\_gambiae\_PM\_2L\_13835438

##### Madina\_gambiae\_PM\_2L\_13857697

##### Madina\_gambiae\_PM\_2R\_2909619

##### Madina\_gambiae\_PM\_2R\_2924284

##### Madina\_gambiae\_PM\_2R\_2937977

##### Madina\_gambiae\_PM\_2R\_2952746

##### Madina\_gambiae\_PM\_2R\_2966523

##### Madina\_gambiae\_PM\_2R\_3022442

##### Madina\_gambiae\_PM\_2R\_3035126

##### Madina\_gambiae\_PM\_2R\_3048496

##### Madina\_gambiae\_PM\_2R\_3066659

##### Madina\_gambiae\_PM\_2R\_3082612

##### Madina\_gambiae\_PM\_2R\_3098810

##### Madina\_gambiae\_PM\_3L\_11845057

##### Madina\_gambiae\_PM\_3L\_8379569

##### Madina\_gambiae\_PM\_3R\_42366567

##### Madina\_gambiae\_PM\_X\_10930275

##### Madina\_gambiae\_PM\_X\_10958263

##### Madina\_gambiae\_PM\_X\_14496426

##### Madina\_gambiae\_PM\_X\_14521612

---

#### Obuasi\_*gambiae*\_PM

Obuasi\_gambiae\_PM\_2R:3018207  
Obuasi\_gambiae\_PM\_3L:39831247  
Obuasi\_gambiae\_PM\_3L:39848513  
Obuasi\_gambiae\_PM\_3R:48480402  
Obuasi\_gambiae\_PM\_3R:48535299  
Obuasi\_gambiae\_PM\_X:14032886  
Obuasi\_gambiae\_PM\_X:14078096  
Obuasi\_gambiae\_PM\_X:14375402  
Obuasi\_gambiae\_PM\_X:14696755  
Obuasi\_gambiae\_PM\_X:14729619

##### Obuasi\_gambiae\_PM\_2R\_3018207

##### Obuasi\_gambiae\_PM\_3L\_39831247

##### Obuasi\_gambiae\_PM\_3L\_39848513

##### Obuasi\_gambiae\_PM\_3R\_48480402

##### Obuasi\_gambiae\_PM\_3R\_48535299

##### Obuasi\_gambiae\_PM\_X\_14032886

##### Obuasi\_gambiae\_PM\_X\_14078096

##### Obuasi\_gambiae\_PM\_X\_14375402

##### Obuasi\_gambiae\_PM\_X\_14696755

##### Obuasi\_gambiae\_PM\_X\_14729619

---
