## Supplementary Data S6 for "Genome-wide association studies reveal novel loci associated with pyrethroid and organophosphate resistance in *Anopheles gambiae* s.l.": Supplementary_Data_S6.html

gwas\_summary


### GAARD GWAS results summary

Each plot shows the 1000 most significant SNPs in the GWAS for that
sample set. At the top, the P value for each SNP is shown, colour-coded
by chromosome. Underneath is a heatmap showing the pairwise correlation
between all 1000 SNPs, helping to see groups of SNPs that are in linkage
disequilibrium. Each point in the *P*-value plot is lined up with
its column in the heatmap. Beneath the column, lines heatmap columns to
a map of the genome to show the genomic position of each SNP. Black
horizontal bars between the heatmap and the linking lines indicate
genomic regions containing a high density of these top 1000 SNPs (100 kb
windows that contained at least 10 SNPs).

---

Avrankou\_*coluzzii*\_Delta  
Baguida\_*gambiae*\_Delta  
Korle-Bu\_*coluzzii*\_Delta  
Madina\_*gambiae*\_Delta  
Obuasi\_*gambiae*\_Delta  
Baguida\_*gambiae*\_PM  
Korle-Bu\_*coluzzii*\_PM  
Madina\_*gambiae*\_PM  
Obuasi\_*gambiae*\_PM

---

#### Avrankou\_*coluzzii*\_Delta

---

#### Baguida\_*gambiae*\_Delta

---

#### Korle-Bu\_*coluzzii*\_Delta

---

#### Madina\_*gambiae*\_Delta

---

#### Obuasi\_*gambiae*\_Delta

---

#### Baguida\_*gambiae*\_PM

  \_\_\_

#### Korle-Bu\_*coluzzii*\_PM

---

#### Madina\_*gambiae*\_PM

---

#### Obuasi\_*gambiae*\_PM

---
